# Hybridized distance- and contact-based hierarchical structure modeling for folding soluble and membrane proteins

**DOI:** 10.1101/2020.07.05.188466

**Authors:** Rahmatullah Roche, Sutanu Bhattacharya, Debswapna Bhattacharya

## Abstract

Crystallography and NMR system (CNS) is currently the *de facto* standard for fragment-free *ab initio* protein folding from inter-residue distance or contact maps. Despite its widespread use in protein structure prediction, CNS is a decade-old macromolecular structure determination system that was originally developed for solving macromolecular geometry from experimental restraints as opposed to predictive modeling driven by interaction map data. As such, the adaptation of the CNS experimental structure determination protocol for *ab initio* protein folding is intrinsically anomalous that may undermine the folding accuracy of computational protein structure prediction. In this paper, we propose a new CNS-free hierarchical structure modeling method called DConStruct for folding both soluble and membrane proteins driven by distance and contact information. Rigorous experimental validation shows that DConStruct attains much better reconstruction accuracy than CNS when tested with the same input contact map at varying contact thresholds. The hierarchical modeling with iterative self-correction employed in DConStruct scales at a much higher degree of folding accuracy than CNS with the increase in contact thresholds, ultimately approaching near-optimal reconstruction accuracy at higher-thresholded contact maps. The folding accuracy of DConStruct can be further improved by exploiting distance-based hybrid interaction maps at tri-level thresholding, as demonstrated by the better performance of our method in folding difficult free modeling targets from the 12th and 13th rounds of the Critical Assessment of techniques for protein Structure Prediction (CASP) experiments compared to several popular CNS- and fragment-based approaches, some of which even using much finer-grained distance maps than ours. Additional large-scale benchmarking shows that DConStruct can significantly improve the folding accuracy of membrane proteins compared to a CNS-based approach. These results collectively demonstrate the feasibility of greatly improving the accuracy of *ab initio* protein folding by optimally exploiting the information encoded in inter-residue interaction maps beyond what is possible by CNS.

**Author summary:** Predicting the folded and functional 3-dimensional structure of a protein molecule from its amino acid sequence is of central importance to structural biology. Recently, promising advances have been made in *ab initio* protein folding due to the reasonably accurate estimation of inter-residue interaction maps at increasingly higher resolutions that range from binary contacts to finer-grained distances. Despite the progress in predicting the interaction maps, approaches for turning the residue-residue interactions projected in these maps into their precise spatial positioning heavily rely on a decade-old experimental structure determination protocol that is not suitable for predictive modeling. This paper presents a new hierarchical structure modeling method, DConStruct, which can better exploit the information encoded in the interaction maps at multiple granularities, from binary contact maps to distance-based hybrid maps at tri-level thresholding, for improved *ab initio* folding. Multiple large-scale benchmarking experiments show that our proposed method can substantially improve the folding accuracy for both soluble and membrane proteins compared to state-of-the-art approaches. DConStruct is licensed under the GNU General Public License v3 and freely available at https://github.com/Bhattacharya-Lab/DConStruct.

## Introduction

The development of a computational method that can successfully predict the functional 3-dimensional (3D) structure of a protein molecule purely from its amino acid sequence is of central importance to structural biology [1]. In the recent past, promising progress has been made in this endeavor mediated by reasonably accurate prediction of inter-residue distance or contact maps using sequence co-evolution coupled with deep learning [2–7], and performing data-assisted folding driven by such predicted interaction maps [8–10]. Inter-residue interaction maps contain spatial proximity information that can be translated into geometric constraints to directly construct protein 3D models by maximal constraint satisfaction. Therefore, the prediction of inter-residue distance or contact interaction maps and predicted interaction-assisted 3D structure modeling has fueled considerable research efforts in the community [11, 12].

Despite the rapid advances in predicting interaction maps by utilizing state-of-the-art deep learning architectures [4], progress in building 3D models from the predicted maps has been disproportionately slow. ROSETTA molecular modeling suite [13] offers various functions to integrate predicted interaction maps into its internal scoring function as additional constraints for fragment assembly-based *ab initio* folding [14] that can be coupled with loop perturbation sampling [15], but fragment-based folding involves extensive conformational sampling requiring a large amount of computing power. Beyond the realm of time-consuming conformational sampling with fragments, majority of fragment-free distance or contact-based protein folding methods [16–19] rely on a decade-old experimental protein structure determination software called Crystallography and NMR system (CNS) [20], which was originally developed for solving macromolecular geometry from experimental nuclear overhauser enhancement (NOE) restraints as opposed to predicted data and therefore intrinsically incompatible for data-driven predictive modeling. Even the most recent advances in protein structure prediction [8, 9] are primarily due to the progress made in predicting finer-grained interaction maps, but 3D model building from the predicted fine-grained maps still utilize CNS-based experimental structure determination protocol. Because of the dependency on CNS, folding accuracy of computational protein structure modeling methods may get compromised, hindering the realization of their full potential. Thus, there is a critical need to develop a fragment-free folding protocol specifically suitable for predicted inter-residue interaction map data rather than relying on the CNS-based structure determination approach.

CNS-based macromolecular structure determination protocol (CNSsolve) follows several conventions that can be revised for improved folding from predicted inter-residue interaction maps. First, CNS adopts a molecular topology file (.mtf) format for representing polypeptide geometry in all-atom representation, while distance or contact maps are usually defined at a coarse-grained level (e.g., between C_β_–C_β_ or C_α_–C_α_ atom pairs). As such, adaptation of a detailed all-atom representation may not be necessary for distance or contact-assisted folding, at least during the early stages, to accurately predict the backbone geometry while the side-chain atoms can be added conditioned on the backbone conformation subsequently. Adopting a coarse-grained representation reduces the conformational space that may improve folding accuracy and efficiency. Second, CNS has an in-built biophysical force field with bonded and non-bonded terms, some of which may be conflicting or mutually contradictory with the predicted inter-residue interactions, posing difficulties in maximal satisfaction of interaction restraints. Finally, CNS only accepts restraints in a specific format that cannot be easily customized or extended for different applications.

In this article, we present a new inter-residue interaction-assisted hierarchical folding method based on multistage structure modeling with iterative self-correction. Free from the limitations of CNS, our folding method employs 3-stage hierarchical predictive modeling with iterative self-correction driven purely by the geometric restraints induced by inter-residue interactions and secondary structures. Starting from a residue-residue interaction map and secondary structure, our method (DConStruct) can hierarchically estimate the correct overall fold of a target protein in coarse-grained mode to progressively optimize local and non-local interactions while enhancing the secondary structure topology in a self-correcting manner. DConStruct is versatile in that it can exploit the information encoded in the interaction maps at multiple granularities ranging from binary contact maps to distance-based maps at varying thresholds.

We rigorously test DConStruct on several hundred soluble and membrane proteins as well as public data from the latest rounds of the Critical Assessment of techniques for protein Structure Prediction (CASP) experiments. Our experimental results show that DConStruct yields much better folding accuracy than existing CNS-based folding protocols when tested with the same input. Our method attains better performance in folding difficult CASP free modeling targets compared to both CNS- and fragment-based approaches, some of which even using much finer-grained interaction maps than ours. The open-source DConStruct software package, licensed under the GNU General Public License v3, is freely available at https://github.com/Bhattacharya-Lab/DConStruct.

## Results

### DConStruct: hybridized distance- and contact-based hierarchical structure modeling

Fig 1 illustrates the DConStruct hierarchical structure modeling protocol. Different from the CNS-based approaches [8,9,16–19], DConStruct employs multiscale predictive modeling with iterative self-correction comprising of three modeling stages. The initial modeling stage employs coarse-grained modeling considering the protein conformation as a string of beads, in which each bead corresponds to the C_α_ atom of an amino acid residue, in order to estimate the overall fold from a sparse set of interatomic interactions and secondary structure information. Given an input interaction map and secondary structure for an amino acid sequence, approximate spatial positioning between the residues is first estimated using prior knowledge of a protein’s backbone geometry, derived from the pseudo-covalent bonds formed between the C_α_ atoms and secondary structure-specific local preferences of the inter-residue distances [21], thus generating a sparse proximity map. For sequentially distant residue pairs, graph-theoretic formulation is adopted to fill the missing entries in the proximity map, which is further refined using the idealized geometry of the secondary structure elements (SSEs) to enhance physical realism. Multidimensional Scaling (MDS) [22–24] is then employed to estimate the 3D coordinates from the proximity map, resulting in a pool of coarse-grained models. The second stage of DConStruct employs iterative self-correction of the coarse-grained model pool using corrective coordinate perturbation heuristics followed by Limited-memory Broyden–Fletcher–Goldfarb–Shanno (LBFGS) [25] optimization. Next, top models are selected based on maximal satisfaction of the geometric restraints induced by the interaction map coupled with a secondary structure-assisted geometric chirality checking [26]. The final modeling stage of DConStruct consists of atomic-level iterative self-correction. First, MODELLER [27] is used to generate all-atom models from the selected top models. Then, unsatisfied high-confidence interactions, non-interactions, and secondary structure restraints are identified and cumulatively applied with iterative self-correction and model combination to generate the final folded conformation.

**Fig 1.**
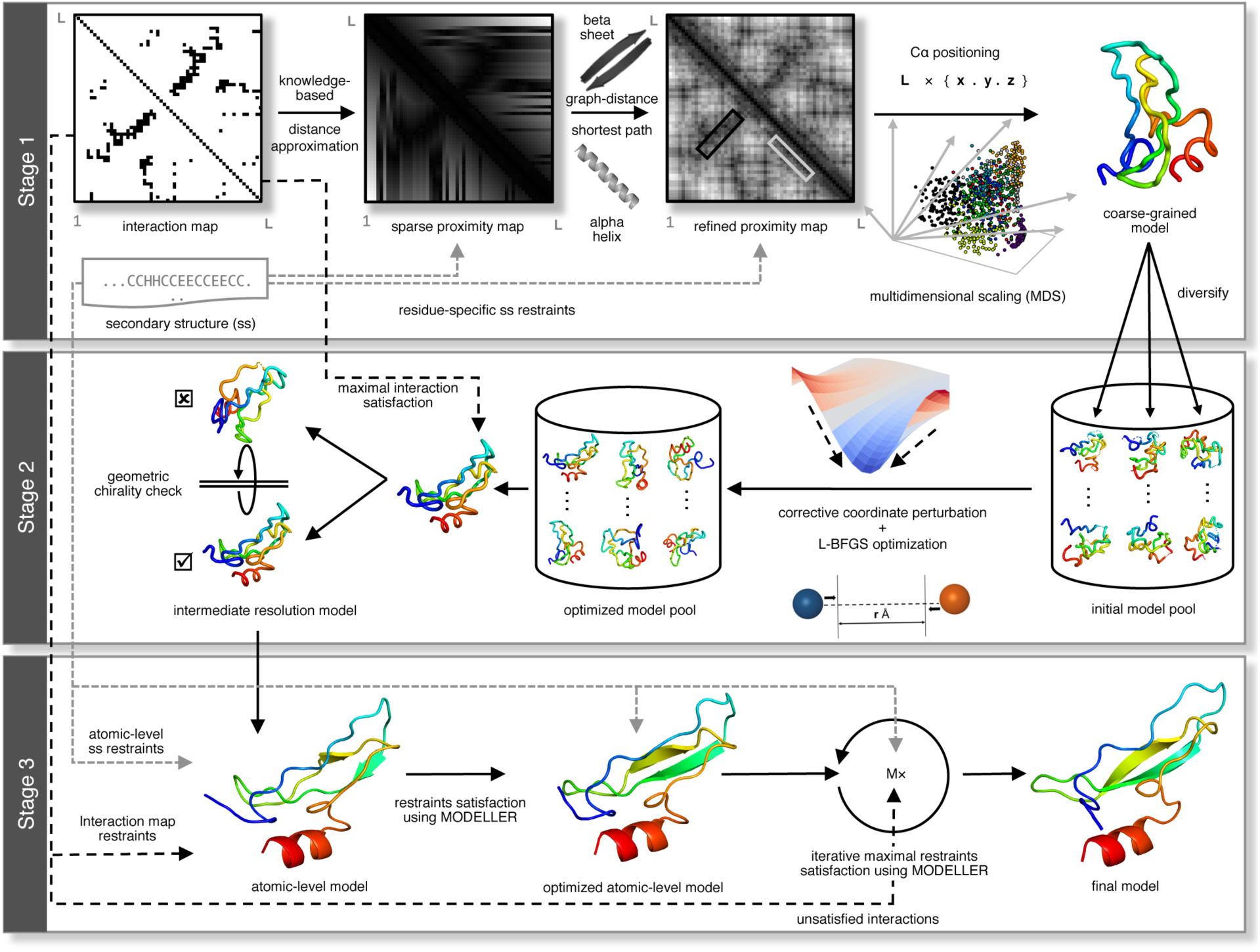
The 3-stage hierarchical structure modeling protocol of DConStruct.

Our test data include 150 single domain soluble proteins from the FRAGFOLD dataset [28], 40 Free Modeling (FM) target domains with publicly available experimental structures from the 12th and 13th editions of CASP [29, 30], 510 membrane proteins [31, 32], and 15 targets from the EVfold set [16]. To evaluate the reconstruction of 3D structural models, we use true residue-residue interaction maps and true secondary structures. To assess *ab initio* folding performance, we use DMPfold [8] distance predictor and SPIDER3 [33] secondary structure predictor, both employing cutting-edge deep learning architectures. We install and run DMPfold locally to predict distance histogram maps directly from the multiple sequence alignments (MSA) [34] without using iterative refinement (i.e., rawdistpred.current files) containing 20 distance bins with associated likelihoods between interacting residue pairs. We also predict secondary structures by locally installing and running SPIDER3 with default parameter settings. We compare our new method DConStruct against a pure contact-driven approach FT-COMAR [35], CNS-based contact- and secondary structure-driven CONFOLD-like protocols [17,18,31], fragment-based contact- and secondary structure-driven pipelines employing ROSETTA [14, 15], and CNS-based distance- and secondary structure-driven method DMPfold [8]. For CNS- and ROSETTA-based contact-assisted *ab initio* folding methods [14, 18], RaptorX [4] contact maps, obtained by submitting jobs to its web server, are used. CGLFold results are taken directly from its published paper. All other methods are run locally with parameters set according to their published papers.

### Reconstruction of soluble proteins

We evaluate the reconstruction performance of soluble proteins using the true three-state secondary structures computed from the experimental structures by DSSP [36] and the true C_α_–C_α_ and C_β_–C_β_ contact maps calculated at kÅ threshold (k = 8, 8.5, 9, 9.5, 10, 10.5, 11, 11.5, 12). We define a contact when the Euclidian distance between two representative atoms (C_α_–C_α_ or C_β_–C_β_) for a residue pair is at most kÅ with a minimum sequence separation of 6 residues. We compare our new method DConStruct with the standalone contact-based method FT-COMAR [35] and the CNS-based CONFOLD protocol [17] on the 150 FRAGFOLD soluble protein domains with length ranging from 50 to 266 residues. FT-COMAR is a pure distance geometry-based structure reconstruction method that can take only C_α_–C_α_ contacts but no secondary structure, whereas CONFOLD is a CNS-based state-of-the-art contact-assisted method that utilizes contact and secondary structure for structure reconstruction. All methods are tested with the same input. The reconstruction accuracy is evaluated for the top predicted models using TM-score [37], a widely used metric for evaluating the quality of 3D structural models.

As shown in **Tables 1–2**, DConStruct significantly outperforms the tested methods FT-COMAR and CONFOLD on the 150 FRAGFOLD soluble protein domains across all thresholds for both C_α_–C_α_ (**Table 1**) and C_β_–C_β_ (**Table 2**) contact maps. For the C_α_–C_α_ contact maps at the standard threshold of 8Å, the mean TM-score of the top models generated by DConStruct is 0.85, which exceeds that of CONFOLD and FT-COMAR by 0.04 and 0.36 TM-score points, respectively, and statistically significantly better compared to both CONFOLD (*p*-value 1.49e-13) and FT-COMAR (*p*-value 2.3e-46) at 95% confidence level. Moreover, DConStruct attains a better TM-score than FT-COMAR and CONFOLD for ∼99% and ∼89% targets, respectively (**S1 Table**). It is interesting to note that for the C_α_–C_α_ maps at higher contact thresholds beyond 8Å, in addition to outperforming FT-COMAR by a very large margin, the reconstruction accuracy of DConStruct becomes progressively better compared to CONFOLD as contact threshold increases with DConStruct attaining ∼7% (mean TM-scores of 0.90 vs. 0.84), ∼9% (mean TM-scores of 0.94 vs. 0.86), and ∼9% (mean TM-scores of 0.95 vs. 0.87) better performance compared to CONFOLD at 9, 10, and 11Å contact thresholds, respectively. Between 11Å and 12Å, performance improvement stays at ∼9% level with DConStruct attaining a mean TM-score ∼0.95, which is much higher than that of CONFOLD (mean TM-score ≤ 0.88). Fig 2A shows the TM-score distributions of the reconstructed models at various contact thresholds. For all thresholds, the DConStruct distributions are skewed toward higher TM-score regions, indicating better reconstruction performance compared to FT-COMAR and CONFOLD that gets progressively better at higher thresholds. With the increase in contact thresholds, DConStruct results in much more higher number of near-optimal reconstruction cases having TM-score → 1.0. For example, at 8 Å threshold, only 3 out of 150 reconstructed models using DConStruct have TM-score > 0.95, while CONFOLD fails to reconstruct any structure with TM-score > 0.95. At 12Å threshold, 105 out of 150 (70%) reconstructed models using DConStruct have TM-score > 0.95, whereas CONFOLD attains TM-score > 0.95 only for 11 out of 150 (∼7%) cases. Similar trends are observed for the reconstruction with C_β_–C_β_ contact maps, for which DConStruct continues to significantly outperform CONFOLD across all contact thresholds. For the C_β_–C_β_ contact maps at 8Å threshold, DConStruct is statistically significantly better than CONFOLD (*p*-value 2.07e-06) and attains better TM-score than CONFOLD for ∼73% targets (**S2 Table**). As shown in Fig 2B, for C_β_–C_β_ contacts at higher-thresholded contact maps beyond 8Å, the reconstruction performance of DConStruct gets progressively better compared to CONFOLD with the DConStruct distributions being skewed more and more towards higher TM-score regions. Collectively, the results demonstrate that DConStruct attains much better reconstruction accuracy compared to the CNS-based CONFOLD protocol across multiple contact thresholds for both C_α_–C_α_ and C_β_–C_β_ contacts in addition to greatly outperforming the pure distance geometry-based FT-COMAR method. Notably, the reconstruction performance of DConStruct becomes markedly better than CONFOLD for higher-thresholded contact maps beyond the standard contact threshold of 8Å currently used by the community. Additional controlled experiments reveal that the better performance of DConStruct at higher-thresholded contact maps is due to the cooperativity between various stages of its hierarchical modeling paradigm, as discussed later.

**Fig 2.**
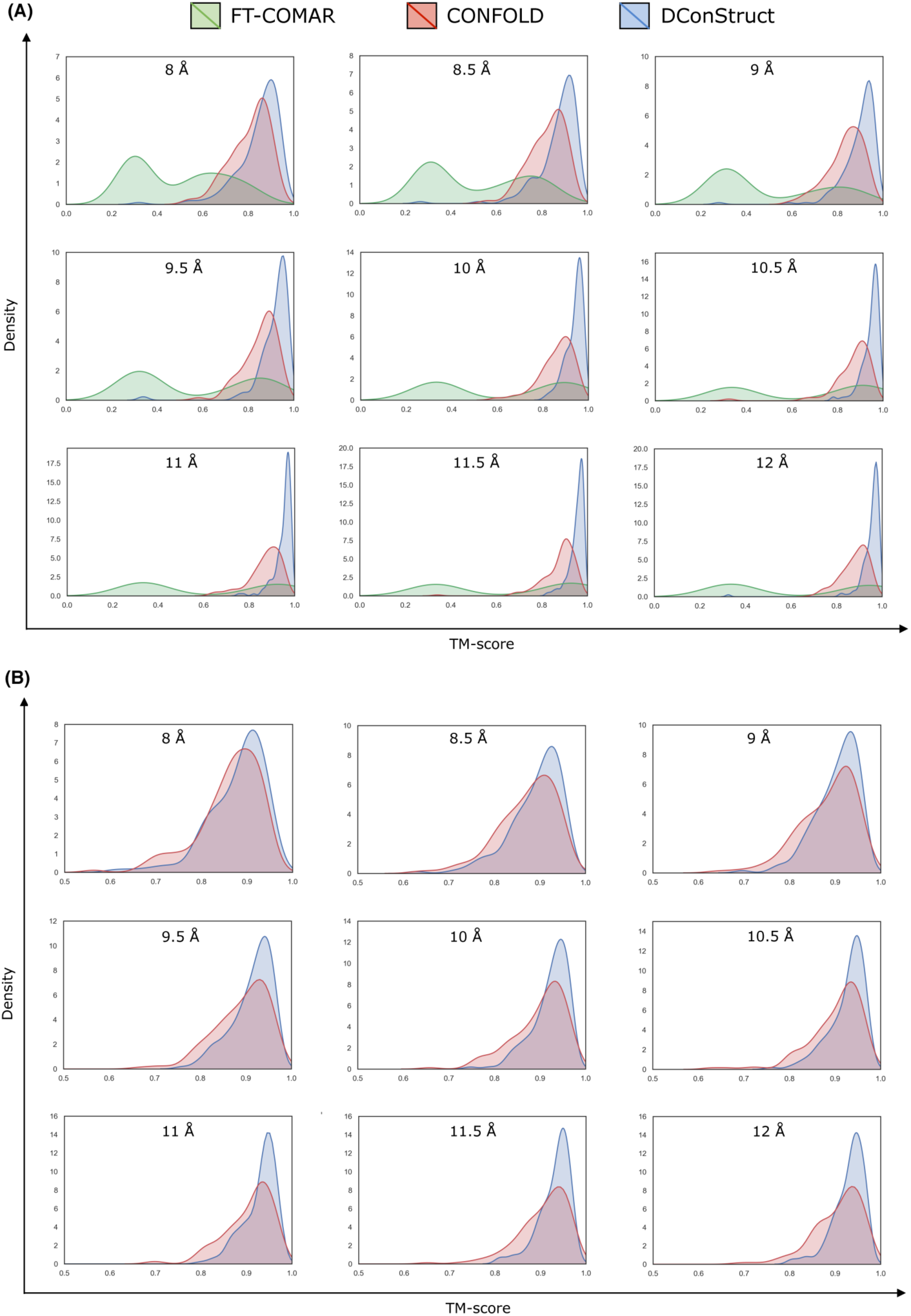
TM-score distributions of the reconstructed models on 150 FRAGFOLD soluble protein domains using (A) FT-COMAR, CONFOLD, and DConStruct for true C_α_–C_α_ contact maps; and (B) CONFOLD and DConStruct for true C_β_–C_β_ contact maps.

**Table 1.** Reconstruction performance of soluble proteins on 150 FRAGFOLD domains for true C_α_-C_α_ contact maps. The mean TM-score of top predicted models are reported. Values in bold represent the best performance.

| Method | 8Å | 8.5Å | 9Å | 9.5Å | 10Å | 10.5Å | 11Å | 11.5Å | 12Å |
| --- | --- | --- | --- | --- | --- | --- | --- | --- | --- |
| DConStruct | **0.85** | **0.87** | **0.9** | **0.91** | **0.94** | **0.95** | **0.95** | **0.95** | **0.95** |
| CONFOLD | 0.81 | 0.82 | 0.84 | 0.85 | 0.86 | 0.87 | 0.87 | 0.88 | 0.88<sup>a</sup> |
| FT-COMAR | 0.49 | 0.51 | 0.51 | 0.57 | 0.62 | 0.65 | 0.62 | 0.65 | 0.63 |

<sup>a</sup> CONFOLD fails to run for the target 1dixA at 12Å, and the mean TM-score for CONFOLD is measured over the rest 149 targets at 12Å.

**Table 2.** Reconstruction performance of soluble proteins on 150 FRAGFOLD domains for true C_β_-C_β_ contact maps. The mean TM-score of top predicted models are reported. Values in bold represent the best performance.

| Method | 8Å | 8.5Å | 9Å | 9.5Å | 10Å | 10.5Å | 11Å | 11.5Å | 12Å |
| --- | --- | --- | --- | --- | --- | --- | --- | --- | --- |
| DConStruct | **0.88** | **0.89** | **0.90** | **0.91** | **0.92** | **0.93** | **0.93** | **0.93** | **0.93** |
| CONFOLD | 0.86 | 0.87<sup>a</sup> | 0.88 | 0.89 | 0.9 | 0.90<sup>b</sup> | 0.91 | 0.9 | 0.9 |

<sup>a, b</sup> CONFOLD fails to run for target 1smxA at 8.5Å, and for targets 1fcyA, 1ql0A using 10.5Å, and the mean TM-score for CONFOLD is measured over the rest 149 targets at 8.5Å, and over the rest 148 targets at 10.5Å.

Two representative examples shown in Fig 3 illustrate the advantage of DConStruct over CNS-based CONFOLD, especially at higher-thresholded contact maps. The first (Fig 3A) is a c2 domain from novel protein kinase C epsilon from Rat (PDB ID: 1gmi), a mainly β protein of 135 residues. For this target, the TM-scores of the reconstructed models using DConStruct and CONFOLD at 8Å threshold are 0.79 and 0.74, respectively; whereas at 10Å and 12Å thresholds, DConStruct reconstructs much higher quality models with the TM-scores of 0.9 and 0.96, respectively, which are substantially better than CONFOLD having a TM-score of 0.82 at both 10Å and 12Å thresholds. Of note, DConStruct is able to reach sub-angstrom reconstruction accuracy at 12Å threshold for this target with a C_α_ root mean squared deviation (rmsd) of 0.96Å, outperforming CONFOLD by a large margin. The second (Fig 3B) is a phosphotransferase IIa-mannitol protein from Escherichia coli (PDB ID: 1a3a). Reconstruction for this α+β target of 145 residues at 8Å threshold results in a TM-score of 0.93 for DConStruct and 0.84 for CONFOLD. At 12Å distance threshold, reconstruction with DConStruct results in a TM-score of 0.97, whereas CONFOLD attains a TM-score of 0.88. Once again, DConStruct reaches sub-angstrom reconstruction accuracy at 12Å threshold with a C_α_ rmsd of 0.82Å that is substantially better than CONFOLD.

**Fig 3.**
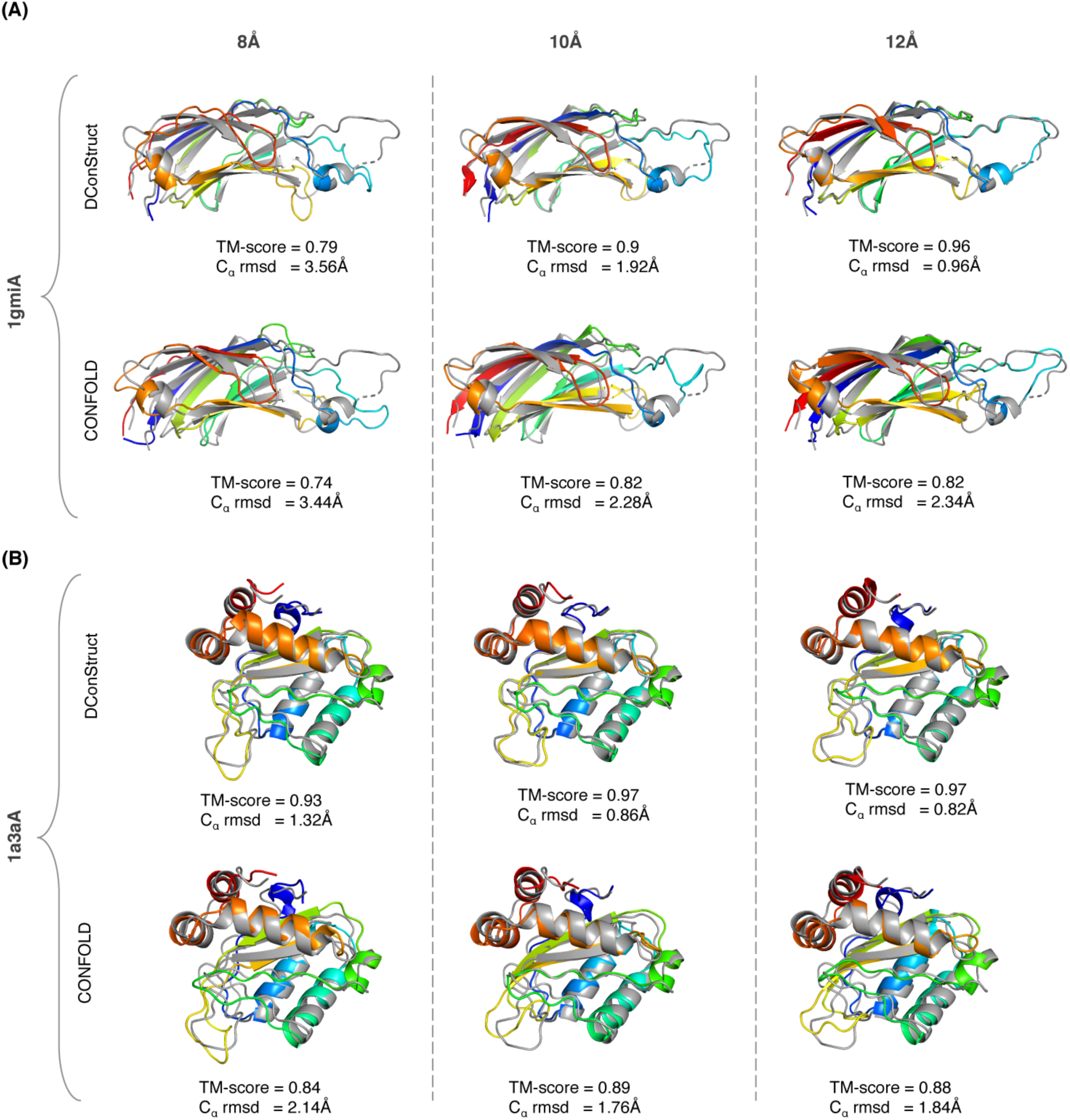
Superimpositions between the reconstructed models (rainbow) and the corresponding experimental structures (gray) for two soluble proteins (A) PDB ID 1gmi and chain A; and (B) PDB ID 1a3a and chain A; using DConStruct and CONFOLD for true C_α_-C_α_ contact maps at 8, 10, and 12Å thresholds.

The results not only demonstrate the advantage of DConStruct over CNS-based CONFOLD, but open up some important follow-up questions. First, recognizing that DConStruct scales at a much better degree of reconstruction accuracy than CONFOLD with the increase in contact thresholds, a natural question to ask is can we improve the performance even further by combining multiple contact thresholds into some form of hybrid interaction maps? Second, does the better reconstruction performance of DConStruct translate to better *ab initio* protein folding? Finally, can we use DConStruct to improve the folding accuracy of membrane proteins? We systematically examine these questions by performing rigorous experiments.

To examine whether it is possible to further improve the reconstruction performance by combining multiple contact thresholds, we formulate hybrid interaction maps at tri-level thresholding. Instead of using a single contact threshold, our hybrid interaction maps use tri-level thresholding with variable upper bounds of 8, 10, and 12Å derived from the Euclidian distance between two representative atoms (C_α_–C_α_ or C_β_– C_β_) for a residue pair having a minimum sequence separation of 6 residues. That is, we interpolate the real-valued Euclidian distance between a residue pair to one of the three upper bounds, thus resulting in tri-level thresholding. We reconstruct the same set of 150 soluble proteins after feeding the hybrid interaction maps and three-state secondary structures into DConStruct, and compute the TM-scores of the reconstructed models.

For the C_α_–C_α_ maps, running DConStruct with the hybrid interaction maps results in near-optimal reconstruction with a mean TM-score of 0.97, which not only outperforms the mean TM-score of reconstruction with binary contact maps across all thresholds but also yields better TM-scores for 149 out of 150 (∼99%) cases than 8 and 10Å thresholds and 144 out of 150 (∼96%) cases than 12Å threshold (**S3 Table**). For the C_β_–C_β_ maps, reconstruction with the hybrid interaction maps leads to better TM-scores than reconstruction with binary contact maps for 145 out of 150 (∼97%), 99 out of 150 (∼66%), and 87 out of 150 (∼58%) cases for 8, 10, and 12Å thresholds, respectively (**S4 Table**). That is, DConStruct with hybrid interaction maps at tri-level thresholding yields better reconstruction performance than contact maps at a fixed threshold. As shown in Fig 4, running DConStruct with hybrid maps further improves the reconstruction accuracy for the two representative proteins described in Fig 3 with the first protein (PDB ID: 1gmi) attaining a TM-score of 0.97 and the second protein (PDB ID: 1a3a) achieving a TM-score of 0.98, both reaching improved sub-angstrom C_α_ rmsds of 0.78Å and 0.62Å for 1gmi and 1a3a, respectively.

**Fig 4.**
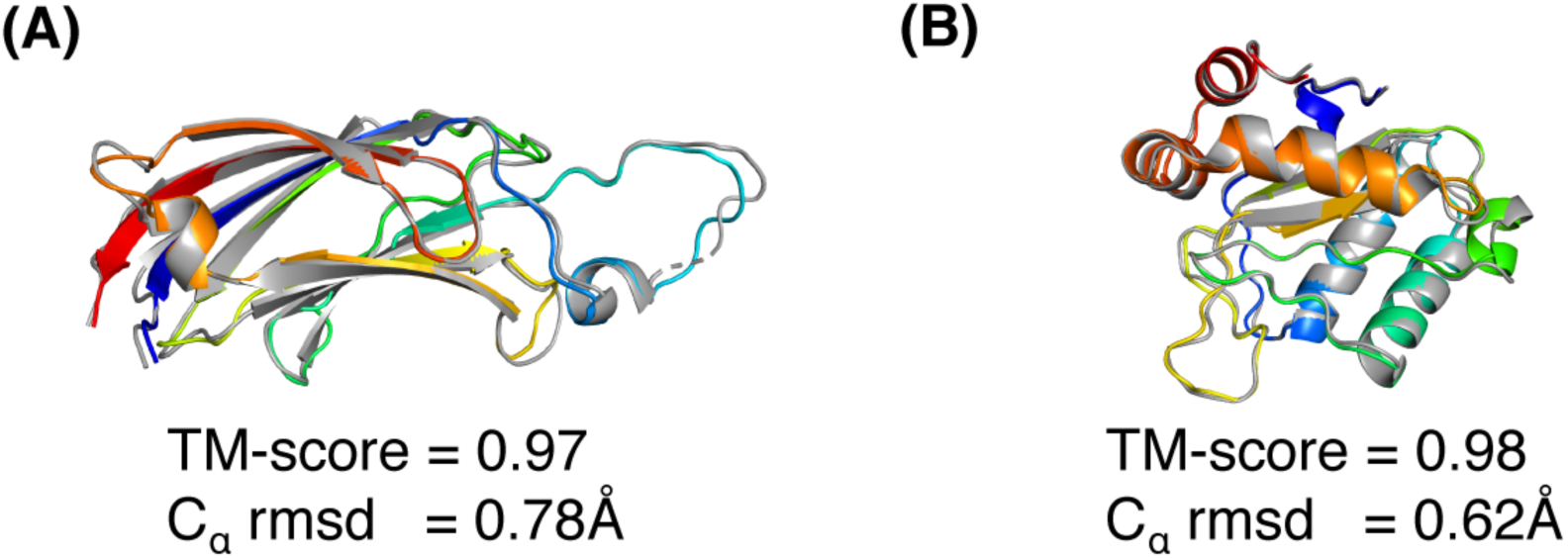
Superimpositions between the reconstruction models (rainbow) and its native structure (gray) for two soluble proteins (A) PDB ID 1gmi and chain A; and (B) PDB ID 1a3a and chain A; generated by DConStruct with hybrid interaction maps at tri-level thresholding.

### Folding CASP free modeling targets

To investigate whether the improved reconstruction performance of DConStruct translates to better *ab initio* folding, we perform predictive modeling for 40 free modeling (FM) targets with publicly available experimental structures from the 12th and 13th rounds of the Critical Assessment of techniques for protein Structure Prediction (CASP) experiments. We compare the predictive modeling performance of DConStruct with two CNS-based approaches: DMPfold [8] and CONFOLD2 [18], as well as two fragment-based methods: ROSETTA [13] as used in the PConsFold protocol [14] and CGLfold [15]. DMPfold [8] is a cutting-edge *ab initio* folding method that employs deep learning to predict inter-atomic distance bounds, torsion angles, and hydrogen bonds and feeds these constraints into CNS to build models in an iterative fashion. CONFOLD2 [18] employs CNS to integrate predicted contacts and secondary structure in a two-stage modeling pipeline in which unsatisfied contacts are filtered out after initial model generation. Popular fragment-based method ROSETTA [13] adds constraints from predicted contacts into the well-established fragment assembly engine as implemented in the PconsFold [14] protocol. CGLfold [15] is a recent fragment-based method that combines global exploration and loop perturbation using predicted contacts. We run DMPfold with default parameter settings to predict structural models using its CNS-based iterative modeling. CONFOLD2 only accepts contact maps at 8Å threshold and PconsFold protocol utilizes ROSETTA’s FADE function for contact constraints with its parameters set for 8Å contacts. We, therefore, perform *ab initio* folding using CONFOLD2 and ROSETTA by feeding 8Å contact maps predicted from the state-of-the-art RaptorX contact prediction method [4] together with secondary structures predicted using SPIDER3 [33]. For CGLFold, we collect results for the 29 CASP FM targets from their published paper [15]. To perform predictive modeling using DConStruct driven by hybrid interaction maps at tri-level thresholding, we collect the DMPfold predicted initial distance histograms (rawdispred.current files) containing 20 distance bins with associated likelihoods and convert them into hybrid interaction maps with variable upper bounds of 8, 10, and 12Å by summing up the likelihoods for distance bins below the three distance thresholds of 8, 10, and 12Å and subsequently select the top contacts based on their likelihoods, resulting in predicted hybrid interaction maps at tri-level thresholding. In addition to predicted hybrid interaction maps, we also feed three-state secondary structures predicted using SPIDER3 [33] into DConStruct. Unlike DMPfold, we do not use any other predicted structural features such as torsion angles and hydrogen bonds or perform any CNS-based iterative modeling in DConStruct. To evaluate *ab initio* folding performance, we compare TM-scores of the top predicted models from each of the tested methods. Additionally, we evaluate the number of models with correct overall folds having TM-score > 0.5 [38].

As reported in **Table 3**, DConStruct outperforms all other methods by attaining better TM-scores and correctly folding more FM targets. DConStruct attains the highest mean TM-score of 0.46, which is statistically significantly better than the second-best performing method DMPfold (mean TM-score of 0.42) at 95% confidence level (*p*-value of 0.03329857). The median TM-score of DConStruct is 0.5, which is also the highest and significantly better than all competing methods, including the CNS-based protocols DMPfold and CONFOLD2 as well as the fragment-based approaches ROSETTA and CGLFold (**S5 Table**). DConStruct correctly folds 20 out of 40 CASP FM targets, whereas the number of correct folds for DMPfold, CONFOLD2, CGLFold, and ROSETTA are only 15, 10, 8, and 6, respectively. The results demonstrate that DConStruct delivers much better *ab initio* folding performance compared to the CNS- and fragment-based approaches, including the state-of-the-art DMPfold protocol employing CNS-based iterative modeling using much finer-grained distance maps than ours together with additional predicted structural features such as torsion angles and hydrogen bonds. Of note, the hybrid interaction maps used in DConStruct are derived from the DMPfold predicted initial distance histograms. That is, even with lower-resolution interaction maps and much less information, DConStruct leads to better *ab initio* folding accuracy than CNS, underscoring its effectiveness in predictive modeling with inter-residue interaction maps beyond what is possible by CNS.

**Table 3.** Folding performance of CASP free modeling protein targets on 40 CASP12 and CASP13 free modeling target domains. Values in bold represent the best performance.

| Method | Mean TM-score | Median TM-score | # TM-scores > 0.5 |
| --- | --- | --- | --- |
| DConStruct | **0.46** | **0.50** | **20/40** |
| DMPfold | 0.42 | 0.38 | 15/40 |
| CGLFold* | 0.40 | 0.43 | 8/29 |
| CONFOLD2 | 0.38 | 0.32 | 10/40 |
| ROSETTA | 0.37 | 0.36 | 6/40 |

\* CGLFold results are adopted directly from the published results containing 29 CASP free modeling target domains.

Fig 5 shows the *ab initio* models predicted by DMPfold and DConStruct for four representative CASP FM targets. For the two CASP12 FM targets T0898-D1 and T0870-D1, DMPfold fails to attain the correct overall fold whereas DConStruct correctly folds both targets attaining TM-scores of 0.65 and 0.67 for T0898-D1 and T0870-D1, respectively. T0968s2-D1 and T0957s2-D1 are two CASP13 FM targets, for both of which DConStruct predicts more accurate models with TM-score ≥ 0.7, much better than DMPfold. In summary, the advantage of DConStruct over the state-of-the-art CNS-based *ab initio* folding method DMPfold is significant.

**Fig 5.**
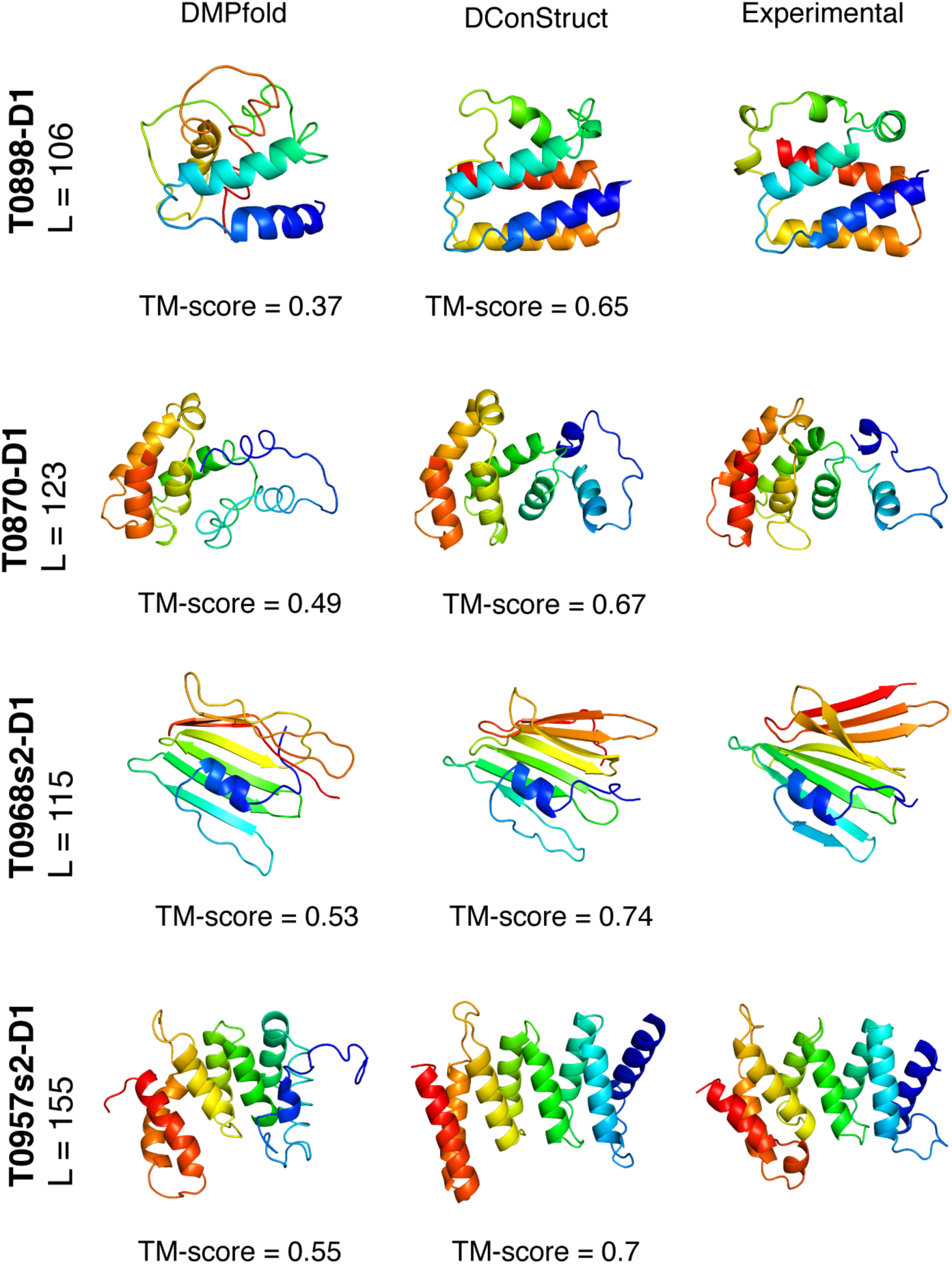
Ribbon diagrams of 3D models for the four CASP FM targets: T0898-D1, T0870-D1, T0968s2-D1, and T0957s2-D1; predicted by DMPfold and DConStruct along with the experimental structures. All molecules are rainbow colored blue to red from the N-to C-termini. Models are optimally superimposed on the experimental structures, and then separated by translations along the horizontal direction.

### Folding membrane proteins

Membrane proteins (MPs) have significant therapeutic values because of their importance in drug design [39]. However, only a small fraction of MPs are amenable to homology modeling, partially due to the lack of sufficient MPs with experimentally solved structures. Recently, the Xu group has developed a deep transfer learning (DTL) method for predicting the interaction maps from the sequences of the MPs that can be fed into CNS-based *ab initio* folding to predict the 3D structure of the MPs. The method (hereafter called Xu’s DTL with CNS) has demonstrated state-of-the-art performance on a dataset of 510 non-redundant MPs [31]. Noticing the ability of DConStruct to achieve improved *ab initio* folding accuracy for CASP FM targets, we examine whether DConStruct can improve *ab initio* folding accuracy of the MPs by exploiting distance-based hybrid interaction maps. For the 510 MPs, we follow the same protocol adopted for *ab initio* folding of CASP FM targets by obtaining the hybrid interaction maps from DMPfold predicted distance histograms and feeding them to DConStruct along with SPIDER3 predicted secondary structures for predicting the 3D structures of the MPs. We collect the top predicted 3D models using Xu’s DTL with CNS from the Mendeley Data provided in the published paper [31] to directly compare with DConStruct. As shown in **Table 4**, DConStruct attains an improved mean TM-score of 0.55, which is higher than Xu’s DTL with CNS (mean TM-score of 0.52). For DConStruct, the TM-scores have a median of 0.54, better than that of Xu’s DTL with CNS having a median TM-score of 0.5. The performance of DConStruct is statistically significantly better at 95% confidence level (*p*-value = 6.54e-06). Furthermore, DConStruct correctly folds 294 MP targets with a success rate of ∼58%, which is ∼8% higher than the success rate of Xu’s DTL with CNS (50%) that can fold only 255 MP targets correctly (**S6 Table**). The results confirm that DConStruct leads to improved *ab initio* folding accuracy even for MPs.

**Table 4.** Folding performance of 510 MPs using Xu’s deep transfer learning (DTL) with CNS and DConStruct. Values in bold represent the best performance.

| Method | Mean TM-scores | Median TM-score | # TM-score > 0.5 |
| --- | --- | --- | --- |
| DConStruct | **0.55** | **0.54** | **294/510** |
| Xu's DTL with CNS | 0.52 | 0.50 | 255/510 |

Fig 6 shows the 3D models predicted by Xu’s DTL with CNS and DConStruct for five MPs with lengths varying from 210 to 534 residues. For all five targets, DConStruct predicts correct overall fold attaining a TM-score of at least 0.7 and reaching as high as 0.84 TM-score for the target 4g1uA, whereas the Xu’s DTL with CNS fails to correctly fold any of the targets having a maximum TM-score of only 0.47.

**Fig 6.**
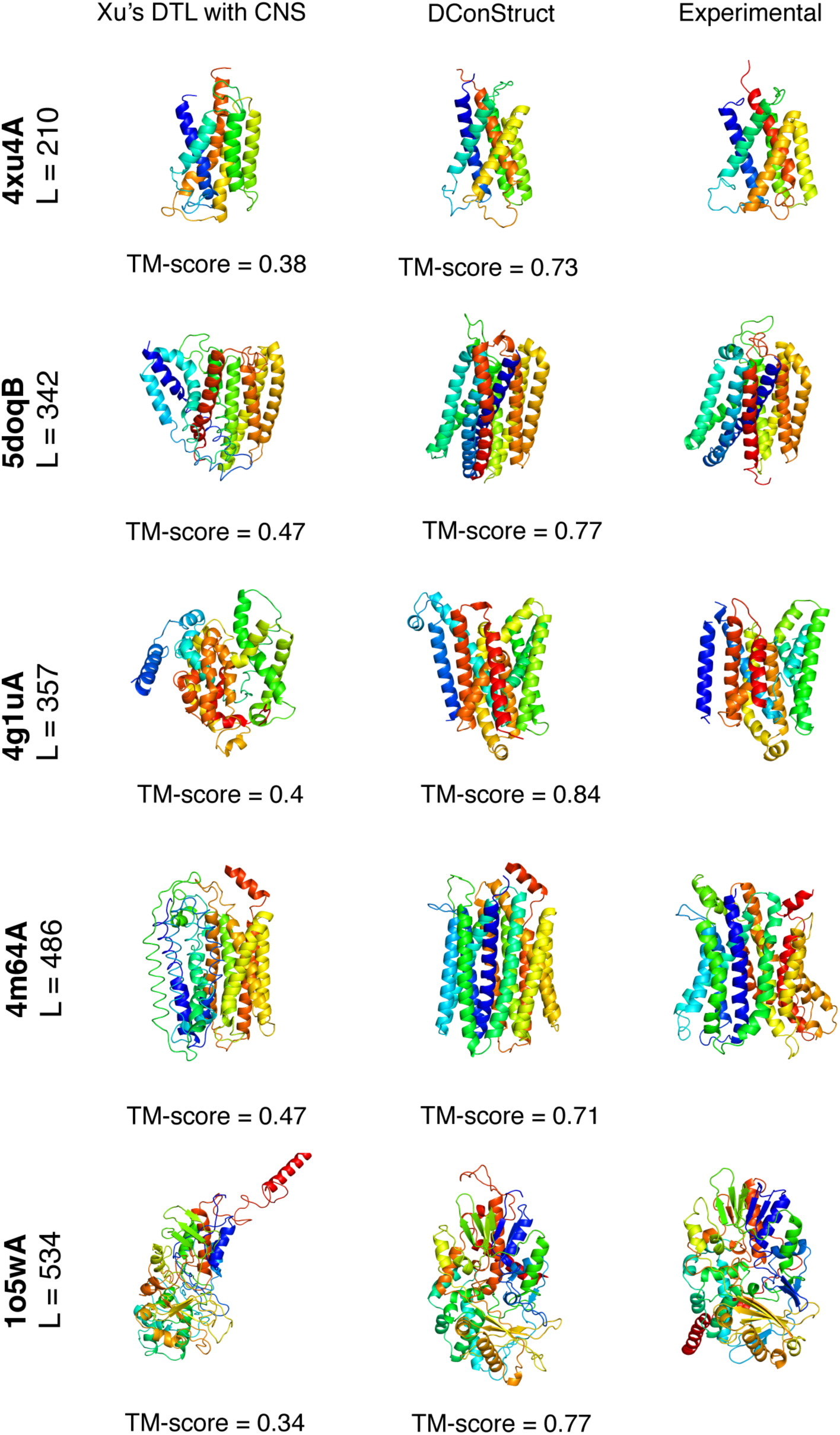
Ribbon diagrams of 3D models of MPs predicted using Xu’s DTL with CNS and DConStruct along with the experimental structures for five protein targets: PDB ID 4xu4 and chain A, PDB ID 5doq and chain B, PDB ID 4g1u and chain A, PDB ID 4m64 and chain A, and PDB ID 1o5w and chain A. All molecules are rainbow-colored blue to red from the N-to C-termini. Models are optimally superimposed on the experimental structures and then separated by translations along the horizontal direction.

### The three stages of DConStruct and their implications in hierarchical protein folding

To evaluate the relative contributions of the three stages adopted in DConStruct hierarchical structure modeling, we perform stage-by-stage 3D reconstruction of 15 protein targets tested in EVfold [16] and study their folding accuracy and correctness of secondary structure topology. For each protein, we use the true three-state secondary structure computed from the experimental structures using DSSP [36] and the true C_β_– C_β_ contact maps calculated at 8Å threshold having a minimum sequence separation of 6 residues. The rationale for using true input information is to prevent any bias caused by possible prediction noise in stage-wise hierarchical folding. In addition to measuring the stagewise folding accuracy using TM-score, we compute the percentage of correctly recovered secondary structure topology for the helix (Q_H_) and beta strand (Q_E_) residues.

As reported in **Table 5**, the mean TM-score after employing only stage 1 is 0.54 and 6 out of 15 targets fail to attain correct overall fold with poor secondary structure topology having a mean Q_H_ of ∼13% and a mean Q_E_ of only ∼1%. By introducing stage 2, the mean TM-score significantly improves to 0.76 with all proteins attaining correct overall folds. There is also a marked improvement in the helix topology having a mean Q_H_ of ∼59%, although beta strand topology still remains suboptimal at a mean Q_E_ of ∼13%. Stage 3 further improves the folding accuracy, attaining a mean TM-score of 0.83 but it is not as pronounced as the difference between stage 2 and stage 1. Much better mean Q_H_ of ∼94% is achieved with the introduction of stage 3, indicating near-optimal helix topology along with substantial improvement in beta strand topology having a mean Q_E_ of ∼62%. The results offer some interesting insights. First, as shown in Fig 7, the folding accuracy gain is substantial with the introduction of stage 2 with the TM-score distributions getting shifted to higher accuracy regions (0.22 mean TM-score gain from stage 1) compared to stage 3 (only 0.08 mean TM-score gain from stage 2), indicating that iterative self-correction with local structural perturbation is very effective in improving the overall fold-level accuracy. Second, stage 2 not only greatly improves the overall fold but also facilitates the formation of short-range hydrogen bonds, as demonstrated by significantly higher mean Q_H_ compared to stage 1. Third, while stage 3 has only minor contribution in boosting the overall fold-level accuracy, it optimizes the secondary structure topology through iterative self-correction by facilitating long-range hydrogen bonds formation for stabilizing the beta sheet geometry as revealed by much higher mean Q_E_ in addition to attaining near-optimal short-range hydrogen bonds by reaching close-to-optimal mean Q_H_. We note that the final mean Q_E_ is still far from optimal, indicating that there is room for improvement in accurately modeling the beta strands. In summary, the three stages of the hierarchical structure modeling adopted in DConStruct have complementary roles. Stage 2 is primarily responsible for improving the global folding accuracy beyond what can be achieved by the coarse-grained modeling in stage 1, whereas stage 3 is responsible for the refinement of global fold while stabilizing the local secondary structure topology. All three stages, working cooperatively in a hierarchic manner, contribute to enhancing 3D folding at both global and local levels.

**Fig 7.**
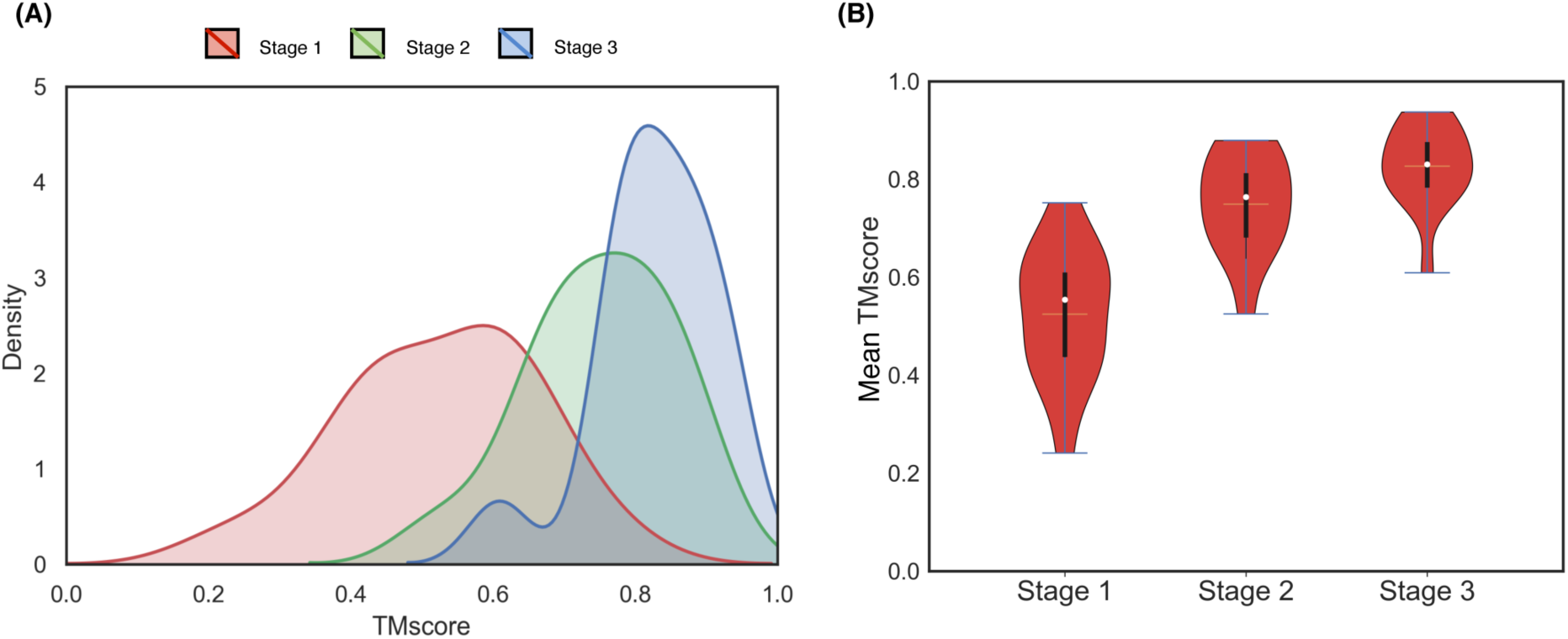
Stagewise TM-score distributions of the reconstructed models on 15 proteins from the EVfold set. Stage-by-stage (A) density plots and (B) violin plots are shown with the means indicated using the unfilled circles, the medians indicated using the horizontal yellow lines, and the interquartile ranges indicated using the vertical black strips.

**Table 5.** Stage-by-stage 3D reconstruction results for 15 protein targets in EVfold dataset using true C_β_-C_β_ contact maps at 8Å threshold and true secondary structures*.

| Target | Stage 1 |  |  | Stage 1 + Stage 2 |  |  | Stage 1 + Stage 2 + Stage 3 |  |  |
| --- | --- | --- | --- | --- | --- | --- | --- | --- | --- |
| | TM-score | $Q_H$ | $Q_E$ | TM-score | $Q_H$ | $Q_E$ | TM-score | $Q_H$ | $Q_E$ |
| 1bkrA | 0.75 | 37.14 | - | 0.87 | 74.29 | - | 0.90 | 98.57 | - |
| 1e6kA | 0.61 | 37.25 | 0.00 | 0.79 | 70.59 | 20.00 | 0.87 | 92.16 | 85.00 |
| 1f21A | 0.65 | 29.31 | 4.17 | 0.82 | 63.79 | 27.08 | 0.88 | 98.28 | 60.42 |
| 1g2eA | 0.43 | 14.29 | 0.00 | 0.67 | 80.95 | 0.00 | 0.77 | 100.00 | 36.00 |
| 1hzxA | 0.75 | 1.1 | 0.00 | 0.87 | 60.22 | 0.00 | 0.91 | 86.19 | 25.00 |
| 1oddA | 0.60 | 9.38 | 0.00 | 0.67 | 59.38 | 0.00 | 0.77 | 100.00 | 42.86 |
| 1r9hA | 0.44 | 0.00 | 0.00 | 0.74 | 42.86 | 0.00 | 0.82 | 100.00 | 77.78 |
| 1rqmA | 0.59 | 10.26 | 0.00 | 0.80 | 51.28 | 24.00 | 0.84 | 89.74 | 68.00 |
| 1wvnA | 0.45 | 19.36 | 0.00 | 0.71 | 74.19 | 17.65 | 0.78 | 100.00 | 70.59 |
| 2hdaA | 0.24 | - | 5.26 | 0.53 | - | 0.00 | 0.61 | - | 47.37 |
| 2it6A | 0.52 | 11.11 | 0.00 | 0.79 | 66.67 | 8.82 | 0.86 | 100.00 | 58.82 |
| 2o72A | 0.38 | - | 2.13 | 0.71 | - | 27.66 | 0.81 | - | 87.23 |
| 3tgiE | 0.64 | 0.00 | 2.63 | 0.88 | 0.00 | 13.16 | 0.94 | 57.14 | 71.05 |
| 5p21A | 0.61 | 0.00 | 0.00 | 0.88 | 75.81 | 5.13 | 0.93 | 98.39 | 74.36 |
| 5ptiA | 0.44 | 0.00 | 0.00 | 0.64 | 50.00 | 40.00 | 0.79 | 100.00 | 60.00 |
| **Mean** | **0.54** | **13.02** | **1.01** | **0.76** | **59.23** | **13.11** | **0.83** | **93.88** | **61.75** |

\* For target 1bkrA, $Q_E$ scores are ignored since there is no beta strand residue in the experimental structure; and for targets 2hdaA and 2o72A, $Q_H$ scores are ignored since there are no helix residues in their experimental structures.

A representative example from the EVfold dataset shown in Fig 8 may help further elucidate the relative contributions of the three stages used in DConStruct. This is an α/β protein of 165 residues (PDB ID: 5p21 chain A). While stage 1 is able to attain the correct overall fold with a TM-score of 0.61, the resulting model lacks any secondary structure with 0% Q_H_ and Q_E_. The addition of Stage 2 significantly improves the folding accuracy, attaining a TM-score of 0.88 (TM-score gain of 0.27 from stage 1), as well as the helix topology, pushing the Q_H_ to > 75%, but not so much in beta strand topology having a Q_E_ of only ∼5%. The introduction of stage 3 further boosts the TM-score to 0.93 (TM-score gain of 0.05 from stage 2) and achieves a near-optimal helix topology with a Q_H_ of > 98% along with significant enhancement in beta strand topology with a Q_E_ close to 75%. The pronounced improvement in the global fold-level accuracy caused by stage 2, and its further refinement together with the stabilization of local secondary structure topology caused by stage 3 is visually noticeable in Fig 8.

**Fig 8.**
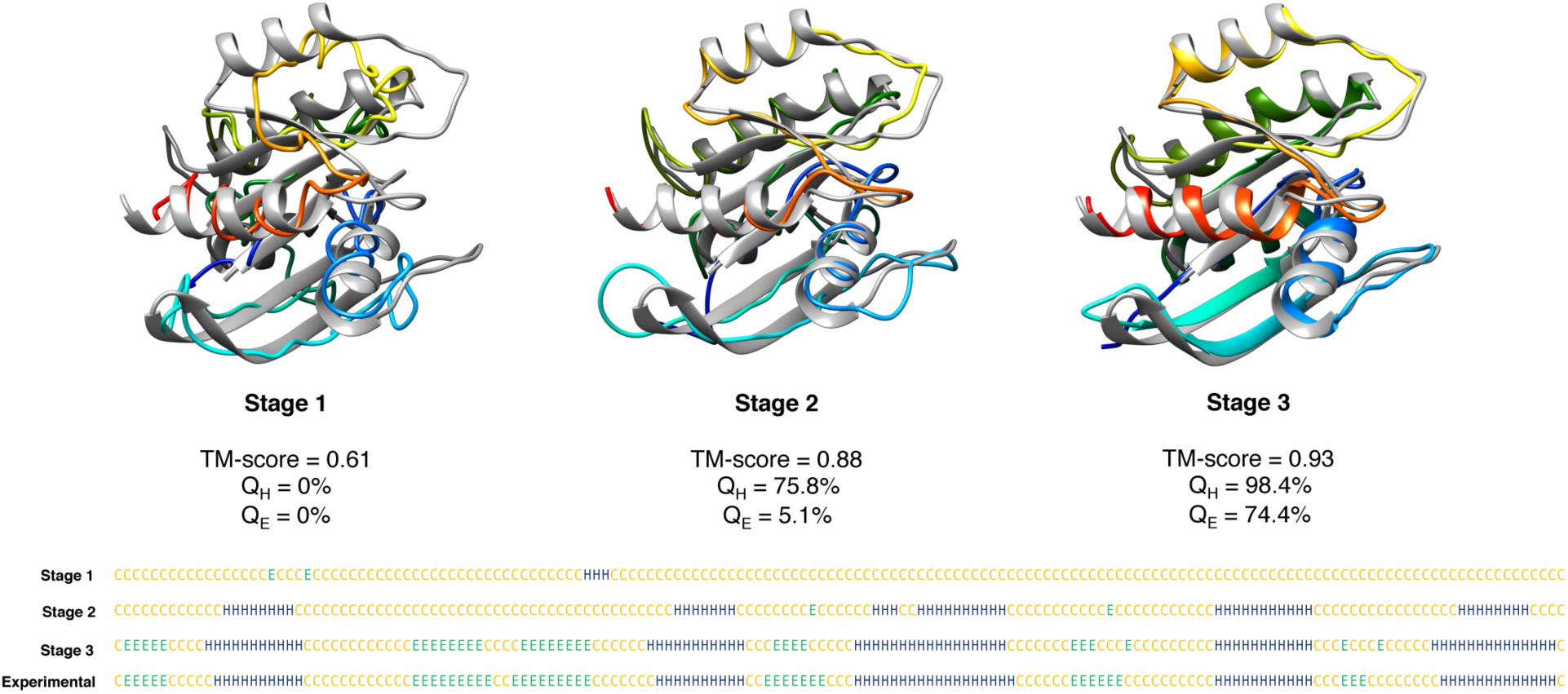
Visualization of effect of the three stages adopted in DConStruct for a representative protein target from EVfold dataset (PDB ID 5p21 and chain A).

### Why does DConStruct attain better accuracy at higher-thresholded contact maps?

An interesting finding is that DConStruct attains better folding performance for the interaction maps at higher contact thresholds such as 10 and 12Å. To unravel the underlying cause of such trend, we perform stage-by-stage 3D reconstruction on the same EVfold dataset comprising of 15 proteins by varying contact thresholds from 8 to 12Å in a step size of 2Å with true secondary structures, and study the gain in folding performance caused by the introduction of stage 2 (Δ_2_) and stage 3 (Δ_3_).

As reported in **Table 6** by using true C_β_-C_β_ contacts, the mean Δ_2_ (gain by stage 2) significantly increases with the increase in contact threshold in that Δ_2_ gets more than doubled from 0.22 at 8Å threshold to 0.46 at 12Å threshold. The mean Δ_3_ (gain by stage 3), on the other hand, remains relatively constant at around 0.07. This implies that stage 2 is the main driver of the improved folding accuracy for higher-thresholded contact maps (**S7 Table**). The results remain very similar when we repeat the experiment with the true C_α_-C_α_ contacts (**S8 Table**). To further understand whether an increase in distance threshold has any effect on the secondary structure topology, we study stage-wise Q_H_ and Q_E_ at varying contact thresholds from 8 to 12Å using both C_β_-C_β_ (**S9 Table**) and C_α_-C_α_ contacts (**S10 Table**). This time, stage 3 shows noticeable gain in beta strand topology with ∼10-15% increase in Q_E_ as contact threshold increases from 8 to 12Å, while the gain in Q_H_ achieved by stage 2 remains almost constant at higher-thresholded contact maps. In summary, these results further emphasize the cooperative aspect of the hierarchical modeling approach used in DConStruct, and indicate that the improved folding accuracy at higher-thresholded contact maps is the result of the combined improvement in the fold-level accuracy by stage 2 and enhancement in long-range hydrogen bonds formation for stabilizing the beta sheet geometry by stage 3.

**Table 6.** TM-score gain Δ_2_ (TM-score of stage 2 – TM-score of stage 1) and Δ_3_ (TM-score of stage 3 – TM-score of stage 2) after performing stage-by-stage 3D reconstruction for 15 protein targets in EVfold dataset using true C_β_-C_β_ contact maps at 8, 10, and 12Å thresholds and true secondary structures.

| Target | 8Å |  | 10Å |  | 12Å |  |
| --- | --- | --- | --- | --- | --- | --- |
| | $\Delta_2$ | $\Delta_3$ | $\Delta_2$ | $\Delta_3$ | $\Delta_2$ | $\Delta_3$ |
| 1bkrA | 0.12 | 0.04 | 0.27 | 0.06 | 0.46 | 0.06 |
| 1e6kA | 0.19 | 0.08 | 0.33 | 0.07 | 0.46 | 0.05 |
| 1f21A | 0.17 | 0.06 | 0.33 | 0.06 | 0.46 | 0.03 |
| 1g2eA | 0.24 | 0.10 | 0.37 | 0.07 | 0.48 | 0.09 |
| 1hzxA | 0.12 | 0.04 | 0.31 | 0.04 | 0.51 | 0.02 |
| 1oddA | 0.08 | 0.10 | 0.32 | 0.12 | 0.39 | 0.14 |
| 1r9hA | 0.30 | 0.08 | 0.38 | 0.06 | 0.52 | 0.05 |
| 1rqmA | 0.21 | 0.04 | 0.35 | 0.02 | 0.48 | 0.03 |
| 1wvnA | 0.26 | 0.07 | 0.35 | 0.05 | 0.42 | 0.05 |
| 2hdaA | 0.28 | 0.08 | 0.39 | 0.12 | 0.38 | 0.13 |
| 2it6A | 0.27 | 0.08 | 0.39 | 0.09 | 0.51 | 0.05 |
| 2o72A | 0.33 | 0.10 | 0.42 | 0.09 | 0.50 | 0.10 |
| 3tgiE | 0.24 | 0.06 | 0.37 | 0.04 | 0.48 | 0.03 |
| 5p21A | 0.27 | 0.05 | 0.34 | 0.04 | 0.48 | 0.05 |
| 5ptiA | 0.20 | 0.15 | 0.33 | 0.12 | 0.38 | 0.12 |
| **Mean** | **0.22** | **0.08** | **0.35** | **0.07** | **0.46** | **0.07** |

## Discussion

This article has presented a new hybridized distance- and contact-based hierarchical structure modeling method DConStruct that can greatly improve *ab initio* protein folding. In contrast to the existing folding approaches, our method neither depends on the CNS experimental structure determination protocol nor performs time-consuming fragment-based conformational sampling, but rather employs 3-stage hierarchical structure modeling driven purely by the geometric restraints induced by inter-residue interaction maps and secondary structures. Our new predictive modeling method DConStruct is unique in that it can hierarchically estimate the correct overall fold of a target protein in coarse-grained mode to progressively optimize the local and non-local interactions while enhancing secondary structure topology in a self-correcting manner. Rigorous experimental validation reveals that DConStruct not only attains much better contact-driven reconstruction performance than currently popular CNS-based approaches, but also scales at a much higher degree of folding accuracy than CNS with the increase in contact thresholds. DConStruct is also versatile in that it can exploit the information encoded in the interaction maps at multiple granularities ranging from binary contact maps to distance-based hybrid maps at tri-level thresholding, which results in better *ab initio* folding performance for CASP12 and CASP13 FM targets compared to several popular CNS- and fragment-based approaches. Even without using fine-grained distance maps or fragment assembly, *ab initio* folding using DConStruct can yield the correct fold for more CASP FM targets than state-of-the-art approaches. Further, our experimental results show that DConStruct leads to better folding accuracy for membrane proteins compared to a CNS-based approach. We expect that our new structure modeling method can enhance the accuracy of distance- or contact-driven folding of many more non-homologous proteins that lack experimental structures, thereby facilitating structure-based studies for additional protein families leading to new biological insights.

Our method outperforms CNS due to a couple of reasons. First, CNS combines the restraints derived from the interaction maps with an in-built biophysical force field having parameters fine-tuned for experimental data, while our method is free from the influence of such force fields. Second, CNS adopts an all-atom representation throughout the folding simulation, while our new method DConStruct follows a hierarchic approach by first estimating the correct overall fold in coarse-grained mode and then progressively optimizing the local and non-local interactions in atomistic detail. As such, the adaptation of coarse-grained representation in DConStruct during the early stages of folding significantly reduces the conformational space, accelerating the estimation of the overall fold driven purely by the inter-residue interactions defined between the C_α_–C_α_ or C_β_–C_β_ atom pairs. Finally, the introduction of additional folding stages in DConStruct cooperatively improves the overall fold and facilitates hydrogen bonds formation for stabilizing the secondary structure topologies, ultimately resulting in improved folding performance compared to CNS.

The hierarchical structure modeling paradigm employed in DConStruct attains better folding accuracy than CNS-based CONFOLD protocol for higher-thresholded contact maps at 10 and 12Å, beyond the standard contact threshold of 8Å currently used by the community. Experimental results show that this behavior is attributed to the cooperative nature of the 3-stage hierarchical modeling approach adopted in DConStruct, with stage 2 further improving the fold-level accuracy with the increase in contact threshold than the initial fold estimated by the coarse-grained modeling from stage 1, and stage 3 better enhancing long-range hydrogen bonds formation for stabilizing the beta sheet geometry at higher-thresholded contact maps. Moreover, feeding hybrid interaction maps at tri-level thresholding that combines contact maps at 8, 10, and 12Å thresholds further improves the performance of DConStruct, opening the unique possibility of utilizing hybridized distance and contact maps for predictive protein modeling. Indeed, our hybridized distance- and contact-based hierarchical folding method DConStruct delivers better performance in *ab initio* folding of CASP free modeling targets compared to CNS-based CONFOLD2 and DMPfold protocols as well as ROSETTA-based fragment assembly pipeline PconsFold and recent fragment-based CGLfold method. DConStruct convincingly outperforms the state-of-the-art DMPfold protocol, which uses much finer-grained distance maps along with additional predicted structural features such as torsion angles and hydrogen bonds, whereas our method utilizes only hybrid interaction maps at tri-level thresholding derived from the DMPfold-predicted initial distance histograms. The better predictive modeling performance of DConStruct also translates to superior *ab initio* folding of membrane proteins compared to the state of the art. In summary, the advantage of DConStruct in *de novo* protein modeling over the others is significant.

We may further improve the folding accuracy of our new method by extending the hierarchical structure modeling by allowing finer-grained distance intervals, which contain more information than what we are using now. For example, we may directly feed the 20 distance bins from DMPfold with their likelihood values to DConStruct during predictive modeling. Needless to say, accurate estimation of finer-grained distance maps is a very challenging problem, which may suffer from high false-positive rates and thus prone to be noisy. Empowering our method to be robust and noise-tolerant while utilizing finer-grained distance intervals shall result in much better 3D models. We may also improve the hierarchical structure modeling by performing enhanced sampling targeted at the regions not restrained by an input interaction maps. This may improve the quality of the flexible loops or terminal regions, which may not be proximal to the core regions, ultimately enhancing the overall folding accuracy. Finally, in addition to secondary structure, contact and distance information, our method can be extended to use other structural features such as solvent accessibility and disulfide bridges, which contain complementary information about protein conformation and thus, shall benefit predictive structure modeling.

## Methods

### Hierarchical structure modeling employed in DConStruct

#### From interaction map and secondary structure to proximity map

Our folding protocol starts by estimating the proximity map from a given interaction map and secondary structure using knowledge-based and graph-theoretic approach. Simply speaking, a proximity map is an approximation of the inter-residue distance matrix. Using a coarse-grained string of beads representation, in which each bead corresponds to the C_α_ atom of an amino acid residue, we first estimate the proximity between the residue pairs close in sequence from prior knowledge of protein backbone geometry derived from the pseudo-covalent bonds formed between the C_α_ atoms and the secondary structure-specific local structural preferences [21, 35], thus generating a sparse proximity map. For sequentially distant residue pairs, we apply a graph-theoretic approach to fill the missing entries in the proximity map. We treat the input interaction map as an adjacency matrix representing a graph G = (V, E), where V = {v_1_, v_2_, …} is the set of nodes, representing a residue’s C_α_ (or C_β_) atom, and E = {e_ij_} is the set of edges, where e_ij_ represents the interaction (e.g., residue-residue contact) between v_i_ and v_j_. Thus, the graph G encapsulates the mathematical relation of inter-residue spatial proximity in 3D space. A graph distance function [40, 41], defined as the shortest path length among all paths for any given residue pair, can be applied to the graph that shall be measured for the entire set of V. The path length can be calculated by summing the total number of edges connecting any pair of residues (v_i_, v_j_) under consideration. This function approximates the spatial distances and can generate an initial estimate of the proximity map. Here we use Floyd-Warshall all pair shortest path algorithm [42] to compute the path length. We further refine the initial estimate of the proximity map by idealized secondary structure element (SSE) modeling. SSEs are identified from the given secondary structure and are independently modeled by setting the sequence of pseudo angles and pseudo dihedral angels spanning each SSE to their ideal values [21, 43]. The SSEs, modeled in angular space, are subsequently converted into Cartesian space assuming the distance between two consecutive residues is constant (3.8Å) to extract the intra-SSE-all-residue-pairs distances for refining the initial proximity map. The rationale for this step is to enhance the physical realism of the proximity map for the intra-SSE segments.

#### From proximity map to coarse-grained 3D models

We turn the proximity map into a 3D model using an efficient graph realization technique. The basic idea is to treat the graph distance-based proximity information between C_α_ atoms for each residue pair to calculate the corresponding coordinates for all residues by applying the graph-based method, multidimensional scaling (MDS) [22–24], for generating coarse-grained 3D models. MDS starts with dissimilarity matrices that are derived from points in a multidimensional space, and it finds the positioning of the points in a low-dimensional space, where the distances between points resemble the original dissimilarities. Here, we use the proximity map between the C_α_ atoms of each residue pair as dissimilarity matrix and then calculate the coordinates of the C_α_ atom for all residue. If there are n points (each representing the C_α_ atom of a residue) X_k_ ∈ R^3^, k = 1,…,n in 3D. Here we use classical metric MDS (CMDS) [44], which is the first and simplest MDS algorithm. For a perfect dissimilarity (distance) matrix without any error in the Euclidean space, CMDS will exactly reconstruct the configuration of points (or its mirror configuration) with a computational complexity of O(n^3^). But, when there are errors in the dissimilarity matrix, CMDS minimizes the sum of least squared errors between the estimated and the observed distances in the output model for all pairs of points. In practice, the technique gracefully tolerates errors due to the overdetermined nature of the solutions. This is important in our case, as our graph-based proximity map is an approximation of the true Euclidean distance matrix that can be noisy and physically unrealistic, particularly for sequentially distant residue pairs. We iteratively apply this technique to produce an initial pool of 20 coarse-grained 3D models.

#### Improving coarse-grained models through iterative self-correction

We apply iterative self-correction via local perturbation to further optimize the coarse-grained models. Specifically, we identify the residues having inconsistent spatial positioning with respect to the rest of the structure and employ coordinate refinement heuristics [45] to correct its coordinate without introducing new error in the coordinate set. For the residue-pairs having their Euclidean distances close to the distance threshold of the input interaction map, we apply corrective coordinate perturbation to maximally reproduce contacts and non-contacts. We also perform local perturbation for intra-SSE-segments by coordinate adjustments without affecting other correctly positioned residues outside of SSE. Furthermore, we enforce idealized pseudo-covalent bond lengths constraints formed between consecutive C_α_ atoms (3.8Å), and steric clash constraints (defined as two C_α_ atoms that are closer than 3.5Å from each other). Finally, we employ Limited-memory Broyden-Fletcher-Goldfarb-Shannon (L-BFGS) [25] algorithm for minimizing a harmonic function of the observed and expected distance values of C_α_ quadruplets (i – 1, i + 1, i + 2, i+ 3); with distances between the (i - 1)th residue and the (i + 1)th, the (i + 2)th, and the (i+ 3)th residues, respectively. The expected values specific to helices and strands are adopted from [21] to refine the local secondary structure topology. The entire self-correction process described above is iteratively applied to generate a pool of 20 optimized 3D models.

#### Contact violation-based model selection and geometric chirality checking

For the selection of one representative structure from the optimized model pool, we use a contact violation-based score function that combines contact or non-contact satisfaction of a model as: F = ∑ contact-error + ∑ non-contact-error + n’, where contact- and non-contact-errors are calculated using a squared error function, defined as (d_ij_ – threshold)^2^ for residue-pair (i, j) with distance d_ij_ that violates input contacts (or non-contacts), and n’ is the number of residue pairs inconsistent with the input interaction map. We rank the pool of optimized models using this score function to select the highest ranked model as the representative.

One remaining issue in our selected model is that it can be a mirror image of the biologically relevant 3D conformation that has the correct chirality or handedness at the local level. This is a typical issue faced during the reconstruction of 3D structure from a 2D representation since any distance function is invariant to the isometric transformations in 3D: translation, rotation and symmetry. As such, it is not obvious which of the two structures related by mirror symmetry represents a biological relevant structure. Nonetheless, secondary structure can be used to identify the correct chirality of a 3D conformation since the observed chirality is mostly right-handed for α-helices and left-handed β-sheets [26]. We define a geometric cost function for identifying the correct chirality as the sum over tetrapeptides in α-helices and β-sheets by utilizing the secondary structure-specific normalized triple scalar product values adopted from [26]. From among the two mirror images of the selected model, we select the structure with lowest cost as the correct chirality.

#### Atomic-level model building and iterative self-correction

We use MODELLER [27] to generate atomic-level model and perform restraint satisfaction using the secondary structure and interaction map-derived geometric restraints to generate optimized atomic-level model, and subsequently employ restraint satisfaction iteratively using unsatisfied interactions, non-interactions, and secondary structure restraints for self-correction along with model combination to generate the final folded conformation.

### Test datasets and programs to compare

For the 3D reconstruction from the true contact-based interaction map and true secondary structure, we use 150 soluble proteins from FRAGFOLD [28] dataset with sequence length ranging from 50 to 266 residues. We extract the true contact maps from the experimental structures having a sequence separation of at least 6 residues by varying the contact thresholds from 8 to 12Å in a step size of 0.5Å. We use DSSP [36] program to compute the true secondary structures from the experimental structures. We compare DConStruct with two widely used reconstruction methods, FT-COMAR [35] and CONFOLD [17], using the same input. FT-COMAR is a fast and purely distance geometry-based heuristic method that reconstructs the C_α_ trace purely from a given C_α_– C_α_ contact matrix and does not accept secondary structure. On the other hand, CNS-based CONFOLD reconstructs protein 3D structure using a contact map and a secondary structure via a two-stage process. For CONFOLD-based reconstruction, we set the parameters of CONFOLD as ‘-stage2 3’ (model generation at stage 2 using sheet-detection only for true contacts), ‘-rep 0.8’ (used for true contact), ‘-rrtype cb’ (for C_β_–C_β_ contacts) or ‘-rrtype ca’ (for C_α_–C_α_ contacts). Furthermore, we extract the true distance-based hybrid interaction maps (both C_α_–C_α_ and C_β_–C_β_) from the experimental structures and use them together with their true secondary structures for 3D reconstruction using DConStruct.

For *ab initio* folding, 40 free modeling (FM) domains from CASP12 and CASP13 with publicly available experimental structures are used with target lengths ranging from 67 to 356 residues. The performance of DConStruct is compared with DMPfold [8], CONFOLD2 [18], ROSETTA [13] and CGLFold [15]. DMPfold is a deep learning-based CNS-dependent *ab initio* folding method. CONFOLD2 is a modified version of CONFOLD based on CNS-based distance geometry algorithm. For *ab initio* folding using CONFOLD2, RaptorX predicted top 2L contacts, obtained by submitting jobs to its web server, are used according to the published paper [4], together with the secondary structures predicted using SPIDER3 [33]. For ROSETTA, we adopt a similar process as in PconsFold [14] protocol with the only exception of using RaptorX contacts instead of PconsC [46]. We obtain fragments (3-mers and 9-mers) from the ROBETTA server [47] to generate a pool of 2,000 models with a maximum duration of 15 calendar days. The lowest ROSETTA energy models are subsequently selected as the prediction. CGLFold [15] is a recent fragment-based method that combines global exploration and loop perturbation using the predicted contact maps from ResTriplet [48]. The TM-scores for CGLFold 29 FM targets are collected from its published paper. To generate distance-based hybrid interaction maps for DConStruct, we use the DMPfold distance predictor. We feed the multiple sequence alignments (MSAs) [34] to DMPfold and obtain the predicted initial distance histograms (rawdispred.current files) containing 20 distance bins with their associated likelihoods (we do not run any further DMPfold iterations involving CNS-based modeling). The predicted histograms are then converted to hybrid interaction maps with tri-level thresholding with variable upper bounds of 8, 10, and 12Å by summing up the likelihoods for distance bins below the three distance thresholds of 8,10, and 12Å. We select the top 8L C_β_–C_β_ high confidence interacting residue pairs having likelihoods > 0.85 since using the top 8L C_β_–C_β_ contacts delivers the best performance in an independent benchmarking on the EVfold dataset (**S11 Table**) when experimented with top 2L, 4L, 8L, and 16L C_β_–C_β_ contacts. The hybrid interaction maps coupled with the SPIDER3 predicted secondary structures are then fed to DConStruct for *ab initio* folding.

We also evaluate *ab initio* folding of membrane proteins using 510 non-redundant membrane proteins used in [31] with length ranging from 50 to 646 residues. We generate the distance-based hybrid interaction maps using DMPfold as mentioned above and predict their 3D structures using DConStruct to compare our models with that of Xu’s deep transfer learning (DTL) and CNS-based *ab initio* folding method [31].

To evaluate the implications of the 3-stage hierarchical structure modeling approach adopted in DConStruct, we use 15 proteins from the EVfold [16] dataset ranging from 48 to 248 residues in length. We use true contact interactions with a sequence separation of at least 6 residues and true secondary structures to perform stage-by-stage 3D reconstruction and analysis after using the PULCHRA program [49] to generate all-atom models from the coarse-grained models produced in the intermediate stages.

### Evaluation metrics

We use TM-score [37] to evaluate the folding accuracy. TM-score compares the predicted models with the experimental structure to determine their structural similarity. TM-score has the value in (0, 1], with higher value indicating better folding accuracy. Meanwhile, TM-score > 0.5 represents correctly folded models [38]. We measure the correctness of secondary structure topology using the percentage of correctly recovered secondary structure residues for helix (Q_H_) and beta-strand (Q_E_) residues.

## Supporting Information

**S1 Table.** Target-by-target reconstruction performance on 150 soluble proteins for true C_α_–C_α_ contact maps at various thresholds.

**S2 Table.** Target-by-target reconstruction performance on 150 soluble proteins for true C_β_–C_β_ contact maps at various thresholds.

**S3 Table.** Target-by-target reconstruction performance on 150 soluble proteins for true C_α_–C_α_ hybrid interaction maps at tri-level thresholding.

**S4 Table.** Target-by-target reconstruction performance on 150 soluble proteins for true C_β_–C_β_ hybrid interaction maps at tri-level thresholding.

**S5 Table.** Target-by-target *ab initio* folding performance on 40 CASP FM targets.

**S6 Table.** Target-by-target *ab initio* folding performance on 510 membrane proteins.

**S7 Table.** Target-by-target stagewise reconstruction performance on EVfold dataset for true C_β_–C_β_ contact maps at 8, 10, and 12Å thresholds.

**S8 Table.** Target-by-target stagewise reconstruction performance on EVfold dataset for true C_α_–C_α_ contact maps at 8, 10, and 12Å thresholds.

**S9 Table.** Target-by-target stagewise recovery of secondary structure topology on EVfold dataset for true C_β_–C_β_ contact maps at 8, 10, and 12Å thresholds.

**S10 Table.** Target-by-target stagewise recovery of secondary structure topology on EVfold dataset for true C_α_–C_α_ contact maps at 8, 10, and 12Å thresholds.

**S11 Table.** *Ab initio* folding performance of DConStruct on EVfold dataset using top hybrid interaction maps with tri-level thresholding at increasing xL values (x = 2, 4, 8, 16).

## Acknowledgements

This work was made possible in part by a grant of high performance computing resources and technical support from the Alabama Supercomputer Authority and Auburn University Early Career Development Grant to DB.

