## Supporting information for "Hybridized distance- and contact-based hierarchical structure modeling for folding soluble and membrane proteins"

**S1 Table.** Target-by-target reconstruction performance on 150 soluble proteins for true $C_\alpha$–$C_\alpha$ contact maps at various thresholds.

| Target | 8 Å | | | 8.5 Å | | | 9 Å | | | 9.5 Å | | | 10 Å | | | 10.5 Å | | | 11 Å | | | 11.5 Å | | | 12 Å | | |
|---|---|---|---|---|---|---|---|---|---|---|---|---|---|---|---|---|---|---|---|---|---|---|---|---|---|---|---|
| | FT-COMAR | CONFOLD | DConStruct | FT-COMAR | CONFOLD | DConStruct | FT-COMAR | CONFOLD | DConStruct | FT-COMAR | CONFOLD | DConStruct | FT-COMAR | CONFOLD | DConStruct | FT-COMAR | CONFOLD | DConStruct | FT-COMAR | CONFOLD | DConStruct | FT-COMAR | CONFOLD | DConStruct | FT-COMAR | CONFOLD | DConStruct |
| 1a3aA | 0.3212 | 0.8411 | 0.9292 | 0.8245 | 0.8383 | 0.9046 | 0.3303 | 0.8463 | 0.9333 | 0.3425 | 0.865 | 0.9655 | 0.3353 | 0.8921 | 0.9669 | 0.3447 | 0.881 | 0.9613 | 0.3402 | 0.8879 | 0.9721 | 0.3429 | 0.9072 | 0.9712 | 0.9667 | 0.8788 | 0.97 |
| 1a6mA | 0.5619 | 0.8454 | 0.8619 | 0.5882 | 0.904 | 0.9288 | 0.3112 | 0.915 | 0.9369 | 0.7175 | 0.925 | 0.9388 | 0.903 | 0.9138 | 0.9442 | 0.939 | 0.9359 | 0.9698 | 0.9572 | 0.9483 | 0.9706 | 0.9631 | 0.9385 | 0.9741 | 0.358 | 0.9366 | 0.9757 |
| 1a70A | 0.2861 | 0.7759 | 0.843 | 0.7599 | 0.7819 | 0.8181 | 0.801 | 0.8767 | 0.9042 | 0.2909 | 0.8697 | 0.9203 | 0.3073 | 0.8779 | 0.9581 | 0.3013 | 0.8088 | 0.9559 | 0.962 | 0.8829 | 0.9641 | 0.9331 | 0.9088 | 0.9654 | 0.3088 | 0.822 | 0.9549 |
| 1aapA | 0.2937 | 0.7184 | 0.7912 | 0.3087 | 0.7616 | 0.8132 | 0.3315 | 0.7452 | 0.8134 | 0.2993 | 0.7424 | 0.8342 | 0.3629 | 0.7452 | 0.8845 | 0.8061 | 0.792 | 0.8452 | 0.3247 | 0.8021 | 0.8553 | 0.3219 | 0.783 | 0.8659 | 0.3518 | 0.7599 | 0.8162 |
| 1abaA | 0.3167 | 0.7352 | 0.8666 | 0.3002 | 0.7662 | 0.8441 | 0.558 | 0.797 | 0.8849 | 0.3228 | 0.8576 | 0.8714 | 0.8337 | 0.7911 | 0.9109 | 0.7975 | 0.8393 | 0.9086 | 0.346 | 0.8354 | 0.9287 | 0.3637 | 0.8246 | 0.9232 | 0.8966 | 0.8499 | 0.9138 |
| 1ag6A | 0.7028 | 0.8527 | 0.8682 | 0.7219 | 0.7837 | 0.8758 | 0.805 | 0.8096 | 0.9019 | 0.856 | 0.8567 | 0.8911 | 0.8965 | 0.822 | 0.9396 | 0.3203 | 0.8141 | 0.9585 | 0.941 | 0.7961 | 0.9609 | 0.3241 | 0.9092 | 0.9555 | 0.3283 | 0.8393 | 0.9594 |
| 1aoeA | 0.3166 | 0.7831 | 0.8943 | 0.3157 | 0.7803 | 0.9128 | 0.3188 | 0.8481 | 0.9345 | 0.3468 | 0.8489 | 0.9491 | 0.3464 | 0.8859 | 0.9673 | 0.3492 | 0.9167 | 0.9703 | 0.3485 | 0.886 | 0.9773 | 0.9523 | 0.8898 | 0.9812 | 0.3522 | 0.8814 | 0.9731 |
| 1atlA | 0.3242 | 0.9097 | 0.9435 | 0.3329 | 0.8863 | 0.9331 | 0.351 | 0.9075 | 0.9509 | 0.9185 | 0.9391 | 0.9702 | 0.3508 | 0.9281 | 0.9827 | 0.3603 | 0.9429 | 0.9827 | 0.3561 | 0.9477 | 0.9839 | 0.3542 | 0.943 | 0.9855 | 0.355 | 0.9413 | 0.9839 |
| 1atzA | 0.2995 | 0.7568 | 0.819 | 0.2876 | 0.781 | 0.8363 | 0.3182 | 0.79 | 0.8843 | 0.7094 | 0.835 | 0.8925 | 0.7585 | 0.7471 | 0.9183 | 0.7954 | 0.8324 | 0.9333 | 0.8656 | 0.8346 | 0.9328 | 0.838 | 0.8318 | 0.9276 | 0.8734 | 0.8202 | 0.9343 |
| 1avsA | 0.4318 | 0.693 | 0.7493 | 0.5223 | 0.734 | 0.7926 | 0.273 | 0.6514 | 0.7684 | 0.7365 | 0.7907 | 0.8796 | 0.7991 | 0.775 | 0.8845 | 0.3163 | 0.8503 | 0.8873 | 0.8653 | 0.8805 | 0.9432 | 0.8292 | 0.899 | 0.9339 | 0.3213 | 0.8471 | 0.92 |
| 1bdoA | 0.606 | 0.7337 | 0.8448 | 0.2891 | 0.7373 | 0.8313 | 0.2779 | 0.7662 | 0.8696 | 0.7867 | 0.7607 | 0.8983 | 0.8235 | 0.6747 | 0.9166 | 0.8421 | 0.7623 | 0.9133 | 0.8504 | 0.7174 | 0.9317 | 0.8657 | 0.7392 | 0.9446 | 0.9077 | 0.7389 | 0.9378 |
| 1bebA | 0.2926 | 0.7749 | 0.8591 | 0.3125 | 0.8464 | 0.8773 | 0.3195 | 0.8436 | 0.9088 | 0.3234 | 0.8464 | 0.9492 | 0.3208 | 0.8611 | 0.9731 | 0.333 | 0.8245 | 0.976 | 0.3302 | 0.8028 | 0.979 | 0.3275 | 0.8883 | 0.9768 | 0.3335 | 0.8785 | 0.9783 |
| 1behA | 0.7909 | 0.8196 | 0.87 | 0.3186 | 0.8584 | 0.9379 | 0.8991 | 0.8543 | 0.9436 | 0.3485 | 0.8802 | 0.9606 | 0.9491 | 0.8844 | 0.9728 | 0.3449 | 0.8803 | 0.971 | 0.3443 | 0.9006 | 0.9783 | 0.3439 | 0.8854 | 0.9708 | 0.349 | 0.9045 | 0.9799 |
| 1bkrA | 0.5168 | 0.8712 | 0.9018 | 0.3112 | 0.9223 | 0.9293 | 0.307 | 0.8915 | 0.9371 | 0.8439 | 0.906 | 0.9405 | 0.3192 | 0.9171 | 0.9504 | 0.918 | 0.9291 | 0.967 | 0.3188 | 0.9191 | 0.9698 | 0.3221 | 0.9423 | 0.9644 | 0.3233 | 0.942 | 0.9755 |
| 1brfA | 0.515 | 0.6103 | 0.6649 | 0.5728 | 0.6885 | 0.7457 | 0.579 | 0.6816 | 0.2795 | 0.7657 | 0.7325 | 0.788 | 0.7543 | 0.6978 | 0.8561 | 0.3046 | 0.7548 | 0.8562 | 0.3292 | 0.7229 | 0.8811 | 0.7697 | 0.7565 | 0.8539 | 0.3229 | 0.7989 | 0.8811 |
| 1bsgA | 0.8443 | 0.901 | 0.9425 | 0.3557 | 0.9129 | 0.9522 | 0.907 | 0.9419 | 0.9769 | 0.9472 | 0.9355 | 0.9798 | 0.9771 | 0.9575 | 0.9824 | 0.381 | 0.9575 | 0.9877 | 0.3773 | 0.9525 | 0.9836 | 0.3778 | 0.9593 | 0.9897 | 0.9866 | 0.971 | 0.9888 |
| 1c44A | 0.6927 | 0.8009 | 0.7989 | 0.7823 | 0.7826 | 0.8522 | 0.8317 | 0.8512 | 0.8936 | 0.8493 | 0.8479 | 0.8921 | 0.908 | 0.8519 | 0.8987 | 0.8993 | 0.8674 | 0.9077 | 0.937 | 0.8561 | 0.9222 | 0.9252 | 0.871 | 0.9267 | 0.3218 | 0.8491 | 0.9285 |
| 1c52A | 0.2557 | 0.7659 | 0.8481 | 0.2767 | 0.8283 | 0.8964 | 0.7326 | 0.8658 | 0.9018 | 0.8638 | 0.887 | 0.9568 | 0.9287 | 0.8922 | 0.9601 | 0.9152 | 0.872 | 0.9583 | 0.3007 | 0.9028 | 0.9668 | 0.3048 | 0.8494 | 0.9645 | 0.9646 | 0.9062 | 0.9663 |
| 1c9oA | 0.6002 | 0.6976 | 0.7547 | 0.6725 | 0.6926 | 0.734 | 0.2696 | 0.695 | 0.8186 | 0.2823 | 0.7125 | 0.8922 | 0.2858 | 0.7726 | 0.91 | 0.2829 | 0.7287 | 0.9049 | 0.2763 | 0.6466 | 0.8856 | 0.8576 | 0.6862 | 0.9231 | 0.2749 | 0.7542 | 0.914 |
| 1cc8A | 0.2686 | 0.7619 | 0.8234 | 0.3042 | 0.7195 | 0.8354 | 0.6913 | 0.817 | 0.9017 | 0.2898 | 0.8044 | 0.9078 | 0.8229 | 0.8106 | 0.8955 | 0.3268 | 0.843 | 0.9301 | 0.3307 | 0.8346 | 0.9375 | 0.3395 | 0.7471 | 0.9386 | 0.9161 | 0.8459 | 0.9338 |
| 1chdA | 0.3054 | 0.9108 | 0.9367 | 0.3155 | 0.933 | 0.9517 | 0.9061 | 0.9439 | 0.9494 | 0.3287 | 0.9415 | 0.9651 | 0.3218 | 0.9597 | 0.9724 | 0.9618 | 0.9454 | 0.9793 | 0.327 | 0.9521 | 0.9839 | 0.9643 | 0.9601 | 0.984 | 0.983 | 0.9433 | 0.9805 |
| 1cjwA | 0.2835 | 0.8031 | 0.9026 | 0.2979 | 0.783 | 0.9267 | 0.816 | 0.847 | 0.9382 | 0.857 | 0.8074 | 0.9423 | 0.3218 | 0.8274 | 0.9534 | 0.9348 | 0.8585 | 0.9514 | 0.3387 | 0.8103 | 0.9616 | 0.3315 | 0.8493 | 0.9672 | 0.9606 | 0.8744 | 0.9669 |
| 1ckeA | 0.5513 | 0.8502 | 0.8527 | 0.5989 | 0.878 | 0.8409 | 0.7206 | 0.8259 | 0.8636 | 0.7847 | 0.8528 | 0.8766 | 0.831 | 0.8799 | 0.9024 | 0.3576 | 0.8627 | 0.8996 | 0.9312 | 0.8777 | 0.9104 | 0.9251 | 0.8847 | 0.9189 | 0.9516 | 0.8757 | 0.9205 |
| 1ctfA | 0.2767 | 0.6762 | 0.7485 | 0.2923 | 0.7662 | 0.8123 | 0.3009 | 0.7616 | 0.89 | 0.3175 | 0.793 | 0.8825 | 0.9333 | 0.8245 | 0.8947 | 0.8832 | 0.8068 | 0.8944 | 0.3042 | 0.8246 | 0.8967 | 0.3174 | 0.8271 | 0.9171 | 0.3216 | 0.8156 | 0.9284 |
| 1cxyA | 0.2634 | 0.5331 | 0.621 | 0.4244 | 0.6622 | 0.709 | 0.6262 | 0.7332 | 0.7614 | 0.2843 | 0.693 | 0.8354 | 0.3114 | 0.7905 | 0.8975 | 0.2897 | 0.8355 | 0.9073 | 0.3035 | 0.8372 | 0.9384 | 0.301 | 0.7798 | 0.9343 | 0.8883 | 0.7534 | 0.9278 |
| 1cznA | 0.6909 | 0.8855 | 0.9117 | 0.3067 | 0.886 | 0.9267 | 0.3091 | 0.898 | 0.9429 | 0.9095 | 0.9082 | 0.9589 | 0.325 | 0.9416 | 0.9645 | 0.3295 | 0.9224 | 0.9659 | 0.3347 | 0.9433 | 0.9667 | 0.9687 | 0.8884 | 0.97 | 0.3312 | 0.914 | 0.973 |
| 1d0qA | 0.5347 | 0.7866 | 0.8687 | 0.6782 | 0.7739 | 0.8362 | 0.3159 | 0.8453 | 0.8614 | 0.8116 | 0.8154 | 0.8628 | 0.8226 | 0.856 | 0.9195 | 0.8124 | 0.838 | 0.9288 | 0.8602 | 0.8824 | 0.9387 | 0.358 | 0.8842 | 0.9423 | 0.3733 | 0.8685 | 0.9582 |
| 1d1qA | 0.6877 | 0.8569 | 0.9346 | 0.3161 | 0.9062 | 0.9484 | 0.3254 | 0.8667 | 0.9553 | 0.3305 | 0.9072 | 0.9648 | 0.3542 | 0.9072 | 0.9644 | 0.3404 | 0.9055 | 0.9648 | 0.9563 | 0.9168 | 0.9711 | 0.9496 | 0.9104 | 0.978 | 0.9699 | 0.9178 | 0.9702 |
| 1d4oA | 0.7036 | 0.8765 | 0.9251 | 0.763 | 0.913 | 0.9229 | 0.319 | 0.9324 | 0.9524 | 0.9145 | 0.9393 | 0.9673 | 0.32 | 0.9506 | 0.9689 | 0.957 | 0.9402 | 0.9703 | 0.9724 | 0.9363 | 0.9733 | 0.3335 | 0.9303 | 0.9783 | 0.9787 | 0.9462 | 0.9724 |
| 1dbxA | 0.7199 | 0.761 | 0.8705 | 0.3219 | 0.7953 | 0.9371 | 0.3301 | 0.8196 | 0.9308 | 0.9085 | 0.8345 | 0.9434 | 0.9356 | 0.847 | 0.9516 | 0.335 | 0.8213 | 0.9609 | 0.9362 | 0.8421 | 0.9669 | 0.9364 | 0.8216 | 0.9597 | 0.3356 | 0.8256 | 0.9625 |
| 1dixA | 0.7544 | 0.8661 | 0.8856 | 0.8275 | 0.851 | 0.9187 | 0.8881 | 0.8974 | 0.9524 | 0.8899 | 0.9095 | 0.9625 | 0.3538 | 0.904 | 0.9533 | 0.9578 | 0.8956 | 0.9728 | 0.9531 | 0.8944 | 0.9726 | 0.9599 | 0.9115 | 0.9752 | 0.977 | - | 0.9811 |
| 1dlwA | 0.5852 | 0.8894 | 0.8921 | 0.2576 | 0.834 | 0.9076 | 0.7068 | 0.9063 | 0.9303 | 0.2915 | 0.8676 | 0.9419 | 0.3048 | 0.8678 | 0.9526 | 0.3117 | 0.8926 | 0.9598 | 0.3153 | 0.9066 | 0.9726 | 0.3078 | 0.9151 | 0.9628 | 0.3046 | 0.9181 | 0.9605 |
| 1dmgA | 0.2939 | 0.8207 | 0.8789 | 0.7246 | 0.8205 | 0.9091 | 0.3208 | 0.8504 | 0.9121 | 0.3141 | 0.8376 | 0.9226 | 0.8543 | 0.8412 | 0.9283 | 0.886 | 0.8752 | 0.9305 | 0.9058 | 0.862 | 0.934 | 0.9087 | 0.8743 | 0.9325 | 0.3277 | 0.8583 | 0.9396 |
| 1dqgA | 0.293 | 0.8349 | 0.8948 | 0.3067 | 0.837 | 0.9091 | 0.3085 | 0.8653 | 0.9214 | 0.9266 | 0.8294 | 0.9428 | 0.3123 | 0.8932 | 0.9596 | 0.9505 | 0.9012 | 0.9647 | 0.3135 | 0.8895 | 0.967 | 0.3058 | 0.899 | 0.9638 | 0.3139 | 0.8578 | 0.9656 |

| | | | | | | | | | | | | | | | | | | | | | | | | | | | |
|---|---|---|---|---|---|---|---|---|---|---|---|---|---|---|---|---|---|---|---|---|---|---|---|---|---|---|---|
| 1dsxA | 0.2788 | 0.7102 | 0.7611 | 0.3092 | 0.8231 | 0.8947 | 0.2925 | 0.7656 | 0.879 | 0.307 | 0.8024 | 0.8999 | 0.8572 | 0.8612 | 0.9462 | 0.8684 | 0.9131 | 0.9561 | 0.906 | 0.9156 | 0.9579 | 0.3173 | 0.9122 | 0.9466 | 0.9523 | 0.8785 | 0.9596 |
| 1eazA | 0.609 | 0.7569 | 0.7492 | 0.6173 | 0.7579 | 0.7534 | 0.2923 | 0.6607 | 0.801 | 0.3062 | 0.7997 | 0.8938 | 0.8568 | 0.8299 | 0.9391 | 0.3162 | 0.8919 | 0.9501 | 0.8795 | 0.8454 | 0.957 | 0.3251 | 0.8984 | 0.9517 | 0.3177 | 0.9159 | 0.9507 |
| 1ej0A | 0.3108 | 0.8777 | 0.9159 | 0.7945 | 0.8828 | 0.9207 | 0.8569 | 0.8818 | 0.9317 | 0.3424 | 0.8877 | 0.9583 | 0.3421 | 0.9182 | 0.9703 | 0.343 | 0.9261 | 0.977 | 0.3412 | 0.9246 | 0.9841 | 0.3477 | 0.9255 | 0.9823 | 0.9726 | 0.9279 | 0.9781 |
| 1ej8A | 0.5991 | 0.7445 | 0.7645 | 0.7553 | 0.7856 | 0.8451 | 0.881 | 0.801 | 0.8605 | 0.882 | 0.7991 | 0.9155 | 0.3471 | 0.8081 | 0.9392 | 0.3449 | 0.86 | 0.9557 | 0.3448 | 0.8035 | 0.9659 | 0.9351 | 0.8539 | 0.9533 | 0.9591 | 0.7818 | 0.9524 |
| 1ek0A | 0.7319 | 0.826 | 0.8963 | 0.3046 | 0.9044 | 0.9267 | 0.3287 | 0.8956 | 0.9482 | 0.878 | 0.8589 | 0.943 | 0.9325 | 0.882 | 0.9414 | 0.9486 | 0.8914 | 0.9516 | 0.3322 | 0.8832 | 0.956 | 0.9475 | 0.917 | 0.9711 | 0.337 | 0.9154 | 0.961 |
| 1f6bA | 0.7372 | 0.7916 | 0.8816 | 0.7482 | 0.8205 | 0.892 | 0.807 | 0.8281 | 0.8924 | 0.3072 | 0.8382 | 0.9178 | 0.3064 | 0.8523 | 0.9334 | 0.9228 | 0.8545 | 0.9412 | 0.9514 | 0.874 | 0.9518 | 0.3148 | 0.8583 | 0.9354 | 0.3112 | 0.8707 | 0.943 |
| 1fcyA | 0.3343 | 0.9352 | 0.9521 | 0.8219 | 0.9384 | 0.9557 | 0.8747 | 0.9396 | 0.9606 | 0.3607 | 0.9433 | 0.9698 | 0.3674 | 0.9504 | 0.9791 | 0.3714 | 0.9389 | 0.9814 | 0.9776 | 0.9498 | 0.9779 | 0.3706 | 0.964 | 0.9859 | 0.3698 | 0.9539 | 0.9853 |
| 1fk5A | 0.5442 | 0.817 | 0.8829 | 0.3049 | 0.8314 | 0.8893 | 0.7097 | 0.7987 | 0.9206 | 0.3089 | 0.7857 | 0.921 | 0.8854 | 0.8322 | 0.9225 | 0.9067 | 0.8104 | 0.9271 | 0.3232 | 0.8337 | 0.9336 | 0.3193 | 0.8424 | 0.9397 | 0.3282 | 0.8599 | 0.9448 |
| 1fl0A | 0.8219 | 0.8483 | 0.8898 | 0.7979 | 0.8609 | 0.9401 | 0.8634 | 0.8507 | 0.9546 | 0.9189 | 0.8786 | 0.9488 | 0.3312 | 0.9046 | 0.9629 | 0.9415 | 0.895 | 0.9674 | 0.9641 | 0.8739 | 0.9714 | 0.327 | 0.8739 | 0.9744 | 0.9715 | 0.8916 | 0.9759 |
| 1fnaA | 0.6113 | 0.7257 | 0.6968 | 0.5883 | 0.7119 | 0.768 | 0.2863 | 0.7069 | 0.7937 | 0.3002 | 0.8032 | 0.8677 | 0.7794 | 0.827 | 0.9108 | 0.8466 | 0.7657 | 0.9278 | 0.8704 | 0.7485 | 0.9244 | 0.8673 | 0.6958 | 0.9363 | 0.8874 | 0.847 | 0.943 |
| 1fqtA | 0.3117 | 0.8463 | 0.8634 | 0.7551 | 0.8351 | 0.8782 | 0.328 | 0.8653 | 0.8737 | 0.327 | 0.878 | 0.3279 | 0.3418 | 0.8324 | 0.9562 | 0.8974 | 0.8668 | 0.9602 | 0.3406 | 0.8685 | 0.9643 | 0.9369 | 0.8745 | 0.9703 | 0.9531 | 0.8669 | 0.9689 |
| 1fvgA | 0.6974 | 0.868 | 0.9271 | 0.3301 | 0.8621 | 0.9356 | 0.3451 | 0.8929 | 0.9435 | 0.8729 | 0.8952 | 0.9539 | 0.3458 | 0.9119 | 0.9571 | 0.3491 | 0.9232 | 0.974 | 0.3471 | 0.93 | 0.976 | 0.3427 | 0.9094 | 0.9781 | 0.9728 | 0.9286 | 0.9791 |
| 1fvkA | 0.6634 | 0.8883 | 0.9132 | 0.7258 | 0.8963 | 0.9152 | 0.3327 | 0.8857 | 0.9342 | 0.8487 | 0.9114 | 0.9579 | 0.8871 | 0.8779 | 0.965 | 0.9302 | 0.9088 | 0.9631 | 0.961 | 0.8747 | 0.9718 | 0.9508 | 0.9282 | 0.9692 | 0.3614 | 0.9308 | 0.9765 |
| 1fx2A | 0.4305 | 0.6404 | 0.8108 | 0.5364 | 0.6708 | 0.6874 | 0.4568 | 0.7184 | 0.7528 | 0.621 | 0.5949 | 0.7408 | 0.3468 | 0.5992 | 0.8282 | 0.6866 | 0.8063 | 0.829 | 0.7154 | 0.6651 | 0.7764 | 0.8181 | 0.7809 | 0.897 | 0.8447 | 0.8206 | 0.9245 |
| 1g2rA | 0.2515 | 0.8432 | 0.8898 | 0.2777 | 0.8119 | 0.8779 | 0.273 | 0.8397 | 0.8812 | 0.2834 | 0.8514 | 0.9044 | 0.8493 | 0.8332 | 0.915 | 0.8493 | 0.8484 | 0.9398 | 0.8638 | 0.8831 | 0.9502 | 0.3287 | 0.9002 | 0.9483 | 0.9193 | 0.8679 | 0.9223 |
| 1g9oA | 0.5924 | 0.6924 | 0.7367 | 0.6021 | 0.6924 | 0.7419 | 0.711 | 0.7457 | 0.8109 | 0.7308 | 0.8269 | 0.8551 | 0.2892 | 0.7995 | 0.8875 | 0.8652 | 0.8184 | 0.9132 | 0.8717 | 0.7797 | 0.9043 | 0.8593 | 0.8036 | 0.9054 | 0.3113 | 0.8094 | 0.9193 |
| 1gbsA | 0.3014 | 0.8708 | 0.9144 | 0.3217 | 0.8882 | 0.9304 | 0.3223 | 0.8886 | 0.9367 | 0.328 | 0.8952 | 0.9597 | 0.9351 | 0.9039 | 0.9547 | 0.3262 | 0.9122 | 0.9544 | 0.9769 | 0.9219 | 0.9626 | 0.3247 | 0.903 | 0.9672 | 0.321 | 0.8913 | 0.9691 |
| 1gmiA | 0.708 | 0.7369 | 0.7947 | 0.7047 | 0.7572 | 0.753 | 0.7908 | 0.7538 | 0.8436 | 0.8179 | 0.7933 | 0.8716 | 0.8631 | 0.8211 | 0.9036 | 0.3263 | 0.7603 | 0.9269 | 0.8981 | 0.8034 | 0.9437 | 0.9163 | 0.8405 | 0.9297 | 0.3472 | 0.8167 | 0.9589 |
| 1gmxA | 0.6057 | 0.7684 | 0.8102 | 0.6367 | 0.7884 | 0.8563 | 0.2872 | 0.889 | 0.8956 | 0.3098 | 0.8665 | 0.9332 | 0.911 | 0.8848 | 0.9414 | 0.916 | 0.8754 | 0.9395 | 0.8901 | 0.8702 | 0.9442 | 0.9198 | 0.893 | 0.9474 | 0.3202 | 0.8828 | 0.9605 |
| 1guuA | 0.2271 | 0.6727 | 0.7671 | 0.4597 | 0.7984 | 0.8317 | 0.234 | 0.8268 | 0.8402 | 0.252 | 0.7452 | 0.8597 | 0.7305 | 0.8034 | 0.8527 | 0.7192 | 0.8759 | 0.881 | 0.8034 | 0.8444 | 0.9027 | 0.8076 | 0.8165 | 0.9191 | 0.8513 | 0.912 | 0.904 |
| 1gz2A | 0.6919 | 0.8144 | 0.819 | 0.7923 | 0.7955 | 0.8858 | 0.3183 | 0.8311 | 0.8951 | 0.8658 | 0.7843 | 0.9367 | 0.8881 | 0.8356 | 0.9518 | 0.8775 | 0.853 | 0.9494 | 0.3302 | 0.8619 | 0.9498 | 0.9381 | 0.8503 | 0.9587 | 0.3275 | 0.7929 | 0.9659 |
| 1gzcA | 0.3204 | 0.8652 | 0.8485 | 0.8264 | 0.8902 | 0.8969 | 0.9357 | 0.8897 | 0.934 | 0.3426 | 0.8993 | 0.9624 | 0.9664 | 0.897 | 0.9685 | 0.9608 | 0.9006 | 0.9782 | 0.3443 | 0.91 | 0.9819 | 0.9696 | 0.9145 | 0.9834 | 0.987 | 0.9235 | 0.9859 |
| 1h0pA | 0.8563 | 0.8673 | 0.9247 | 0.8792 | 0.853 | 0.9484 | 0.3108 | 0.8638 | 0.943 | 0.3228 | 0.8834 | 0.9648 | 0.3196 | 0.8806 | 0.9768 | 0.9583 | 0.8708 | 0.98 | 0.9816 | 0.8942 | 0.981 | 0.9619 | 0.8799 | 0.9792 | 0.319 | 0.8705 | 0.9809 |
| 1h2eA | 0.3303 | 0.9102 | 0.935 | 0.3439 | 0.8598 | 0.9427 | 0.8264 | 0.9171 | 0.9502 | 0.896 | 0.9229 | 0.9621 | 0.9484 | 0.9106 | 0.9784 | 0.9591 | 0.909 | 0.9798 | 0.9603 | 0.9464 | 0.9825 | 0.3645 | 0.9515 | 0.9805 | 0.9822 | 0.9473 | 0.9813 |
| 1h4xA | 0.5257 | 0.7644 | 0.8681 | 0.6896 | 0.7852 | 0.9187 | 0.2934 | 0.7743 | 0.923 | 0.304 | 0.8021 | 0.9468 | 0.314 | 0.7892 | 0.9526 | 0.9308 | 0.7967 | 0.9596 | 0.3198 | 0.8686 | 0.9654 | 0.3231 | 0.8754 | 0.9708 | 0.9635 | 0.872 | 0.9727 |
| 1h98A | 0.6987 | 0.8093 | 0.8439 | 0.2753 | 0.8526 | 0.867 | 0.8017 | 0.8049 | 0.8702 | 0.292 | 0.8156 | 0.9055 | 0.2882 | 0.8251 | 0.93 | 0.9156 | 0.8907 | 0.9415 | 0.921 | 0.83 | 0.9319 | 0.2944 | 0.8649 | 0.948 | 0.9301 | 0.8565 | 0.9352 |
| 1hdoA | 0.3156 | 0.9103 | 0.9401 | 0.3169 | 0.914 | 0.9505 | 0.861 | 0.9317 | 0.9598 | 0.3255 | 0.9063 | 0.9647 | 0.3257 | 0.9509 | 0.9783 | 0.3226 | 0.9405 | 0.9831 | 0.3235 | 0.9531 | 0.979 | 0.3277 | 0.9551 | 0.9781 | 0.9826 | 0.9523 | 0.9842 |
| 1hfcA | 0.3129 | 0.8602 | 0.92 | 0.3087 | 0.8655 | 0.9427 | 0.3028 | 0.8265 | 0.9457 | 0.3174 | 0.8984 | 0.9642 | 0.928 | 0.9055 | 0.9655 | 0.9414 | 0.9103 | 0.9779 | 0.3207 | 0.9126 | 0.9707 | 0.324 | 0.9052 | 0.9702 | 0.3203 | 0.9018 | 0.9752 |
| 1hh8A | 0.3501 | 0.9117 | 0.8321 | 0.3587 | 0.9146 | 0.8744 | 0.3576 | 0.9378 | 0.8248 | 0.3668 | 0.9212 | 0.8748 | 0.3882 | 0.9171 | 0.9431 | 0.4022 | 0.9543 | 0.9582 | 0.3995 | 0.9293 | 0.9706 | 0.399 | 0.9422 | 0.9625 | 0.3959 | 0.9176 | 0.9798 |
| 1htwA | 0.6454 | 0.8699 | 0.8952 | 0.7743 | 0.913 | 0.9291 | 0.2852 | 0.923 | 0.9387 | 0.8495 | 0.9323 | 0.9481 | 0.3141 | 0.9166 | 0.9549 | 0.3197 | 0.9187 | 0.969 | 0.3177 | 0.9493 | 0.9724 | 0.9603 | 0.9469 | 0.9722 | 0.9672 | 0.959 | 0.9779 |
| 1hxnA | 0.3231 | 0.8598 | 0.8291 | 0.3465 | 0.8587 | 0.8763 | 0.839 | 0.8593 | 0.9136 | 0.3415 | 0.8852 | 0.9457 | 0.9323 | 0.8829 | 0.9646 | 0.9355 | 0.8918 | 0.9675 | 0.9706 | 0.8887 | 0.9727 | 0.9642 | 0.9032 | 0.9755 | 0.348 | 0.8932 | 0.9774 |
| 1i1jA | 0.5708 | 0.6794 | 0.5241 | 0.6748 | 0.7039 | 0.5207 | 0.3149 | 0.7464 | 0.5842 | 0.8065 | 0.7476 | 0.3498 | 0.8749 | 0.7712 | 0.921 | 0.3211 | 0.7975 | 0.9034 | 0.8892 | 0.8089 | 0.9207 | 0.3334 | 0.8174 | 0.9241 | 0.3331 | 0.7941 | 0.3249 |
| 1i1nA | 0.8203 | 0.9002 | 0.9457 | 0.8846 | 0.9273 | 0.9517 | 0.3132 | 0.9235 | 0.9645 | 0.321 | 0.937 | 0.9698 | 0.3216 | 0.9183 | 0.9744 | 0.323 | 0.9398 | 0.9811 | 0.9713 | 0.9217 | 0.9851 | 0.3223 | 0.9392 | 0.9847 | 0.3212 | 0.9255 | 0.9873 |
| 1i4jA | 0.6563 | 0.8177 | 0.8429 | 0.3368 | 0.7316 | 0.8557 | 0.6283 | 0.7665 | 0.8739 | 0.3544 | 0.7283 | 0.8639 | 0.3703 | 0.7415 | 0.8747 | 0.8939 | 0.7197 | 0.9208 | 0.8763 | 0.7307 | 0.8949 | 0.863 | 0.7982 | 0.8919 | 0.3602 | 0.7366 | 0.8991 |
| 1i58A | 0.5522 | 0.8415 | 0.8937 | 0.3019 | 0.8562 | 0.888 | 0.6555 | 0.8326 | 0.942 | 0.7681 | 0.9006 | 0.9273 | 0.3499 | 0.909 | 0.9443 | 0.3521 | 0.912 | 0.9641 | 0.9262 | 0.8982 | 0.9561 | 0.3465 | 0.9081 | 0.9764 | 0.3534 | 0.9133 | 0.9589 |
| 1i5gA | 0.2946 | 0.8274 | 0.8977 | 0.2894 | 0.8625 | 0.9221 | 0.782 | 0.8584 | 0.9461 | 0.3106 | 0.9116 | 0.9442 | 0.9411 | 0.8954 | 0.9687 | 0.9541 | 0.9004 | 0.9659 | 0.9642 | 0.899 | 0.9739 | 0.957 | 0.8836 | 0.9756 | 0.3164 | 0.9104 | 0.9731 |
| 1i71A | 0.7079 | 0.7228 | 0.7669 | 0.3361 | 0.6944 | 0.8308 | 0.7302 | 0.6974 | 0.8576 | 0.3388 | 0.7592 | 0.8542 | 0.3551 | 0.7371 | 0.8741 | 0.358 | 0.3145 | 0.9016 | 0.8687 | 0.7471 | 0.8871 | 0.8768 | 0.3413 | 0.9062 | 0.8795 | 0.7412 | 0.9194 |

| | | | | | | | | | | | | | | | | | | | | | | | | | | | |
|---|---|---|---|---|---|---|---|---|---|---|---|---|---|---|---|---|---|---|---|---|---|---|---|---|---|---|---|
| 1ihzA | 0.5862 | 0.8491 | 0.8527 | 0.7691 | 0.8246 | 0.88 | 0.3051 | 0.8594 | 0.9032 | 0.788 | 0.8737 | 0.9341 | 0.3371 | 0.8873 | 0.9464 | 0.3331 | 0.8817 | 0.9434 | 0.3312 | 0.8529 | 0.957 | 0.339 | 0.9048 | 0.9668 | 0.9397 | 0.9056 | 0.9624 |
| 1iibA | 0.3012 | 0.8451 | 0.8882 | 0.2935 | 0.8606 | 0.8861 | 0.2971 | 0.8535 | 0.9044 | 0.3132 | 0.8931 | 0.9297 | 0.9218 | 0.9226 | 0.9406 | 0.9394 | 0.9161 | 0.9415 | 0.9557 | 0.9343 | 0.9369 | 0.9462 | 0.9085 | 0.9567 | 0.338 | 0.8918 | 0.9482 |
| 1im5A | 0.3121 | 0.8819 | 0.921 | 0.7905 | 0.88 | 0.9226 | 0.8752 | 0.8821 | 0.96 | 0.3332 | 0.8912 | 0.9604 | 0.3357 | 0.906 | 0.9711 | 0.9373 | 0.9124 | 0.973 | 0.9595 | 0.883 | 0.973 | 0.954 | 0.9141 | 0.9751 | 0.9802 | 0.8936 | 0.9828 |
| 1iwdA | 0.3535 | 0.8997 | 0.9141 | 0.3535 | 0.8557 | 0.9403 | 0.3622 | 0.895 | 0.9546 | 0.9145 | 0.9009 | 0.9666 | 0.9643 | 0.9109 | 0.9744 | 0.954 | 0.9216 | 0.9831 | 0.3607 | 0.9276 | 0.9824 | 0.9615 | 0.9158 | 0.9838 | 0.984 | 0.931 | 0.9865 |
| 1j3aA | 0.2936 | 0.8238 | 0.902 | 0.3032 | 0.8531 | 0.9096 | 0.7467 | 0.8352 | 0.9373 | 0.3217 | 0.8848 | 0.9356 | 0.8483 | 0.8992 | 0.9549 | 0.8937 | 0.9066 | 0.9595 | 0.3275 | 0.8975 | 0.9639 | 0.3249 | 0.8681 | 0.96 | 0.3358 | 0.9249 | 0.959 |
| 1jbeA | 0.2638 | 0.8458 | 0.8892 | 0.2888 | 0.8806 | 0.8996 | 0.2915 | 0.8769 | 0.9334 | 0.8652 | 0.8774 | 0.953 | 0.9179 | 0.9177 | 0.9561 | 0.9304 | 0.9025 | 0.963 | 0.2982 | 0.9159 | 0.9643 | 0.9499 | 0.9144 | 0.956 | 0.9601 | 0.8908 | 0.9659 |
| 1jbkA | 0.2996 | 0.8766 | 0.8978 | 0.7204 | 0.8915 | 0.9141 | 0.3031 | 0.9011 | 0.9563 | 0.8502 | 0.9056 | 0.9578 | 0.3212 | 0.9309 | 0.9703 | 0.9137 | 0.9388 | 0.9763 | 0.3323 | 0.9245 | 0.9708 | 0.9453 | 0.9109 | 0.9746 | 0.3257 | 0.9238 | 0.9696 |
| 1jfuA | 0.3073 | 0.8531 | 0.9086 | 0.3253 | 0.8805 | 0.9293 | 0.3215 | 0.8836 | 0.9488 | 0.9234 | 0.876 | 0.9567 | 0.9631 | 0.8677 | 0.9606 | 0.9512 | 0.9083 | 0.9655 | 0.9723 | 0.9053 | 0.9722 | 0.9692 | 0.9075 | 0.9725 | 0.334 | 0.9356 | 0.9753 |
| 1jfxA | 0.8399 | 0.8402 | 0.9254 | 0.8025 | 0.8723 | 0.9412 | 0.8508 | 0.8924 | 0.9446 | 0.3306 | 0.8983 | 0.9697 | 0.9517 | 0.9268 | 0.9742 | 0.9563 | 0.9279 | 0.9798 | 0.3353 | 0.9297 | 0.9813 | 0.3343 | 0.9202 | 0.9826 | 0.3336 | 0.9216 | 0.9885 |
| 1jkxA | 0.7627 | 0.9053 | 0.9152 | 0.7384 | 0.8677 | 0.9263 | 0.3206 | 0.8956 | 0.9404 | 0.3205 | 0.9003 | 0.9514 | 0.3231 | 0.9064 | 0.9681 | 0.9363 | 0.9125 | 0.9677 | 0.3285 | 0.9321 | 0.9813 | 0.328 | 0.9395 | 0.9758 | 0.9786 | 0.9387 | 0.9827 |
| 1jl1A | 0.59 | 0.7745 | 0.8534 | 0.6631 | 0.8084 | 0.8963 | 0.7092 | 0.8759 | 0.91 | 0.826 | 0.8976 | 0.9283 | 0.8729 | 0.865 | 0.9482 | 0.9005 | 0.8967 | 0.9575 | 0.379 | 0.9016 | 0.9727 | 0.3725 | 0.9213 | 0.9787 | 0.378 | 0.9263 | 0.973 |
| 1jo0A | 0.3312 | 0.8446 | 0.852 | 0.3095 | 0.8075 | 0.9133 | 0.3186 | 0.8373 | 0.9163 | 0.7759 | 0.8493 | 0.9413 | 0.8802 | 0.8641 | 0.9411 | 0.9016 | 0.8331 | 0.965 | 0.3271 | 0.8704 | 0.9688 | 0.3386 | 0.8625 | 0.9676 | 0.3294 | 0.8602 | 0.9539 |
| 1jo8A | 0.2866 | 0.6531 | 0.7114 | 0.2871 | 0.5461 | 0.266 | 0.7371 | 0.6447 | 0.8278 | 0.2828 | 0.731 | 0.8645 | 0.8349 | 0.7274 | 0.8659 | 0.3087 | 0.7974 | 0.8735 | 0.2889 | 0.7719 | 0.8631 | 0.2937 | 0.7976 | 0.8732 | 0.8648 | 0.7316 | 0.9045 |
| 1josA | 0.2847 | 0.7934 | 0.8327 | 0.2852 | 0.8316 | 0.8396 | 0.6635 | 0.83 | 0.8619 | 0.2958 | 0.8515 | 0.8562 | 0.85 | 0.8082 | 0.8822 | 0.3018 | 0.859 | 0.9138 | 0.8913 | 0.8472 | 0.9222 | 0.897 | 0.8508 | 0.9238 | 0.9305 | 0.866 | 0.94 |
| 1jvwA | 0.7399 | 0.8665 | 0.8508 | 0.7487 | 0.7791 | 0.8791 | 0.3491 | 0.8304 | 0.8488 | 0.3515 | 0.8897 | 0.9407 | 0.3781 | 0.8899 | 0.9303 | 0.8848 | 0.8811 | 0.9536 | 0.9065 | 0.872 | 0.9545 | 0.3765 | 0.8975 | 0.9502 | 0.9324 | 0.8599 | 0.9704 |
| 1jwqA | 0.7892 | 0.923 | 0.9601 | 0.8577 | 0.9125 | 0.9629 | 0.3388 | 0.935 | 0.9631 | 0.8927 | 0.9358 | 0.9646 | 0.9444 | 0.9419 | 0.9654 | 0.9581 | 0.9461 | 0.98 | 0.334 | 0.9507 | 0.9785 | 0.9644 | 0.9469 | 0.9801 | 0.9795 | 0.9453 | 0.9822 |
| 1jyhA | 0.298 | 0.6381 | 0.5963 | 0.5041 | 0.6571 | 0.6437 | 0.2843 | 0.7412 | 0.6619 | 0.6531 | 0.7577 | 0.7786 | 0.7915 | 0.8089 | 0.9112 | 0.8463 | 0.8709 | 0.9434 | 0.3215 | 0.8423 | 0.9673 | 0.9131 | 0.8949 | 0.9619 | 0.9423 | 0.8422 | 0.9787 |
| 1k6kA | 0.2998 | 0.8719 | 0.9048 | 0.6068 | 0.8809 | 0.9159 | 0.75 | 0.9025 | 0.9419 | 0.7851 | 0.9304 | 0.9372 | 0.8298 | 0.9427 | 0.9651 | 0.347 | 0.9381 | 0.9621 | 0.9456 | 0.9458 | 0.9712 | 0.3391 | 0.9426 | 0.9772 | 0.3491 | 0.9206 | 0.9754 |
| 1k7cA | 0.3233 | 0.9151 | 0.9259 | 0.8258 | 0.9143 | 0.9447 | 0.8899 | 0.9276 | 0.9518 | 0.3389 | 0.9515 | 0.9696 | 0.3374 | 0.9344 | 0.9713 | 0.3408 | 0.9433 | 0.979 | 0.3402 | 0.951 | 0.9805 | 0.9589 | 0.9482 | 0.9789 | 0.3392 | 0.9604 | 0.978 |
| 1k7jA | 0.316 | 0.8783 | 0.903 | 0.8548 | 0.8857 | 0.9442 | 0.9179 | 0.9182 | 0.9627 | 0.9368 | 0.9122 | 0.9725 | 0.3382 | 0.9289 | 0.9824 | 0.9662 | 0.922 | 0.9811 | 0.9828 | 0.9405 | 0.9864 | 0.9703 | 0.9398 | 0.9849 | 0.34 | 0.9428 | 0.9879 |
| 1kidA | 0.3005 | 0.8731 | 0.8924 | 0.7642 | 0.8716 | 0.9012 | 0.3127 | 0.8341 | 0.9244 | 0.895 | 0.9034 | 0.9415 | 0.336 | 0.8888 | 0.9543 | 0.3349 | 0.9215 | 0.9609 | 0.3194 | 0.9168 | 0.9672 | 0.3188 | 0.9136 | 0.9658 | 0.3277 | 0.9177 | 0.963 |
| 1kq6A | 0.5665 | 0.7554 | 0.8069 | 0.2867 | 0.8143 | 0.8203 | 0.3026 | 0.7918 | 0.86 | 0.632 | 0.8202 | 0.8955 | 0.8703 | 0.8385 | 0.9075 | 0.8449 | 0.8396 | 0.9243 | 0.3289 | 0.8211 | 0.9439 | 0.3356 | 0.8215 | 0.9445 | 0.8547 | 0.864 | 0.9478 |
| 1kqrA | 0.3419 | 0.8414 | 0.8534 | 0.7765 | 0.7713 | 0.8599 | 0.3595 | 0.8584 | 0.9167 | 0.3692 | 0.8436 | 0.939 | 0.3773 | 0.8918 | 0.9605 | 0.3786 | 0.9045 | 0.9625 | 0.3703 | 0.8637 | 0.9654 | 0.3749 | 0.8937 | 0.9719 | 0.3711 | 0.8908 | 0.9695 |
| 1ktgA | 0.2812 | 0.7963 | 0.8614 | 0.7828 | 0.7738 | 0.8516 | 0.325 | 0.8002 | 0.8612 | 0.3333 | 0.8378 | 0.9111 | 0.3591 | 0.8622 | 0.9305 | 0.3565 | 0.8209 | 0.9309 | 0.3431 | 0.8878 | 0.959 | 0.906 | 0.8909 | 0.9531 | 0.9269 | 0.8589 | 0.9465 |
| 1ku3A | 0.282 | 0.6804 | 0.828 | 0.362 | 0.7305 | 0.866 | 0.2912 | 0.7308 | 0.8269 | 0.4235 | 0.6901 | 0.7873 | 0.341 | 0.8312 | 0.8881 | 0.3562 | 0.8321 | 0.8999 | 0.7133 | 0.7634 | 0.8868 | 0.3469 | 0.7963 | 0.9126 | 0.7328 | 0.7955 | 0.9074 |
| 1kw4A | 0.2412 | 0.7539 | 0.8097 | 0.4429 | 0.696 | 0.7885 | 0.5884 | 0.7716 | 0.8229 | 0.2624 | 0.8118 | 0.8867 | 0.2753 | 0.8029 | 0.8796 | 0.304 | 0.8615 | 0.936 | 0.8976 | 0.8547 | 0.9349 | 0.282 | 0.8949 | 0.9498 | 0.2887 | 0.8372 | 0.9408 |
| 1lm4A | 0.3385 | 0.7347 | 0.8608 | 0.7821 | 0.7614 | 0.8959 | 0.8453 | 0.7868 | 0.8812 | 0.3544 | 0.7534 | 0.8173 | 0.8759 | 0.8076 | 0.8248 | 0.3475 | 0.8061 | 0.8143 | 0.3557 | 0.7996 | 0.8209 | 0.9429 | 0.8183 | 0.8352 | 0.9562 | 0.8072 | 0.827 |
| 1lo7A | 0.4452 | 0.6977 | 0.6983 | 0.5273 | 0.7433 | 0.7654 | 0.3058 | 0.8419 | 0.7736 | 0.3328 | 0.7399 | 0.8896 | 0.338 | 0.7887 | 0.9096 | 0.9209 | 0.85 | 0.9285 | 0.3416 | 0.8565 | 0.9318 | 0.9056 | 0.8542 | 0.9297 | 0.9295 | 0.8281 | 0.9166 |
| 1lpyA | 0.4769 | 0.7621 | 0.8057 | 0.5067 | 0.7613 | 0.8794 | 0.3244 | 0.8451 | 0.9365 | 0.695 | 0.8448 | 0.9438 | 0.9238 | 0.8396 | 0.9612 | 0.9215 | 0.8858 | 0.9676 | 0.9315 | 0.8338 | 0.963 | 0.9011 | 0.8316 | 0.9654 | 0.332 | 0.8782 | 0.9693 |
| 1m4jA | 0.3123 | 0.8754 | 0.9193 | 0.7416 | 0.8234 | 0.9256 | 0.33 | 0.8626 | 0.9191 | 0.3353 | 0.8659 | 0.9318 | 0.3431 | 0.8729 | 0.9519 | 0.9352 | 0.9095 | 0.9679 | 0.3462 | 0.8576 | 0.9657 | 0.3477 | 0.8852 | 0.9563 | 0.352 | 0.8875 | 0.9502 |
| 1m8aA | 0.5365 | 0.6745 | 0.7156 | 0.2983 | 0.7366 | 0.7739 | 0.2718 | 0.7651 | 0.8325 | 0.2871 | 0.6954 | 0.8374 | 0.3064 | 0.7839 | 0.8507 | 0.7996 | 0.6653 | 0.8723 | 0.8282 | 0.711 | 0.9257 | 0.7866 | 0.7754 | 0.8932 | 0.8552 | 0.7786 | 0.917 |
| 1mk0A | 0.5954 | 0.8148 | 0.8663 | 0.6709 | 0.8136 | 0.8945 | 0.2948 | 0.8383 | 0.9253 | 0.7719 | 0.8476 | 0.9517 | 0.3258 | 0.8564 | 0.9598 | 0.332 | 0.8732 | 0.9463 | 0.9296 | 0.8839 | 0.9618 | 0.3367 | 0.8752 | 0.9536 | 0.9408 | 0.8941 | 0.959 |
| 1mugA | 0.8158 | 0.8604 | 0.8962 | 0.3031 | 0.8754 | 0.9417 | 0.3086 | 0.8654 | 0.9496 | 0.317 | 0.9045 | 0.9603 | 0.9614 | 0.8973 | 0.9618 | 0.9586 | 0.9174 | 0.9686 | 0.3138 | 0.9181 | 0.9752 | 0.3165 | 0.9247 | 0.9768 | 0.9839 | 0.9254 | 0.976 |
| 1nb9A | 0.6768 | 0.7793 | 0.8545 | 0.3099 | 0.8144 | 0.836 | 0.331 | 0.817 | 0.8631 | 0.3166 | 0.8404 | 0.9186 | 0.8546 | 0.841 | 0.9208 | 0.8778 | 0.8703 | 0.9396 | 0.3305 | 0.8711 | 0.9478 | 0.9319 | 0.8721 | 0.961 | 0.9311 | 0.8737 | 0.9626 |
| 1ne2A | 0.3109 | 0.7826 | 0.8918 | 0.3196 | 0.8556 | 0.9189 | 0.8358 | 0.8252 | 0.8993 | 0.9018 | 0.8292 | 0.8941 | 0.3204 | 0.8577 | 0.9141 | 0.3284 | 0.8618 | 0.9212 | 0.945 | 0.8731 | 0.9329 | 0.9386 | 0.8721 | 0.9496 | 0.3295 | 0.8453 | 0.9377 |
| 1npsA | 0.7943 | 0.7495 | 0.8492 | 0.7831 | 0.7736 | 0.865 | 0.8761 | 0.8126 | 0.8926 | 0.873 | 0.8466 | 0.9315 | 0.905 | 0.8425 | 0.9393 | 0.9101 | 0.8566 | 0.9321 | 0.9263 | 0.8116 | 0.9413 | 0.3148 | 0.8677 | 0.9437 | 0.942 | 0.8195 | 0.9349 |
| 1nrvA | 0.586 | 0.7649 | 0.8045 | 0.6473 | 0.8126 | 0.8611 | 0.7765 | 0.8281 | 0.9259 | 0.3233 | 0.8957 | 0.9282 | 0.3393 | 0.9049 | 0.9541 | 0.9074 | 0.8829 | 0.955 | 0.934 | 0.9206 | 0.9547 | 0.9411 | 0.8947 | 0.9551 | 0.9609 | 0.9078 | 0.956 |

| | | | | | | | | | | | | | | | | | | | | | | | | | | | |
|---|---|---|---|---|---|---|---|---|---|---|---|---|---|---|---|---|---|---|---|---|---|---|---|---|---|---|---|
| 1ny1A | 0.7565 | 0.8943 | 0.9374 | 0.3275 | 0.9168 | 0.9526 | 0.3286 | 0.9041 | 0.9661 | 0.8965 | 0.9622 | 0.9808 | 0.3423 | 0.9468 | 0.984 | 0.9595 | 0.9496 | 0.9855 | 0.9735 | 0.9434 | 0.988 | 0.9731 | 0.9516 | 0.9842 | 0.9859 | 0.9417 | 0.9893 |
| 1o1zA | 0.7278 | 0.8606 | 0.9363 | 0.7773 | 0.8842 | 0.9267 | 0.887 | 0.901 | 0.9467 | 0.8988 | 0.9229 | 0.972 | 0.9346 | 0.9187 | 0.9753 | 0.9437 | 0.9382 | 0.9765 | 0.977 | 0.9371 | 0.9814 | 0.9536 | 0.9389 | 0.9841 | 0.3759 | 0.9433 | 0.982 |
| 1p90A | 0.3143 | 0.7535 | 0.777 | 0.3151 | 0.7387 | 0.7809 | 0.7045 | 0.8215 | 0.8474 | 0.3284 | 0.8862 | 0.9178 | 0.9023 | 0.8852 | 0.9611 | 0.3343 | 0.9347 | 0.9746 | 0.9532 | 0.9229 | 0.9711 | 0.9302 | 0.9102 | 0.9594 | 0.3441 | 0.9125 | 0.9721 |
| 1pchA | 0.2759 | 0.9122 | 0.923 | 0.7902 | 0.86 | 0.9126 | 0.8132 | 0.9104 | 0.9313 | 0.3077 | 0.875 | 0.9222 | 0.9199 | 0.9372 | 0.9638 | 0.3114 | 0.8961 | 0.9647 | 0.946 | 0.9132 | 0.9608 | 0.9532 | 0.9309 | 0.9571 | 0.9561 | 0.9302 | 0.9676 |
| 1pkoA | 0.5722 | 0.7137 | 0.7933 | 0.3091 | 0.7094 | 0.8431 | 0.323 | 0.6957 | 0.8551 | 0.3318 | 0.7074 | 0.888 | 0.3254 | 0.7385 | 0.886 | 0.3463 | 0.3329 | 0.901 | 0.3357 | 0.7181 | 0.9048 | 0.8989 | 0.7011 | 0.901 | 0.9193 | 0.7557 | 0.9164 |
| 1qf9A | 0.5329 | 0.8603 | 0.8912 | 0.5947 | 0.8781 | 0.8959 | 0.3108 | 0.9094 | 0.915 | 0.8012 | 0.8941 | 0.9654 | 0.3292 | 0.9397 | 0.9714 | 0.9405 | 0.9436 | 0.9813 | 0.9469 | 0.9296 | 0.9765 | 0.9468 | 0.9359 | 0.9775 | 0.3336 | 0.9463 | 0.9748 |
| 1qjpA | 0.6582 | 0.6729 | 0.7203 | 0.6489 | 0.7653 | 0.7431 | 0.6837 | 0.7394 | 0.7603 | 0.3724 | 0.7726 | 0.8721 | 0.3696 | 0.764 | 0.8645 | 0.8909 | 0.793 | 0.8786 | 0.875 | 0.8108 | 0.8754 | 0.88 | 0.8097 | 0.8721 | 0.8777 | 0.7962 | 0.8757 |
| 1ql0A | 0.3013 | 0.8531 | 0.9323 | 0.3259 | 0.8926 | 0.948 | 0.9111 | 0.9196 | 0.9562 | 0.9254 | 0.9189 | 0.9735 | 0.3262 | 0.9164 | 0.9737 | 0.3315 | 0.9286 | 0.9812 | 0.3281 | 0.9136 | 0.9773 | 0.9683 | 0.9 | 0.9815 | 0.3317 | 0.9174 | 0.9858 |
| 1r26A | 0.525 | 0.6604 | 0.7972 | 0.6983 | 0.7742 | 0.8907 | 0.3115 | 0.8516 | 0.9269 | 0.3307 | 0.8865 | 0.9367 | 0.9159 | 0.8574 | 0.9414 | 0.3499 | 0.877 | 0.9522 | 0.3463 | 0.874 | 0.9674 | 0.938 | 0.8991 | 0.9742 | 0.9533 | 0.8877 | 0.9688 |
| 1roaA | 0.2536 | 0.6711 | 0.5503 | 0.2914 | 0.7109 | 0.7628 | 0.3158 | 0.7455 | 0.8646 | 0.6738 | 0.7777 | 0.8712 | 0.8023 | 0.7709 | 0.8981 | 0.8244 | 0.7902 | 0.9227 | 0.8793 | 0.8391 | 0.9412 | 0.3537 | 0.7914 | 0.9336 | 0.3626 | 0.8229 | 0.9443 |
| 1rw1A | 0.2879 | 0.8361 | 0.8879 | 0.2927 | 0.8522 | 0.8883 | 0.2959 | 0.8776 | 0.9057 | 0.3135 | 0.8625 | 0.9318 | 0.825 | 0.8668 | 0.9667 | 0.3377 | 0.9092 | 0.9583 | 0.9295 | 0.8381 | 0.9549 | 0.9242 | 0.906 | 0.9541 | 0.9054 | 0.8862 | 0.9605 |
| 1rw7A | 0.802 | 0.9079 | 0.9348 | 0.851 | 0.92 | 0.9523 | 0.8844 | 0.9254 | 0.9592 | 0.9133 | 0.9426 | 0.9687 | 0.9653 | 0.9422 | 0.9806 | 0.9623 | 0.9319 | 0.9828 | 0.3356 | 0.9399 | 0.9835 | 0.9689 | 0.9391 | 0.9798 | 0.9822 | 0.9482 | 0.9846 |
| 1rybA | 0.757 | 0.8762 | 0.9158 | 0.3114 | 0.9123 | 0.9312 | 0.3221 | 0.9093 | 0.9334 | 0.9102 | 0.9275 | 0.9479 | 0.9344 | 0.9236 | 0.9506 | 0.3287 | 0.9146 | 0.9642 | 0.9537 | 0.9262 | 0.9599 | 0.9542 | 0.9308 | 0.9704 | 0.9592 | 0.9397 | 0.9693 |
| 1smxA | 0.6009 | 0.6716 | 0.723 | 0.2669 | 0.7042 | 0.7495 | 0.2858 | 0.6929 | 0.7929 | 0.295 | 0.7241 | 0.8375 | 0.3154 | 0.7865 | 0.8903 | 0.7946 | 0.791 | 0.8939 | 0.8107 | 0.7018 | 0.8878 | 0.772 | 0.7471 | 0.8917 | 0.8593 | 0.7278 | 0.8968 |
| 1svyA | 0.2914 | 0.8628 | 0.8256 | 0.6033 | 0.8718 | 0.8714 | 0.3112 | 0.8404 | 0.8769 | 0.8488 | 0.903 | 0.9128 | 0.9112 | 0.8868 | 0.9432 | 0.3547 | 0.8924 | 0.95 | 0.9166 | 0.9013 | 0.9659 | 0.8998 | 0.922 | 0.9476 | 0.357 | 0.8931 | 0.9614 |
| 1t8kA | 0.6391 | 0.8622 | 0.9097 | 0.282 | 0.8732 | 0.8945 | 0.7439 | 0.8844 | 0.9258 | 0.7747 | 0.9162 | 0.9153 | 0.8822 | 0.8603 | 0.956 | 0.3108 | 0.9081 | 0.9611 | 0.3004 | 0.9066 | 0.9555 | 0.2945 | 0.9306 | 0.9564 | 0.9407 | 0.9191 | 0.9583 |
| 1tifA | 0.2657 | 0.541 | 0.7439 | 0.2644 | 0.5747 | 0.647 | 0.5799 | 0.6696 | 0.7602 | 0.6384 | 0.7264 | 0.7615 | 0.7226 | 0.6464 | 0.8075 | 0.7209 | 0.6944 | 0.7806 | 0.3295 | 0.6515 | 0.7546 | 0.7889 | 0.7317 | 0.8214 | 0.3463 | 0.8178 | 0.8501 |
| 1tqgA | 0.3505 | 0.8971 | 0.9193 | 0.4725 | 0.9115 | 0.9237 | 0.2883 | 0.9457 | 0.9393 | 0.3323 | 0.9623 | 0.9628 | 0.9311 | 0.9084 | 0.9674 | 0.3604 | 0.959 | 0.9728 | 0.3569 | 0.9485 | 0.9684 | 0.3602 | 0.9481 | 0.9606 | 0.944 | 0.9619 | 0.9695 |
| 1tqhA | 0.7313 | 0.9242 | 0.9394 | 0.8079 | 0.8912 | 0.91 | 0.3435 | 0.9222 | 0.9546 | 0.8963 | 0.95 | 0.9804 | 0.9519 | 0.9601 | 0.9766 | 0.3545 | 0.963 | 0.9788 | 0.3492 | 0.9475 | 0.9739 | 0.9529 | 0.9522 | 0.9847 | 0.978 | 0.9479 | 0.9887 |
| 1tzvA | 0.633 | 0.8796 | 0.9393 | 0.2848 | 0.9175 | 0.9455 | 0.6634 | 0.8943 | 0.953 | 0.7906 | 0.91 | 0.9501 | 0.9119 | 0.9336 | 0.9539 | 0.3321 | 0.949 | 0.9772 | 0.3392 | 0.921 | 0.9843 | 0.331 | 0.946 | 0.9754 | 0.9447 | 0.9241 | 0.9709 |
| 1vfyA | 0.4297 | 0.6093 | 0.6611 | 0.4561 | 0.6362 | 0.6914 | 0.2889 | 0.5891 | 0.837 | 0.2981 | 0.5693 | 0.8463 | 0.3086 | 0.5895 | 0.8379 | 0.6933 | 0.6761 | 0.7832 | 0.3209 | 0.6747 | 0.8611 | 0.308 | 0.6719 | 0.8514 | 0.3176 | 0.6925 | 0.8707 |
| 1vhuA | 0.7624 | 0.9166 | 0.9351 | 0.8279 | 0.9232 | 0.9513 | 0.32 | 0.9449 | 0.9699 | 0.9239 | 0.9439 | 0.9692 | 0.3242 | 0.9526 | 0.9806 | 0.3232 | 0.9538 | 0.9815 | 0.3235 | 0.9574 | 0.9827 | 0.9687 | 0.963 | 0.9814 | 0.3188 | 0.9543 | 0.9834 |
| 1vjkA | 0.6128 | 0.817 | 0.7607 | 0.6642 | 0.8693 | 0.8633 | 0.7396 | 0.7858 | 0.8955 | 0.3274 | 0.8789 | 0.9073 | 0.9025 | 0.8707 | 0.9488 | 0.3187 | 0.9192 | 0.9496 | 0.9482 | 0.9044 | 0.9623 | 0.3249 | 0.8845 | 0.9469 | 0.9548 | 0.8968 | 0.9527 |
| 1vmbA | 0.6072 | 0.7352 | 0.7262 | 0.6735 | 0.7646 | 0.7967 | 0.2894 | 0.7649 | 0.8193 | 0.3204 | 0.7865 | 0.8603 | 0.3218 | 0.6898 | 0.9024 | 0.86 | 0.807 | 0.9099 | 0.3301 | 0.7599 | 0.9163 | 0.8616 | 0.7813 | 0.9104 | 0.3541 | 0.8132 | 0.9323 |
| 1vp6A | 0.6077 | 0.8279 | 0.8417 | 0.2991 | 0.819 | 0.8563 | 0.3129 | 0.8383 | 0.9215 | 0.3068 | 0.8595 | 0.9432 | 0.8961 | 0.8963 | 0.9562 | 0.9339 | 0.8901 | 0.966 | 0.9363 | 0.8887 | 0.9754 | 0.3296 | 0.8883 | 0.97 | 0.3303 | 0.9023 | 0.9706 |
| 1w0hA | 0.2834 | 0.8459 | 0.9182 | 0.6884 | 0.8328 | 0.9248 | 0.8069 | 0.8849 | 0.9436 | 0.8325 | 0.9135 | 0.9582 | 0.9399 | 0.9261 | 0.9755 | 0.3536 | 0.8979 | 0.9762 | 0.3608 | 0.9088 | 0.9807 | 0.3539 | 0.8963 | 0.9799 | 0.3541 | 0.9162 | 0.9859 |
| 1whiA | 0.6222 | 0.7686 | 0.8415 | 0.7897 | 0.7321 | 0.8737 | 0.3156 | 0.7808 | 0.8947 | 0.8638 | 0.8188 | 0.9224 | 0.3328 | 0.8264 | 0.9289 | 0.9007 | 0.8565 | 0.9417 | 0.9149 | 0.8441 | 0.9428 | 0.9209 | 0.824 | 0.9491 | 0.3439 | 0.8493 | 0.9423 |
| 1wjxA | 0.5335 | 0.7606 | 0.3196 | 0.2953 | 0.8017 | 0.8251 | 0.2907 | 0.7281 | 0.8991 | 0.8531 | 0.806 | 0.8864 | 0.8463 | 0.8227 | 0.9422 | 0.33 | 0.7963 | 0.9542 | 0.8962 | 0.8287 | 0.9457 | 0.9223 | 0.8417 | 0.9439 | 0.3285 | 0.8341 | 0.9539 |
| 1wkcA | 0.2882 | 0.8941 | 0.8884 | 0.3034 | 0.8718 | 0.9024 | 0.3069 | 0.8556 | 0.9348 | 0.8473 | 0.8884 | 0.9323 | 0.9333 | 0.8901 | 0.9519 | 0.9336 | 0.8888 | 0.9551 | 0.3218 | 0.8911 | 0.9663 | 0.9452 | 0.9033 | 0.9745 | 0.3241 | 0.8934 | 0.9677 |
| 1xdzA | 0.7519 | 0.8507 | 0.8974 | 0.3211 | 0.9279 | 0.9441 | 0.3232 | 0.9267 | 0.9445 | 0.3346 | 0.9206 | 0.9721 | 0.3336 | 0.9477 | 0.9834 | 0.3384 | 0.9515 | 0.9791 | 0.3403 | 0.9389 | 0.9855 | 0.9671 | 0.9473 | 0.9868 | 0.3355 | 0.9207 | 0.9849 |
| 1xffA | 0.8334 | 0.9271 | 0.9377 | 0.3418 | 0.9309 | 0.9603 | 0.3404 | 0.9581 | 0.9707 | 0.3459 | 0.9587 | 0.9765 | 0.3436 | 0.9622 | 0.9846 | 0.3475 | 0.9598 | 0.9856 | 0.344 | 0.9578 | 0.9846 | 0.9716 | 0.955 | 0.9886 | 0.9885 | 0.9525 | 0.9869 |
| 1xkrA | 0.6824 | 0.8724 | 0.8829 | 0.7604 | 0.8889 | 0.9193 | 0.8225 | 0.9118 | 0.9406 | 0.8414 | 0.9262 | 0.9677 | 0.3511 | 0.9516 | 0.9728 | 0.9317 | 0.9554 | 0.9764 | 0.3547 | 0.9396 | 0.981 | 0.9506 | 0.9637 | 0.9796 | 0.979 | 0.9546 | 0.9841 |
| 2arcA | 0.2975 | 0.8249 | 0.9116 | 0.2991 | 0.8654 | 0.9339 | 0.3014 | 0.7851 | 0.932 | 0.3152 | 0.8744 | 0.9517 | 0.9362 | 0.8727 | 0.9672 | 0.9444 | 0.8913 | 0.9693 | 0.3255 | 0.8767 | 0.9674 | 0.9487 | 0.9119 | 0.9752 | 0.3344 | 0.9247 | 0.978 |
| 2cuaA | 0.8022 | 0.7302 | 0.857 | 0.3004 | 0.8122 | 0.8836 | 0.3164 | 0.8001 | 0.8869 | 0.8747 | 0.8004 | 0.9282 | 0.8766 | 0.8118 | 0.9531 | 0.9009 | 0.8004 | 0.9547 | 0.9412 | 0.8803 | 0.9663 | 0.9424 | 0.8054 | 0.9622 | 0.958 | 0.9117 | 0.9618 |
| 2hs1A | 0.2773 | 0.6755 | 0.6549 | 0.6446 | 0.7078 | 0.7194 | 0.6848 | 0.6472 | 0.7285 | 0.7856 | 0.7902 | 0.8861 | 0.322 | 0.8094 | 0.901 | 0.8608 | 0.8174 | 0.9314 | 0.3296 | 0.8275 | 0.9276 | 0.3201 | 0.7694 | 0.9522 | 0.949 | 0.8371 | 0.9523 |
| 2mhrA | 0.3766 | 0.8788 | 0.919 | 0.5819 | 0.8932 | 0.9485 | 0.6326 | 0.8909 | 0.9513 | 0.8059 | 0.8794 | 0.9664 | 0.9406 | 0.9289 | 0.9628 | 0.3376 | 0.9406 | 0.9646 | 0.3468 | 0.952 | 0.972 | 0.3416 | 0.9238 | 0.9776 | 0.962 | 0.9097 | 0.9748 |
| 2phyA | 0.2843 | 0.794 | 0.8309 | 0.2852 | 0.8538 | 0.8969 | 0.2916 | 0.8144 | 0.9027 | 0.8518 | 0.8917 | 0.9361 | 0.9231 | 0.8789 | 0.9438 | 0.9105 | 0.9208 | 0.9658 | 0.9596 | 0.8977 | 0.9663 | 0.954 | 0.8893 | 0.9719 | 0.9636 | 0.921 | 0.9725 |

| | | | | | | | | | | | | | | | | | | | | | | | | | | | |
|---|---|---|---|---|---|---|---|---|---|---|---|---|---|---|---|---|---|---|---|---|---|---|---|---|---|---|---|
| 2tpsA | 0.8409 | 0.9432 | 0.9579 | 0.8715 | 0.9402 | 0.956 | 0.9046 | 0.9413 | 0.9744 | 0.3392 | 0.9456 | 0.9777 | 0.3339 | 0.9654 | 0.9817 | 0.3415 | 0.9586 | 0.9877 | 0.9822 | 0.9429 | 0.9841 | 0.3427 | 0.9655 | 0.9886 | 0.3394 | 0.9688 | 0.988 |
| 2vxnA | 0.3271 | 0.9024 | 0.9104 | 0.3255 | 0.9355 | 0.9559 | 0.8784 | 0.9454 | 0.969 | 0.3321 | 0.9376 | 0.9774 | 0.3271 | 0.956 | 0.981 | 0.9546 | 0.9532 | 0.9847 | 0.972 | 0.9583 | 0.9869 | 0.3349 | 0.9643 | 0.9861 | 0.3288 | 0.9645 | 0.9863 |
| 3borA | 0.2891 | 0.889 | 0.9202 | 0.7854 | 0.8947 | 0.9426 | 0.3169 | 0.9097 | 0.9488 | 0.326 | 0.8985 | 0.9592 | 0.933 | 0.9383 | 0.9615 | 0.9372 | 0.9318 | 0.9583 | 0.9516 | 0.926 | 0.9622 | 0.3386 | 0.9393 | 0.9661 | 0.3353 | 0.9333 | 0.9638 |
| 3dqgA | 0.3311 | 0.561 | 0.6975 | 0.3334 | 0.669 | 0.7386 | 0.3564 | 0.6033 | 0.8093 | 0.3643 | 0.7108 | 0.8346 | 0.3723 | 0.6662 | 0.907 | 0.3941 | 0.649 | 0.9176 | 0.3893 | 0.6359 | 0.916 | 0.3857 | 0.8088 | 0.95 | 0.3898 | 0.793 | 0.9219 |
| 5ptpA | 0.8649 | 0.8669 | 0.931 | 0.3579 | 0.8716 | 0.9544 | 0.3533 | 0.8961 | 0.9618 | 0.9362 | 0.8785 | 0.9718 | 0.3553 | 0.9137 | 0.9788 | 0.9599 | 0.9061 | 0.9807 | 0.355 | 0.8816 | 0.9798 | 0.9665 | 0.9071 | 0.98 | 0.9835 | 0.8977 | 0.9823 |
| Mean | 0.49 | 0.81 | 0.85 | 0.51 | 0.82 | 0.87 | 0.51 | 0.84 | 0.90 | 0.57 | 0.85 | 0.91 | 0.62 | 0.86 | 0.94 | 0.65 | 0.87 | 0.95 | 0.62 | 0.87 | 0.95 | 0.65 | 0.88 | 0.95 | 0.63 | 0.88 | 0.95 |
| Median | 0.52 | 0.83 | 0.87 | 0.46 | 0.84 | 0.90 | 0.33 | 0.85 | 0.92 | 0.37 | 0.87 | 0.94 | 0.76 | 0.88 | 0.95 | 0.81 | 0.89 | 0.96 | 0.38 | 0.88 | 0.97 | 0.82 | 0.90 | 0.96 | 0.38 | 0.89 | 0.97 |

**S10 Table.** Target-by-target stagewise recovery of secondary structure topology on EVfold dataset for true $C_\alpha$–$C_\alpha$ contact maps at 8, 10, and 12Å thresholds.

| Target | 8 Å | | | | | | 10 Å | | | | | | 12 Å | | | | | |
|---|---|---|---|---|---|---|---|---|---|---|---|---|---|---|---|---|---|---|
| | Stage 1 | | Stage 2 | | Stage 3 | | Stage 1 | | Stage 2 | | Stage 3 | | Stage 1 | | Stage 2 | | Stage 3 | |
| | $Q_H$ | $Q_E$ | $Q_H$ | $Q_E$ | $Q_H$ | $Q_E$ | $Q_H$ | $Q_E$ | $Q_H$ | $Q_E$ | $Q_H$ | $Q_E$ | $Q_H$ | $Q_E$ | $Q_H$ | $Q_E$ | $Q_H$ | $Q_E$ |
| 1bkrA | 18.57142857 | | 72.85714286 | | 95.71428571 | | 21.42857143 | | 77.14285714 | | 97.14285714 | | 28.57142857 | | 75.71428571 | | 97.14285714 | |
| 1e6kA | 5.882352941 | 0 | 68.62745098 | 0 | 94.11764706 | 80 | 5.882352941 | 0 | 68.62745098 | 30 | 96.07843137 | 45 | 1.960784314 | 0 | 80.39215686 | 20 | 96.07843137 | 40 |
| 1f21A | 0 | 0 | 63.79310345 | 16.66666667 | 94.82758621 | 68.75 | 32.75862069 | 0 | 86.20689655 | 47.91666667 | 96.55172414 | 62.5 | 13.79310345 | 0 | 86.20689655 | 29.16666667 | 100 | 75 |
| 1g2eA | 14.28571429 | 0 | 71.42857143 | 0 | 95.23809524 | 8 | 0 | 4 | 76.19047619 | 0 | 100 | 24 | 0 | 0 | 90.47619048 | 0 | 100 | 16 |
| 1hzxA | 17.12707182 | 0 | 62.98342541 | 0 | 83.97790055 | 0 | 12.15469613 | 25 | 70.71823204 | 0 | 87.84530387 | 0 | 13.8121547 | 0 | 78.45303867 | 0 | 87.29281768 | 12.5 |
| 1oddA | 15.625 | 0 | 65.625 | 0 | 96.875 | 100 | 0 | 0 | 93.75 | 0 | 96.875 | 71.42857143 | 0 | 0 | 90.625 | 28.57142857 | 100 | 71.42857143 |
| 1r9hA | 0 | 0 | 28.57142857 | 0 | 64.28571429 | 16.66666667 | 28.57142857 | 0 | 57.14285714 | 11.11111111 | 78.57142857 | 91.66666667 | 0 | 2.777777778 | 57.14285714 | 16.66666667 | 78.57142857 | 80.55555556 |
| 1rqmA | 0 | 0 | 43.58974359 | 36 | 89.74358974 | 88 | 28.20512821 | 8 | 66.66666667 | 24 | 92.30769231 | 88 | 30.76923077 | 0 | 71.79487179 | 32 | 89.74358974 | 72 |
| 1wvnA | 22.58064516 | 0 | 70.96774194 | 0 | 100 | 47.05882353 | 0 | 0 | 87.09677419 | 0 | 100 | 70.58823529 | 0 | 11.76470588 | 77.41935484 | 23.52941176 | 100 | 82.35294118 |
| 2hdaA | | 0 | | 0 | | 10.52631579 | | 0 | | 21.05263158 | | 63.15789474 | | 0 | | 0 | | 89.47368421 |
| 2it6A | 11.11111111 | 0 | 70.37037037 | 0 | 85.18518519 | 23.52941176 | 0 | 5.882352941 | 81.48148148 | 23.52941176 | 96.2962963 | 58.82352941 | 0 | 5.882352941 | 48.14814815 | 0 | 100 | 58.82352941 |
| 2o72A | | 0 | | 17.0212766 | | 74.46808511 | | 4.255319149 | | 12.76595745 | | 80.85106383 | | 0 | | 31.91489362 | | 65.95744681 |
| 3tgiE | 0 | 0 | 42.85714286 | 23.68421053 | 100 | 67.10526316 | 0 | 5.263157895 | 57.14285714 | 32.89473684 | 100 | 81.57894737 | 0 | 1.315789474 | 42.85714286 | 48.68421053 | 100 | 76.31578947 |
| 5p21A | 11.29032258 | 0 | 69.35483871 | 30.76923077 | 100 | 66.66666667 | 14.51612903 | 2.564102564 | 85.48387097 | 35.8974359 | 98.38709677 | 58.97435897 | 17.74193548 | 0 | 79.03225806 | 38.46153846 | 100 | 87.17948718 |
| 5ptiA | 12.5 | 0 | 87.5 | 26.66666667 | 100 | 86.66666667 | 0 | 0 | 87.5 | 26.66666667 | 100 | 86.66666667 | 0 | 0 | 62.5 | 13.33333333 | 100 | 86.66666667 |
| Mean | 9.921049729 | 0 | 62.9635354 | 10.77200366 | 92.30500031 | 52.67413567 | 11.03976362 | 3.926066611 | 76.55003235 | 18.988187 | 95.38891004 | 63.08828103 | 8.203741329 | 1.552901863 | 72.36632316 | 20.1662964 | 96.06377881 | 65.30383371 |

**S11 Table.** *Ab initio* folding performance of DConStruct on EVfold dataset using top hybrid interaction maps with tri-level thresholding at increasing xL values (x = 2, 4, 8, 16).

| Target | 2L | 4L | 8L | 16L |
|---|---|---|---|---|
| 1bkrA | 0.8383 | 0.8267 | 0.8257 | 0.7377 |
| 1e6kA | 0.8332 | 0.8516 | 0.8972 | 0.784 |
| 1f21A | 0.7557 | 0.7753 | 0.8162 | 0.7386 |
| 1g2eA | 0.5946 | 0.7795 | 0.7998 | 0.6301 |
| 1hzxA | 0.6165 | 0.715 | 0.7326 | 0.6934 |
| 1oddA | 0.194 | 0.2153 | 0.2592 | 0.2625 |
| 1r9hA | 0.544 | 0.7663 | 0.7929 | 0.6209 |
| 1rqmA | 0.7086 | 0.7168 | 0.7427 | 0.6888 |
| 1wvnA | 0.719 | 0.807 | 0.8566 | 0.5849 |
| 2hdaA | 0.2272 | 0.2188 | 0.2816 | 0.1883 |
| 2it6A | 0.1248 | 0.2079 | 0.2617 | 0.2008 |
| 2o72A | 0.1724 | 0.1568 | 0.1749 | 0.1777 |
| 3tgiE | 0.729 | 0.7821 | 0.8513 | 0.8407 |
| 5p21A | 0.781 | 0.7736 | 0.8141 | 0.7656 |
| 5ptiA | 0.6135 | 0.7171 | 0.7072 | 0.5867 |
| Mean | 0.563453333 | 0.620653333 | 0.654246667 | 0.566713333 |

**S2 Table.** Target-by-target reconstruction performance on 150 soluble proteins for true $C_\beta$–$C_\beta$ contact maps at various thresholds.

| Target | 8 Å | | 8.5 Å | | 9 Å | | 9.5 Å | | 10 Å | | 10.5 Å | | 11 Å | | 11.5 Å | | 12 Å | |
|---|---|---|---|---|---|---|---|---|---|---|---|---|---|---|---|---|---|---|
| | CONFOLD | DConStruct | CONFOLD | DConStruct | CONFOLD | DConStruct | CONFOLD | DConStruct | CONFOLD | DConStruct | CONFOLD | DConStruct | CONFOLD | DConStruct | CONFOLD | DConStruct | CONFOLD | DConStruct |
| 1a3aA | 0.8825 | 0.9127 | 0.9001 | 0.9107 | 0.8827 | 0.9364 | 0.8936 | 0.9391 | 0.8921 | 0.9486 | 0.9143 | 0.9435 | 0.9188 | 0.9511 | 0.9173 | 0.9435 | 0.9242 | 0.946 |
| 1a6mA | 0.9026 | 0.8707 | 0.9105 | 0.8721 | 0.9254 | 0.8829 | 0.9217 | 0.8955 | 0.9369 | 0.9103 | 0.9429 | 0.9302 | 0.9425 | 0.9279 | 0.948 | 0.9291 | 0.9503 | 0.9357 |
| 1a70A | 0.8531 | 0.8698 | 0.8717 | 0.8795 | 0.9137 | 0.8968 | 0.9152 | 0.9043 | 0.8976 | 0.9258 | 0.93 | 0.9155 | 0.9305 | 0.9263 | 0.9268 | 0.9306 | 0.9219 | 0.9311 |
| 1aapA | 0.693 | 0.8124 | 0.6657 | 0.761 | 0.8189 | 0.8427 | 0.8387 | 0.8309 | 0.8094 | 0.7888 | 0.7997 | 0.8315 | 0.8048 | 0.8244 | 0.7985 | 0.8119 | 0.7759 | 0.8055 |
| 1abaA | 0.8352 | 0.8567 | 0.8502 | 0.8844 | 0.8498 | 0.8661 | 0.8454 | 0.8198 | 0.9073 | 0.8571 | 0.8497 | 0.8738 | 0.8959 | 0.8784 | 0.9159 | 0.8988 | 0.8743 | 0.8834 |
| 1ag6A | 0.8397 | 0.8947 | 0.9043 | 0.9085 | 0.8558 | 0.902 | 0.9222 | 0.9183 | 0.9183 | 0.93 | 0.9195 | 0.9124 | 0.9365 | 0.936 | 0.9287 | 0.9309 | 0.9289 | 0.9331 |
| 1aoeA | 0.909 | 0.9163 | 0.8958 | 0.9463 | 0.9211 | 0.9433 | 0.9311 | 0.9378 | 0.919 | 0.9529 | 0.9325 | 0.9577 | 0.8871 | 0.959 | 0.8608 | 0.9574 | 0.9275 | 0.9638 |
| 1atlA | 0.9213 | 0.9436 | 0.9186 | 0.941 | 0.9286 | 0.9514 | 0.9418 | 0.9563 | 0.9415 | 0.9643 | 0.9569 | 0.9637 | 0.9277 | 0.9622 | 0.9429 | 0.9605 | 0.9389 | 0.9676 |
| 1atzA | 0.8048 | 0.7988 | 0.799 | 0.8384 | 0.8083 | 0.8317 | 0.7703 | 0.8761 | 0.8618 | 0.8693 | 0.8812 | 0.9137 | 0.8741 | 0.891 | 0.8497 | 0.9043 | 0.8475 | 0.9135 |
| 1avsA | 0.8387 | 0.887 | 0.8373 | 0.8944 | 0.903 | 0.8558 | 0.874 | 0.8415 | 0.9059 | 0.8775 | 0.8972 | 0.8694 | 0.8921 | 0.8763 | 0.8945 | 0.8446 | 0.913 | 0.8584 |
| 1bdoA | 0.7247 | 0.8398 | 0.7373 | 0.8414 | 0.7361 | 0.8476 | 0.2968 | 0.8686 | 0.794 | 0.9036 | 0.7939 | 0.9021 | 0.7994 | 0.9236 | 0.307 | 0.9097 | 0.2983 | 0.9176 |
| 1bebA | 0.8714 | 0.932 | 0.8815 | 0.9323 | 0.8818 | 0.9413 | 0.862 | 0.9478 | 0.8883 | 0.9563 | 0.9198 | 0.9603 | 0.8996 | 0.9614 | 0.919 | 0.9676 | 0.9225 | 0.9662 |
| 1behA | 0.8967 | 0.9336 | 0.9026 | 0.9345 | 0.8911 | 0.9389 | 0.925 | 0.9524 | 0.9239 | 0.9518 | 0.9226 | 0.9527 | 0.9107 | 0.949 | 0.92 | 0.9522 | 0.9177 | 0.9552 |
| 1bkrA | 0.8926 | 0.9051 | 0.9023 | 0.8959 | 0.9279 | 0.9139 | 0.9191 | 0.9293 | 0.9482 | 0.9343 | 0.9469 | 0.9448 | 0.939 | 0.9408 | 0.9534 | 0.9421 | 0.9587 | 0.9513 |
| 1brfA | 0.7719 | 0.7853 | 0.8071 | 0.7505 | 0.7804 | 0.7778 | 0.8011 | 0.8316 | 0.8255 | 0.8275 | 0.8119 | 0.801 | 0.8104 | 0.8675 | 0.876 | 0.81 | 0.8697 | 0.8352 |
| 1bsgA | 0.9487 | 0.9481 | 0.9457 | 0.9581 | 0.9655 | 0.9615 | 0.9588 | 0.9639 | 0.9721 | 0.9712 | 0.9705 | 0.9725 | 0.9709 | 0.9699 | 0.9754 | 0.9684 | 0.9682 | 0.9684 |
| 1c44A | 0.821 | 0.8417 | 0.8287 | 0.8377 | 0.839 | 0.8259 | 0.88 | 0.8549 | 0.8843 | 0.8691 | 0.883 | 0.8684 | 0.8798 | 0.8671 | 0.9008 | 0.8422 | 0.8947 | 0.875 |
| 1c52A | 0.888 | 0.9165 | 0.8992 | 0.9101 | 0.9305 | 0.9235 | 0.9083 | 0.9352 | 0.919 | 0.9372 | 0.9361 | 0.9394 | 0.9174 | 0.9391 | 0.9425 | 0.9334 | 0.9438 | 0.9358 |
| 1c9oA | 0.7234 | 0.8217 | 0.7186 | 0.7863 | 0.7058 | 0.8184 | 0.7054 | 0.8005 | 0.7713 | 0.8785 | 0.7105 | 0.8435 | 0.7071 | 0.8308 | 0.7228 | 0.841 | 0.7223 | 0.8398 |
| 1cc8A | 0.7367 | 0.8167 | 0.7917 | 0.8441 | 0.8128 | 0.8527 | 0.8201 | 0.839 | 0.8515 | 0.8503 | 0.8125 | 0.8982 | 0.8288 | 0.8926 | 0.8839 | 0.9001 | 0.8533 | 0.9017 |
| 1chdA | 0.9461 | 0.9462 | 0.9398 | 0.952 | 0.9475 | 0.9597 | 0.9645 | 0.9639 | 0.9505 | 0.9595 | 0.9567 | 0.9628 | 0.9595 | 0.9653 | 0.9658 | 0.9594 | 0.9646 | 0.9572 |
| 1cjwA | 0.8221 | 0.9108 | 0.8295 | 0.9155 | 0.8213 | 0.9097 | 0.8559 | 0.9379 | 0.8576 | 0.9327 | 0.8622 | 0.942 | 0.8597 | 0.9393 | 0.8637 | 0.9526 | 0.8458 | 0.9398 |
| 1ckeA | 0.8681 | 0.8322 | 0.8648 | 0.8278 | 0.8981 | 0.8512 | 0.8848 | 0.8739 | 0.8921 | 0.8817 | 0.8949 | 0.8779 | 0.9012 | 0.8992 | 0.883 | 0.8976 | 0.9046 | 0.903 |
| 1ctfA | 0.821 | 0.753 | 0.8399 | 0.7952 | 0.8174 | 0.7875 | 0.7728 | 0.8248 | 0.8236 | 0.8542 | 0.8062 | 0.76 | 0.8233 | 0.7988 | 0.8224 | 0.834 | 0.8008 | 0.8028 |
| 1cxyA | 0.7626 | 0.7989 | 0.7933 | 0.835 | 0.784 | 0.8557 | 0.8008 | 0.881 | 0.8329 | 0.8483 | 0.8404 | 0.8558 | 0.8612 | 0.8566 | 0.8292 | 0.8669 | 0.8355 | 0.9048 |
| 1cznA | 0.9145 | 0.9266 | 0.935 | 0.9311 | 0.9377 | 0.9357 | 0.9454 | 0.9414 | 0.9572 | 0.946 | 0.9591 | 0.9421 | 0.9526 | 0.9512 | 0.961 | 0.9459 | 0.9591 | 0.9482 |
| 1d0qA | 0.8118 | 0.8294 | 0.8476 | 0.8428 | 0.8392 | 0.8304 | 0.8769 | 0.8316 | 0.912 | 0.8952 | 0.8866 | 0.9038 | 0.9352 | 0.906 | 0.8862 | 0.8881 | 0.8938 | 0.9055 |
| 1d1qA | 0.904 | 0.9298 | 0.9274 | 0.9291 | 0.925 | 0.9285 | 0.9315 | 0.9499 | 0.9299 | 0.9367 | 0.9283 | 0.9436 | 0.9412 | 0.9374 | 0.9419 | 0.9403 | 0.9395 | 0.9267 |
| 1d4oA | 0.9366 | 0.9331 | 0.9459 | 0.9304 | 0.9468 | 0.9454 | 0.9553 | 0.9434 | 0.9663 | 0.9551 | 0.9568 | 0.9622 | 0.9557 | 0.9576 | 0.9508 | 0.9495 | 0.9545 | 0.9544 |
| 1dbxA | 0.8338 | 0.9087 | 0.8283 | 0.917 | 0.841 | 0.914 | 0.8589 | 0.932 | 0.8633 | 0.9313 | 0.8848 | 0.942 | 0.8619 | 0.9298 | 0.8699 | 0.9242 | 0.8689 | 0.9357 |
| 1dixA | 0.8994 | 0.9115 | 0.9132 | 0.9166 | 0.8993 | 0.936 | 0.9073 | 0.9363 | 0.9134 | 0.9324 | 0.94 | 0.9396 | 0.9306 | 0.9505 | 0.9188 | 0.9397 | 0.9082 | 0.945 |
| 1dlwA | 0.8739 | 0.8927 | 0.9019 | 0.9036 | 0.9046 | 0.9025 | 0.8994 | 0.915 | 0.9433 | 0.9161 | 0.9214 | 0.9234 | 0.9575 | 0.9509 | 0.9663 | 0.9457 | 0.9388 | 0.9455 |
| 1dmgA | 0.8571 | 0.884 | 0.858 | 0.8787 | 0.8827 | 0.8854 | 0.865 | 0.8961 | 0.8954 | 0.9085 | 0.8687 | 0.8928 | 0.893 | 0.9097 | 0.8785 | 0.9091 | 0.862 | 0.9072 |
| 1dqgA | 0.8828 | 0.9097 | 0.883 | 0.9171 | 0.8839 | 0.9402 | 0.9158 | 0.9351 | 0.9251 | 0.9351 | 0.9136 | 0.9405 | 0.9206 | 0.9412 | 0.9239 | 0.9422 | 0.9063 | 0.9362 |

| | | | | | | | | | | | | | | | | | | |
|---|---|---|---|---|---|---|---|---|---|---|---|---|---|---|---|---|---|---|
| 1dsxA | 0.8932 | 0.8774 | 0.8529 | 0.8425 | 0.8409 | 0.8648 | 0.8343 | 0.8573 | 0.8991 | 0.9119 | 0.9324 | 0.9169 | 0.9355 | 0.9151 | 0.9162 | 0.9206 | 0.9242 | 0.9219 |
| 1eazA | 0.8745 | 0.8819 | 0.8511 | 0.8883 | 0.884 | 0.8941 | 0.9036 | 0.9292 | 0.9305 | 0.9302 | 0.9158 | 0.9417 | 0.9091 | 0.9442 | 0.9102 | 0.9427 | 0.9362 | 0.9499 |
| 1ej0A | 0.9174 | 0.9103 | 0.9279 | 0.9305 | 0.9237 | 0.9411 | 0.9327 | 0.9397 | 0.9336 | 0.9422 | 0.9483 | 0.9558 | 0.9293 | 0.951 | 0.9448 | 0.9469 | 0.9303 | 0.9436 |
| 1ej8A | 0.8259 | 0.8427 | 0.808 | 0.865 | 0.87 | 0.9029 | 0.8684 | 0.9172 | 0.842 | 0.9332 | 0.8422 | 0.9283 | 0.8483 | 0.9348 | 0.8364 | 0.9362 | 0.8529 | 0.9354 |
| 1ek0A | 0.8958 | 0.902 | 0.899 | 0.8932 | 0.8765 | 0.9162 | 0.8919 | 0.9213 | 0.9193 | 0.9262 | 0.9265 | 0.9345 | 0.9189 | 0.941 | 0.9181 | 0.9466 | 0.9168 | 0.9439 |
| 1f6bA | 0.8258 | 0.8725 | 0.8773 | 0.8882 | 0.8563 | 0.8924 | 0.873 | 0.9004 | 0.8736 | 0.9029 | 0.895 | 0.9164 | 0.8968 | 0.9149 | 0.8779 | 0.9134 | 0.8822 | 0.9149 |
| 1fcyA | 0.9379 | 0.942 | 0.9534 | 0.9412 | 0.9577 | 0.9502 | 0.9543 | 0.9569 | 0.9674 | 0.964 | - | 0.9693 | 0.9665 | 0.9683 | 0.9607 | 0.9665 | 0.9677 | 0.9694 |
| 1fk5A | 0.8631 | 0.8569 | 0.8851 | 0.8506 | 0.8824 | 0.88 | 0.8802 | 0.8594 | 0.8763 | 0.8651 | 0.8689 | 0.8614 | 0.857 | 0.879 | 0.8431 | 0.8773 | 0.8873 | 0.8686 |
| 1fl0A | 0.9016 | 0.9142 | 0.8893 | 0.9314 | 0.9246 | 0.9282 | 0.916 | 0.9348 | 0.9156 | 0.9204 | 0.9216 | 0.9436 | 0.9261 | 0.951 | 0.9291 | 0.9561 | 0.9136 | 0.9517 |
| 1fnaA | 0.7947 | 0.806 | 0.8248 | 0.8519 | 0.7714 | 0.8626 | 0.8175 | 0.857 | 0.7885 | 0.8991 | 0.8831 | 0.8903 | 0.8889 | 0.8828 | 0.8857 | 0.8906 | 0.858 | 0.8982 |
| 1fqtA | 0.8724 | 0.8923 | 0.8589 | 0.879 | 0.8749 | 0.8967 | 0.856 | 0.9052 | 0.8823 | 0.9171 | 0.9018 | 0.9103 | 0.921 | 0.939 | 0.8939 | 0.921 | 0.9089 | 0.9308 |
| 1fvgA | 0.8783 | 0.9054 | 0.9096 | 0.95 | 0.9127 | 0.9386 | 0.9336 | 0.9542 | 0.929 | 0.9612 | 0.9339 | 0.9501 | 0.9403 | 0.9578 | 0.9493 | 0.953 | 0.9406 | 0.9392 |
| 1fvkA | 0.8981 | 0.8694 | 0.9067 | 0.8858 | 0.9284 | 0.903 | 0.9453 | 0.9299 | 0.9487 | 0.9468 | 0.9424 | 0.9329 | 0.9281 | 0.9413 | 0.9467 | 0.9443 | 0.9373 | 0.9382 |
| 1fx2A | 0.8005 | 0.7958 | 0.7094 | 0.8429 | 0.8299 | 0.8863 | 0.8237 | 0.8579 | 0.7672 | 0.906 | 0.6632 | 0.8694 | 0.7941 | 0.8339 | 0.8345 | 0.9071 | 0.8631 | 0.9022 |
| 1g2rA | 0.8276 | 0.8223 | 0.8314 | 0.8835 | 0.8475 | 0.8436 | 0.8581 | 0.8991 | 0.8917 | 0.9274 | 0.8935 | 0.9213 | 0.8782 | 0.927 | 0.9162 | 0.9047 | 0.8861 | 0.917 |
| 1g9oA | 0.7579 | 0.7917 | 0.8049 | 0.7975 | 0.8127 | 0.8194 | 0.7853 | 0.8202 | 0.7728 | 0.8738 | 0.8333 | 0.8859 | 0.8048 | 0.8794 | 0.831 | 0.8667 | 0.8233 | 0.8773 |
| 1gbsA | 0.8857 | 0.8953 | 0.9054 | 0.9152 | 0.8927 | 0.9268 | 0.9279 | 0.9161 | 0.9286 | 0.9227 | 0.9317 | 0.9299 | 0.9125 | 0.9266 | 0.9381 | 0.9245 | 0.9395 | 0.9258 |
| 1gmiA | 0.8046 | 0.8416 | 0.81 | 0.8592 | 0.805 | 0.8664 | 0.8273 | 0.8696 | 0.8412 | 0.8943 | 0.8378 | 0.8866 | 0.825 | 0.9058 | 0.88 | 0.9161 | 0.849 | 0.9201 |
| 1gmxA | 0.878 | 0.8715 | 0.883 | 0.8769 | 0.905 | 0.9018 | 0.9057 | 0.9176 | 0.9159 | 0.9167 | 0.9301 | 0.9296 | 0.9241 | 0.9297 | 0.9191 | 0.9228 | 0.9375 | 0.9197 |
| 1guuA | 0.858 | 0.7803 | 0.7863 | 0.7688 | 0.8773 | 0.8395 | 0.8602 | 0.8584 | 0.8326 | 0.8597 | 0.8734 | 0.8848 | 0.8782 | 0.8862 | 0.9018 | 0.9103 | 0.9025 | 0.8976 |
| 1gz2A | 0.8641 | 0.8885 | 0.8503 | 0.897 | 0.8689 | 0.8962 | 0.87 | 0.9124 | 0.8909 | 0.9229 | 0.8951 | 0.936 | 0.8927 | 0.9377 | 0.8908 | 0.9476 | 0.9129 | 0.9376 |
| 1gzcA | 0.9203 | 0.9452 | 0.934 | 0.9403 | 0.9359 | 0.9491 | 0.9381 | 0.9553 | 0.9581 | 0.9616 | 0.9514 | 0.966 | 0.9613 | 0.9734 | 0.9575 | 0.9684 | 0.9628 | 0.9725 |
| 1h0pA | 0.8648 | 0.9376 | 0.8959 | 0.9401 | 0.8998 | 0.9438 | 0.8937 | 0.9546 | 0.9185 | 0.9594 | 0.9167 | 0.9664 | 0.9327 | 0.9474 | 0.9091 | 0.9477 | 0.907 | 0.9506 |
| 1h2eA | 0.9222 | 0.9187 | 0.9252 | 0.942 | 0.9409 | 0.9566 | 0.9534 | 0.959 | 0.9511 | 0.9635 | 0.9538 | 0.9655 | 0.9598 | 0.9697 | 0.9569 | 0.9638 | 0.9414 | 0.9642 |
| 1h4xA | 0.7863 | 0.8921 | 0.7788 | 0.9111 | 0.8369 | 0.915 | 0.8464 | 0.9238 | 0.884 | 0.9373 | 0.8801 | 0.9336 | 0.8673 | 0.9329 | 0.873 | 0.9365 | 0.8815 | 0.9288 |
| 1h98A | 0.8642 | 0.8793 | 0.8288 | 0.8614 | 0.8479 | 0.8802 | 0.8973 | 0.8802 | 0.8585 | 0.9112 | 0.9139 | 0.9209 | 0.8776 | 0.9109 | 0.8983 | 0.9138 | 0.9237 | 0.891 |
| 1hdoA | 0.9428 | 0.9426 | 0.9481 | 0.9499 | 0.9561 | 0.9578 | 0.9557 | 0.9639 | 0.9625 | 0.963 | 0.9666 | 0.9589 | 0.9679 | 0.9607 | 0.9475 | 0.9584 | 0.971 | 0.961 |
| 1hfcA | 0.9003 | 0.9089 | 0.9246 | 0.9159 | 0.9266 | 0.9301 | 0.9357 | 0.9246 | 0.9546 | 0.9393 | 0.9408 | 0.9474 | 0.9329 | 0.952 | 0.9473 | 0.9488 | 0.9532 | 0.9369 |
| 1hh8A | 0.9068 | 0.8301 | 0.9359 | 0.8553 | 0.9371 | 0.8787 | 0.9527 | 0.9187 | 0.9683 | 0.9359 | 0.9613 | 0.9553 | 0.9613 | 0.9464 | 0.9586 | 0.9296 | 0.9699 | 0.9557 |
| 1htwA | 0.9296 | 0.9029 | 0.9365 | 0.9325 | 0.9336 | 0.9337 | 0.9514 | 0.9375 | 0.9637 | 0.9401 | 0.9558 | 0.9516 | 0.9626 | 0.9477 | 0.9658 | 0.9511 | 0.9656 | 0.9451 |
| 1hxnA | 0.9002 | 0.9283 | 0.9142 | 0.9307 | 0.9218 | 0.9393 | 0.9244 | 0.9459 | 0.9342 | 0.9469 | 0.9221 | 0.9458 | 0.9296 | 0.9572 | 0.9361 | 0.9559 | 0.9419 | 0.9582 |
| 1i1jA | 0.7618 | 0.6078 | 0.7795 | 0.6284 | 0.8183 | 0.8796 | 0.829 | 0.8841 | 0.8363 | 0.9031 | 0.8618 | 0.9031 | 0.8554 | 0.3215 | 0.8552 | 0.9089 | 0.8642 | 0.9043 |
| 1i1nA | 0.9289 | 0.9551 | 0.9346 | 0.95 | 0.942 | 0.9579 | 0.9466 | 0.9601 | 0.946 | 0.9623 | 0.9531 | 0.9603 | 0.946 | 0.955 | 0.9582 | 0.9576 | 0.9546 | 0.9634 |
| 1i4jA | 0.8113 | 0.8271 | 0.7931 | 0.8414 | 0.8082 | 0.8591 | 0.8247 | 0.8561 | 0.8257 | 0.865 | 0.8295 | 0.9015 | 0.8356 | 0.9204 | 0.8118 | 0.8946 | 0.7813 | 0.8995 |
| 1i58A | 0.8893 | 0.9029 | 0.9126 | 0.9163 | 0.9247 | 0.9255 | 0.9183 | 0.9316 | 0.9377 | 0.9325 | 0.935 | 0.9453 | 0.9432 | 0.9412 | 0.9454 | 0.9459 | 0.9391 | 0.9467 |
| 1i5gA | 0.8824 | 0.9136 | 0.8861 | 0.9235 | 0.9087 | 0.9324 | 0.923 | 0.9452 | 0.9161 | 0.9514 | 0.925 | 0.9471 | 0.9151 | 0.9485 | 0.9297 | 0.9527 | 0.9365 | 0.9519 |
| 1i71A | 0.7052 | 0.8159 | 0.7613 | 0.8382 | 0.8256 | 0.8667 | 0.7958 | 0.8603 | 0.8459 | 0.8433 | 0.8518 | 0.8628 | 0.8149 | 0.8702 | 0.7887 | 0.8902 | 0.8082 | 0.9065 |

| | | | | | | | | | | | | | | | | | | |
|---|---|---|---|---|---|---|---|---|---|---|---|---|---|---|---|---|---|---|
| 1ihzA | 0.8645 | 0.8811 | 0.8795 | 0.9025 | 0.9085 | 0.9064 | 0.8925 | 0.9092 | 0.9186 | 0.9325 | 0.9219 | 0.9377 | 0.9248 | 0.9478 | 0.9078 | 0.9499 | 0.9186 | 0.9623 |
| 1iibA | 0.8821 | 0.8892 | 0.8823 | 0.8921 | 0.9132 | 0.8987 | 0.8959 | 0.9033 | 0.9398 | 0.8949 | 0.9282 | 0.9108 | 0.9347 | 0.9157 | 0.9241 | 0.9131 | 0.9251 | 0.8867 |
| 1im5A | 0.9095 | 0.9242 | 0.8968 | 0.9348 | 0.9078 | 0.9395 | 0.9342 | 0.9631 | 0.94 | 0.9603 | 0.9452 | 0.956 | 0.9257 | 0.9597 | 0.9537 | 0.9602 | 0.9424 | 0.9539 |
| 1iwdA | 0.9327 | 0.9387 | 0.9303 | 0.9492 | 0.95 | 0.9647 | 0.9336 | 0.9569 | 0.9394 | 0.953 | 0.9402 | 0.9559 | 0.9515 | 0.9556 | 0.9473 | 0.9526 | 0.9339 | 0.9553 |
| 1j3aA | 0.8616 | 0.9049 | 0.8843 | 0.9272 | 0.9023 | 0.9284 | 0.9192 | 0.9441 | 0.9051 | 0.9439 | 0.9158 | 0.9523 | 0.9376 | 0.9475 | 0.9221 | 0.9501 | 0.9457 | 0.9512 |
| 1jbeA | 0.8653 | 0.9042 | 0.9018 | 0.9114 | 0.9196 | 0.9197 | 0.9183 | 0.931 | 0.9065 | 0.9463 | 0.9308 | 0.9483 | 0.9369 | 0.9412 | 0.9162 | 0.9441 | 0.9294 | 0.9373 |
| 1jbkA | 0.9095 | 0.9285 | 0.9066 | 0.9231 | 0.9198 | 0.9288 | 0.9476 | 0.9407 | 0.9485 | 0.9533 | 0.9412 | 0.9508 | 0.9511 | 0.952 | 0.9542 | 0.9505 | 0.9516 | 0.9444 |
| 1jfuA | 0.887 | 0.9229 | 0.9172 | 0.9334 | 0.9286 | 0.9341 | 0.9315 | 0.9456 | 0.9416 | 0.9513 | 0.9349 | 0.9515 | 0.9293 | 0.9541 | 0.9305 | 0.9516 | 0.9271 | 0.9481 |
| 1jfxA | 0.9265 | 0.9413 | 0.9325 | 0.9366 | 0.9289 | 0.9286 | 0.943 | 0.9571 | 0.9434 | 0.9618 | 0.9493 | 0.9635 | 0.9374 | 0.9597 | 0.9578 | 0.9619 | 0.951 | 0.9638 |
| 1jkxA | 0.9085 | 0.9224 | 0.9108 | 0.9503 | 0.9103 | 0.9487 | 0.9444 | 0.9522 | 0.9329 | 0.9521 | 0.9385 | 0.9592 | 0.916 | 0.9573 | 0.9512 | 0.9685 | 0.9258 | 0.9681 |
| 1jl1A | 0.8901 | 0.9116 | 0.9088 | 0.9176 | 0.9317 | 0.9145 | 0.9313 | 0.9288 | 0.9118 | 0.9468 | 0.9232 | 0.9353 | 0.9292 | 0.95 | 0.9532 | 0.9453 | 0.9347 | 0.9401 |
| 1jo0A | 0.8378 | 0.8921 | 0.8666 | 0.9186 | 0.8508 | 0.9345 | 0.8434 | 0.9449 | 0.8189 | 0.9404 | 0.8497 | 0.9385 | 0.8143 | 0.9463 | 0.8648 | 0.95 | 0.8492 | 0.9535 |
| 1jo8A | 0.6665 | 0.7258 | 0.7371 | 0.8019 | 0.782 | 0.8531 | 0.7952 | 0.8891 | 0.7806 | 0.8348 | 0.8 | 0.8629 | 0.8131 | 0.8556 | 0.806 | 0.8879 | 0.7643 | 0.8833 |
| 1josA | 0.8274 | 0.8168 | 0.8084 | 0.8394 | 0.8236 | 0.8283 | 0.8605 | 0.8855 | 0.868 | 0.8808 | 0.8551 | 0.8874 | 0.8756 | 0.8956 | 0.8676 | 0.8992 | 0.8745 | 0.92 |
| 1jvwA | 0.8999 | 0.8376 | 0.8979 | 0.8733 | 0.8814 | 0.8973 | 0.8837 | 0.9137 | 0.894 | 0.9132 | 0.9041 | 0.9388 | 0.9029 | 0.935 | 0.8761 | 0.9228 | 0.8943 | 0.9134 |
| 1jwqA | 0.9282 | 0.94 | 0.9371 | 0.9476 | 0.9432 | 0.9542 | 0.9398 | 0.9555 | 0.9445 | 0.963 | 0.9539 | 0.9665 | 0.9564 | 0.9721 | 0.9493 | 0.9657 | 0.9542 | 0.97 |
| 1jyhA | 0.8136 | 0.7026 | 0.8059 | 0.7666 | 0.8742 | 0.9033 | 0.913 | 0.9401 | 0.9111 | 0.937 | 0.9247 | 0.9538 | 0.9282 | 0.9441 | 0.908 | 0.9393 | 0.9216 | 0.9425 |
| 1k6kA | 0.9361 | 0.8768 | 0.95 | 0.9235 | 0.9491 | 0.9295 | 0.9535 | 0.9418 | 0.9513 | 0.95 | 0.9342 | 0.9558 | 0.9368 | 0.9406 | 0.9588 | 0.9572 | 0.9335 | 0.9423 |
| 1k7cA | 0.941 | 0.9534 | 0.9413 | 0.9511 | 0.9635 | 0.9524 | 0.9694 | 0.9577 | 0.9674 | 0.9653 | 0.9723 | 0.9721 | 0.9687 | 0.9704 | 0.9706 | 0.9711 | 0.9732 | 0.9669 |
| 1k7jA | 0.914 | 0.9487 | 0.9189 | 0.9482 | 0.9248 | 0.9523 | 0.9469 | 0.97 | 0.9383 | 0.9708 | 0.9574 | 0.9679 | 0.9561 | 0.9655 | 0.9565 | 0.9687 | 0.9569 | 0.9727 |
| 1kidA | 0.8937 | 0.8983 | 0.9236 | 0.9063 | 0.921 | 0.9195 | 0.9202 | 0.9194 | 0.9407 | 0.9354 | 0.9221 | 0.9352 | 0.9456 | 0.9422 | 0.9294 | 0.9394 | 0.928 | 0.9449 |
| 1kq6A | 0.8205 | 0.862 | 0.8143 | 0.8725 | 0.8295 | 0.8705 | 0.8365 | 0.9179 | 0.8686 | 0.9159 | 0.8877 | 0.9269 | 0.8642 | 0.9013 | 0.8668 | 0.932 | 0.8676 | 0.9133 |
| 1kqrA | 0.8877 | 0.9116 | 0.8625 | 0.9102 | 0.8878 | 0.9243 | 0.9089 | 0.9263 | 0.8878 | 0.9387 | 0.8872 | 0.9332 | 0.8909 | 0.9376 | 0.8717 | 0.9458 | 0.8849 | 0.9458 |
| 1ktgA | 0.8604 | 0.9039 | 0.8438 | 0.9152 | 0.8429 | 0.9181 | 0.8579 | 0.9385 | 0.8537 | 0.9267 | 0.8643 | 0.9338 | 0.8965 | 0.9475 | 0.9139 | 0.9481 | 0.8505 | 0.9533 |
| 1ku3A | 0.7056 | 0.7961 | 0.8186 | 0.8303 | 0.8424 | 0.8411 | 0.8572 | 0.8616 | 0.8685 | 0.8686 | 0.891 | 0.8466 | 0.8846 | 0.9003 | 0.8687 | 0.8758 | 0.8717 | 0.8984 |
| 1kw4A | 0.8518 | 0.7973 | 0.8828 | 0.8508 | 0.8835 | 0.8773 | 0.8206 | 0.8674 | 0.9042 | 0.9014 | 0.9083 | 0.889 | 0.9033 | 0.888 | 0.8853 | 0.8951 | 0.8816 | 0.8811 |
| 1lm4A | 0.8461 | 0.8553 | 0.8579 | 0.881 | 0.8137 | 0.8748 | 0.8466 | 0.8944 | 0.8305 | 0.8862 | 0.8685 | 0.8932 | 0.8488 | 0.8884 | 0.8651 | 0.8905 | 0.8481 | 0.8972 |
| 1lo7A | 0.8526 | 0.8408 | 0.8605 | 0.8767 | 0.8499 | 0.8629 | 0.8192 | 0.8971 | 0.8781 | 0.9167 | 0.9013 | 0.9055 | 0.861 | 0.9008 | 0.8863 | 0.9217 | 0.882 | 0.9216 |
| 1lpyA | 0.85 | 0.821 | 0.8305 | 0.8635 | 0.8886 | 0.8833 | 0.8388 | 0.8931 | 0.872 | 0.903 | 0.8771 | 0.9065 | 0.9032 | 0.9238 | 0.8766 | 0.9317 | 0.8615 | 0.9402 |
| 1m4jA | 0.8383 | 0.8907 | 0.848 | 0.9261 | 0.8629 | 0.9315 | 0.903 | 0.9157 | 0.8826 | 0.9275 | 0.8886 | 0.9414 | 0.8804 | 0.9342 | 0.8463 | 0.9359 | 0.8652 | 0.9343 |
| 1m8aA | 0.6981 | 0.8043 | 0.7904 | 0.8222 | 0.7401 | 0.7879 | 0.7926 | 0.7973 | 0.7628 | 0.8395 | 0.7357 | 0.8851 | 0.8534 | 0.8651 | 0.7537 | 0.8694 | 0.7919 | 0.8902 |
| 1mk0A | 0.8502 | 0.9119 | 0.824 | 0.9153 | 0.8427 | 0.9079 | 0.8766 | 0.9301 | 0.9194 | 0.9451 | 0.9164 | 0.9524 | 0.8997 | 0.9556 | 0.8966 | 0.9334 | 0.8779 | 0.9304 |
| 1mugA | 0.8913 | 0.9252 | 0.9361 | 0.923 | 0.9346 | 0.9331 | 0.9395 | 0.941 | 0.9364 | 0.948 | 0.9416 | 0.9496 | 0.9341 | 0.9479 | 0.9415 | 0.9509 | 0.9409 | 0.9488 |
| 1nb9A | 0.8602 | 0.8605 | 0.8458 | 0.8784 | 0.8966 | 0.8994 | 0.8842 | 0.9257 | 0.8831 | 0.9336 | 0.8734 | 0.9343 | 0.8613 | 0.9439 | 0.9066 | 0.9391 | 0.8877 | 0.9361 |
| 1ne2A | 0.8725 | 0.8852 | 0.8468 | 0.8834 | 0.8729 | 0.899 | 0.8631 | 0.8918 | 0.8718 | 0.9019 | 0.8697 | 0.9117 | 0.9044 | 0.906 | 0.8867 | 0.9065 | 0.8858 | 0.9073 |
| 1npsA | 0.8641 | 0.8658 | 0.8692 | 0.8697 | 0.855 | 0.8892 | 0.8774 | 0.8938 | 0.8812 | 0.9214 | 0.8994 | 0.9211 | 0.8975 | 0.9306 | 0.9003 | 0.9108 | 0.9017 | 0.9122 |
| 1nrvA | 0.8643 | 0.8386 | 0.8872 | 0.8613 | 0.932 | 0.8966 | 0.9112 | 0.9149 | 0.9321 | 0.9103 | 0.9224 | 0.9361 | 0.9304 | 0.939 | 0.9279 | 0.9504 | 0.9113 | 0.9403 |

| | | | | | | | | | | | | | | | | | | |
|---|---|---|---|---|---|---|---|---|---|---|---|---|---|---|---|---|---|---|
| 1ny1A | 0.9516 | 0.9553 | 0.9472 | 0.9539 | 0.959 | 0.9556 | 0.9575 | 0.9636 | 0.9601 | 0.9667 | 0.9682 | 0.9708 | 0.9613 | 0.9731 | 0.9723 | 0.977 | 0.9641 | 0.9733 |
| 1o1zA | 0.9089 | 0.9365 | 0.9337 | 0.9475 | 0.9433 | 0.9558 | 0.9465 | 0.9663 | 0.9576 | 0.9661 | 0.9549 | 0.9672 | 0.9602 | 0.9611 | 0.9399 | 0.9628 | 0.9591 | 0.9653 |
| 1p90A | 0.8641 | 0.8815 | 0.9023 | 0.8738 | 0.913 | 0.9023 | 0.9036 | 0.9151 | 0.9145 | 0.9318 | 0.9133 | 0.9393 | 0.9258 | 0.9367 | 0.926 | 0.939 | 0.9227 | 0.9394 |
| 1pchA | 0.9106 | 0.9277 | 0.9416 | 0.925 | 0.9392 | 0.9265 | 0.9421 | 0.9227 | 0.9496 | 0.933 | 0.9601 | 0.9354 | 0.9507 | 0.9319 | 0.9542 | 0.9409 | 0.9563 | 0.9303 |
| 1pkoA | 0.7952 | 0.8411 | 0.8163 | 0.8587 | 0.7558 | 0.8611 | 0.7215 | 0.8922 | 0.7696 | 0.8954 | 0.8136 | 0.8773 | 0.7972 | 0.903 | 0.7691 | 0.9045 | 0.8021 | 0.8923 |
| 1qf9A | 0.9305 | 0.9285 | 0.9197 | 0.947 | 0.9548 | 0.9521 | 0.9359 | 0.9554 | 0.9444 | 0.9557 | 0.9563 | 0.9565 | 0.9569 | 0.9657 | 0.9572 | 0.9647 | 0.9509 | 0.9644 |
| 1qjpA | 0.7107 | 0.791 | 0.7633 | 0.7719 | 0.7944 | 0.7642 | 0.7764 | 0.7937 | 0.8157 | 0.8203 | 0.8 | 0.836 | 0.8205 | 0.8707 | 0.8254 | 0.8905 | 0.8688 | 0.8941 |
| 1ql0A | 0.9324 | 0.9305 | 0.9334 | 0.9463 | 0.9343 | 0.9558 | 0.9446 | 0.966 | 0.9358 | 0.9686 | - | 0.9643 | 0.9579 | 0.9686 | 0.9619 | 0.969 | 0.9605 | 0.9688 |
| 1r26A | 0.882 | 0.885 | 0.8642 | 0.8994 | 0.8631 | 0.9247 | 0.8682 | 0.9281 | 0.8703 | 0.9486 | 0.8851 | 0.9405 | 0.8883 | 0.9463 | 0.8831 | 0.9481 | 0.8575 | 0.9553 |
| 1roaA | 0.7676 | 0.8515 | 0.8476 | 0.9003 | 0.8073 | 0.9224 | 0.8379 | 0.8857 | 0.8003 | 0.8798 | 0.8532 | 0.9073 | 0.8301 | 0.9036 | 0.8396 | 0.9032 | 0.8429 | 0.9196 |
| 1rw1A | 0.8708 | 0.9005 | 0.837 | 0.9033 | 0.9267 | 0.9176 | 0.9071 | 0.924 | 0.9323 | 0.9335 | 0.9506 | 0.9333 | 0.9252 | 0.9397 | 0.9372 | 0.9357 | 0.8994 | 0.9431 |
| 1rw7A | 0.9429 | 0.946 | 0.9541 | 0.9511 | 0.9519 | 0.9533 | 0.9555 | 0.9597 | 0.9441 | 0.9604 | 0.9608 | 0.95 | 0.958 | 0.9588 | 0.9697 | 0.9584 | 0.9676 | 0.9451 |
| 1rybA | 0.9284 | 0.9057 | 0.9148 | 0.9172 | 0.9308 | 0.9388 | 0.9441 | 0.9388 | 0.9365 | 0.937 | 0.9526 | 0.9384 | 0.947 | 0.939 | 0.9481 | 0.9372 | 0.9422 | 0.9418 |
| 1smxA | 0.6853 | 0.7306 | - | 0.7942 | 0.7947 | 0.8347 | 0.7874 | 0.8256 | 0.7615 | 0.8421 | 0.8063 | 0.8527 | 0.8387 | 0.8638 | 0.7732 | 0.8621 | 0.8076 | 0.8652 |
| 1svyA | 0.8698 | 0.8505 | 0.8698 | 0.894 | 0.9183 | 0.9225 | 0.9251 | 0.9093 | 0.9066 | 0.9237 | 0.9298 | 0.9439 | 0.9183 | 0.9295 | 0.9267 | 0.9367 | 0.8986 | 0.9196 |
| 1t8kA | 0.8409 | 0.9077 | 0.9372 | 0.9129 | 0.8889 | 0.9012 | 0.9029 | 0.8869 | 0.9107 | 0.9496 | 0.94 | 0.9455 | 0.9302 | 0.9447 | 0.9408 | 0.9514 | 0.9352 | 0.9395 |
| 1tifA | 0.7453 | 0.6819 | 0.7113 | 0.7081 | 0.6813 | 0.6963 | 0.8038 | 0.7596 | 0.8287 | 0.7474 | 0.8321 | 0.8248 | 0.869 | 0.857 | 0.8311 | 0.8125 | 0.8209 | 0.8213 |
| 1tqgA | 0.9352 | 0.9058 | 0.9287 | 0.9263 | 0.9518 | 0.9139 | 0.9427 | 0.946 | 0.9633 | 0.945 | 0.9574 | 0.959 | 0.9625 | 0.9462 | 0.9488 | 0.958 | 0.955 | 0.9495 |
| 1tqhA | 0.9226 | 0.9305 | 0.9473 | 0.9433 | 0.9636 | 0.9443 | 0.9557 | 0.967 | 0.9569 | 0.9604 | 0.9669 | 0.9716 | 0.9771 | 0.9738 | 0.963 | 0.9769 | 0.9707 | 0.9732 |
| 1tzvA | 0.9256 | 0.9244 | 0.9399 | 0.9456 | 0.9448 | 0.9486 | 0.955 | 0.9482 | 0.9427 | 0.9693 | 0.9418 | 0.9602 | 0.9285 | 0.9587 | 0.9652 | 0.9652 | 0.9628 | 0.9737 |
| 1vfyA | 0.5636 | 0.6441 | 0.6332 | 0.7346 | 0.6395 | 0.8188 | 0.6679 | 0.8192 | 0.6579 | 0.8343 | 0.6263 | 0.8166 | 0.6901 | 0.8679 | 0.6511 | 0.8833 | 0.6893 | 0.838 |
| 1vhuA | 0.9431 | 0.9534 | 0.9389 | 0.958 | 0.9452 | 0.954 | 0.9592 | 0.961 | 0.9621 | 0.9654 | 0.9611 | 0.9646 | 0.9571 | 0.9553 | 0.9615 | 0.9612 | 0.9517 | 0.9567 |
| 1vjkA | 0.8664 | 0.8599 | 0.8862 | 0.8658 | 0.9075 | 0.89 | 0.887 | 0.8976 | 0.9248 | 0.9071 | 0.9191 | 0.922 | 0.9252 | 0.9276 | 0.9298 | 0.9266 | 0.9116 | 0.938 |
| 1vmbA | 0.8019 | 0.8249 | 0.8065 | 0.8217 | 0.7964 | 0.8166 | 0.853 | 0.8316 | 0.8393 | 0.8831 | 0.8479 | 0.9141 | 0.8448 | 0.8988 | 0.8028 | 0.9061 | 0.852 | 0.9216 |
| 1vp6A | 0.8662 | 0.908 | 0.8691 | 0.9197 | 0.8667 | 0.9302 | 0.8885 | 0.9428 | 0.9086 | 0.9456 | 0.8981 | 0.9521 | 0.8864 | 0.9485 | 0.9149 | 0.9471 | 0.9206 | 0.9514 |
| 1w0hA | 0.9052 | 0.9353 | 0.8826 | 0.9441 | 0.9158 | 0.9559 | 0.9236 | 0.9506 | 0.9296 | 0.9595 | 0.9325 | 0.9666 | 0.9364 | 0.965 | 0.9318 | 0.968 | 0.9381 | 0.9648 |
| 1whiA | 0.8322 | 0.8763 | 0.827 | 0.8956 | 0.8409 | 0.8935 | 0.7931 | 0.9108 | 0.8124 | 0.9238 | 0.8189 | 0.9106 | 0.8487 | 0.9 | 0.8862 | 0.9187 | 0.8569 | 0.8801 |
| 1wjxA | 0.8162 | 0.8537 | 0.8589 | 0.8893 | 0.8364 | 0.8805 | 0.8718 | 0.9128 | 0.8383 | 0.9151 | 0.8748 | 0.9283 | 0.8751 | 0.9242 | 0.8662 | 0.9288 | 0.8645 | 0.9331 |
| 1wkcA | 0.9198 | 0.9089 | 0.9003 | 0.9258 | 0.9367 | 0.9235 | 0.9167 | 0.9442 | 0.9232 | 0.9414 | 0.9376 | 0.9469 | 0.94 | 0.9511 | 0.9468 | 0.9509 | 0.9392 | 0.9383 |
| 1xdzA | 0.9336 | 0.9433 | 0.9099 | 0.9446 | 0.9364 | 0.9514 | 0.9422 | 0.9541 | 0.9446 | 0.9623 | 0.9448 | 0.9627 | 0.9533 | 0.966 | 0.9529 | 0.9607 | 0.9493 | 0.9579 |
| 1xffA | 0.9449 | 0.9481 | 0.9542 | 0.9579 | 0.9578 | 0.9609 | 0.9613 | 0.9687 | 0.9693 | 0.9688 | 0.9763 | 0.9687 | 0.969 | 0.9674 | 0.973 | 0.9694 | 0.9712 | 0.9664 |
| 1xkrA | 0.9338 | 0.9195 | 0.9231 | 0.9353 | 0.9551 | 0.9534 | 0.9531 | 0.9504 | 0.958 | 0.9593 | 0.9613 | 0.9662 | 0.9715 | 0.9612 | 0.9662 | 0.9652 | 0.9665 | 0.9642 |
| 2arcA | 0.9092 | 0.9183 | 0.8805 | 0.9257 | 0.9139 | 0.9445 | 0.917 | 0.9539 | 0.9183 | 0.9568 | 0.9138 | 0.9553 | 0.9241 | 0.9603 | 0.9325 | 0.9576 | 0.9332 | 0.9563 |
| 2cuaA | 0.9265 | 0.8892 | 0.8836 | 0.9097 | 0.9153 | 0.925 | 0.888 | 0.932 | 0.9248 | 0.9312 | 0.8776 | 0.9368 | 0.9291 | 0.9369 | 0.9132 | 0.9323 | 0.9114 | 0.9224 |
| 2hs1A | 0.8225 | 0.8099 | 0.811 | 0.7852 | 0.7954 | 0.8634 | 0.8296 | 0.8891 | 0.8445 | 0.8993 | 0.8648 | 0.8665 | 0.8581 | 0.8659 | 0.8663 | 0.91 | 0.8798 | 0.9028 |
| 2mhrA | 0.919 | 0.9052 | 0.9442 | 0.9194 | 0.9419 | 0.9321 | 0.9418 | 0.9394 | 0.9598 | 0.9439 | 0.9477 | 0.9448 | 0.9661 | 0.9515 | 0.9278 | 0.9525 | 0.9548 | 0.965 |
| 2phyA | 0.877 | 0.9052 | 0.8883 | 0.9094 | 0.8753 | 0.9165 | 0.8964 | 0.9267 | 0.9123 | 0.9457 | 0.9107 | 0.9521 | 0.9544 | 0.9488 | 0.9194 | 0.9538 | 0.9244 | 0.9374 |

| | | | | | | | | | | | | | | | | | | |
|---|---|---|---|---|---|---|---|---|---|---|---|---|---|---|---|---|---|---|
| 2tpsA | 0.9348 | 0.9421 | 0.9619 | 0.955 | 0.9698 | 0.9512 | 0.9648 | 0.9632 | 0.9697 | 0.9598 | 0.9707 | 0.9573 | 0.968 | 0.9634 | 0.9744 | 0.9652 | 0.9749 | 0.9625 |
| 2vxnA | 0.9416 | 0.954 | 0.9528 | 0.951 | 0.9183 | 0.9583 | 0.9641 | 0.9617 | 0.9709 | 0.9617 | 0.9573 | 0.9699 | 0.9789 | 0.9729 | 0.9698 | 0.9756 | 0.9667 | 0.9682 |
| 3borA | 0.9184 | 0.9339 | 0.9354 | 0.941 | 0.9415 | 0.9429 | 0.9475 | 0.9471 | 0.9373 | 0.9534 | 0.9421 | 0.9436 | 0.9516 | 0.9485 | 0.9483 | 0.9475 | 0.9549 | 0.9468 |
| 3dqgA | 0.7395 | 0.8137 | 0.7476 | 0.841 | 0.7516 | 0.8591 | 0.7986 | 0.8992 | 0.7481 | 0.9071 | 0.7996 | 0.8979 | 0.7807 | 0.8967 | 0.8526 | 0.9285 | 0.8603 | 0.9178 |
| 5ptpA | 0.9219 | 0.9448 | 0.9056 | 0.9517 | 0.9259 | 0.9564 | 0.9249 | 0.959 | 0.9325 | 0.9571 | 0.9193 | 0.9606 | 0.9259 | 0.9634 | 0.9336 | 0.9603 | 0.9352 | 0.9546 |
| Mean | 0.86 | 0.88 | 0.87 | 0.89 | 0.88 | 0.90 | 0.89 | 0.91 | 0.90 | 0.92 | 0.90 | 0.93 | 0.91 | 0.93 | 0.90 | 0.93 | 0.90 | 0.93 |
| Median | 0.87 | 0.90 | 0.88 | 0.91 | 0.90 | 0.92 | 0.90 | 0.93 | 0.92 | 0.93 | 0.92 | 0.94 | 0.92 | 0.94 | 0.92 | 0.94 | 0.92 | 0.94 |

**S3 Table.** Target-by-target reconstruction performance on 150 soluble proteins for true $C_\alpha$–$C_\alpha$ hybrid interaction maps at tri-level thresholding.

| | |
|---|---|
| 1a3aA | 0.9821 |
| 1a6mA | 0.98 |
| 1a70A | 0.9733 |
| 1aapA | 0.8596 |
| 1abaA | 0.9519 |
| 1ag6A | 0.9733 |
| 1aoeA | 0.9848 |
| 1atlA | 0.9882 |
| 1atzA | 0.934 |
| 1avsA | 0.9555 |
| 1bdoA | 0.9554 |
| 1bebA | 0.9849 |
| 1behA | 0.987 |
| 1bkrA | 0.9767 |
| 1brfA | 0.8998 |
| 1bsgA | 0.9916 |
| 1c44A | 0.932 |
| 1c52A | 0.9761 |
| 1c9oA | 0.9363 |
| 1cc8A | 0.9561 |
| 1chdA | 0.9858 |
| 1cjwA | 0.9655 |
| 1ckeA | 0.9241 |
| 1ctfA | 0.9374 |
| 1cxyA | 0.9528 |
| 1cznA | 0.9785 |
| 1d0qA | 0.9704 |
| 1d1qA | 0.977 |
| 1d4oA | 0.9815 |
| 1dbxA | 0.9682 |
| 1dixA | 0.9831 |
| 1dlwA | 0.9658 |
| 1dmgA | 0.9441 |
| 1dqgA | 0.9805 |
| 1dsxA | 0.9696 |
| 1eazA | 0.9699 |
| 1ej0A | 0.9829 |
| 1ej8A | 0.9762 |
| 1ek0A | 0.9755 |

| | |
|---|---|
| 1f6bA | 0.9569 |
| 1fcyA | 0.9879 |
| 1fk5A | 0.9528 |
| 1fl0A | 0.9823 |
| 1fnaA | 0.9624 |
| 1fqtA | 0.9744 |
| 1fvgA | 0.9818 |
| 1fvkA | 0.9803 |
| 1fx2A | 0.9411 |
| 1g2rA | 0.9546 |
| 1g9oA | 0.9185 |
| 1gbsA | 0.9793 |
| 1gmiA | 0.973 |
| 1gmxA | 0.9697 |
| 1guuA | 0.9274 |
| 1gz2A | 0.9631 |
| 1gzcA | 0.9884 |
| 1h0pA | 0.9875 |
| 1h2eA | 0.9882 |
| 1h4xA | 0.9795 |
| 1h98A | 0.9474 |
| 1hdoA | 0.9875 |
| 1hfcA | 0.9806 |
| 1hh8A | 0.9794 |
| 1htwA | 0.9808 |
| 1hxnA | 0.9844 |
| 1i1jA | 0.9522 |
| 1i1nA | 0.9883 |
| 1i4jA | 0.9251 |
| 1i58A | 0.9645 |
| 1i5gA | 0.9816 |
| 1i71A | 0.9528 |
| 1ihzA | 0.9747 |
| 1iibA | 0.9618 |
| 1im5A | 0.9853 |
| 1iwdA | 0.9873 |
| 1j3aA | 0.9708 |
| 1jbeA | 0.9747 |
| 1jbkA | 0.9754 |
| 1jfuA | 0.9812 |
| 1jfxA | 0.9916 |

| | |
|---|---|
| 1jkxA | 0.9845 |
| 1jl1A | 0.9742 |
| 1jo0A | 0.9767 |
| 1jo8A | 0.9451 |
| 1josA | 0.945 |
| 1jvwA | 0.9732 |
| 1jwqA | 0.9868 |
| 1jyhA | 0.9773 |
| 1k6kA | 0.9798 |
| 1k7cA | 0.9889 |
| 1k7jA | 0.9906 |
| 1kidA | 0.9763 |
| 1kq6A | 0.9496 |
| 1kqrA | 0.9775 |
| 1ktgA | 0.9648 |
| 1ku3A | 0.9281 |
| 1kw4A | 0.9486 |
| 1lm4A | 0.8353 |
| 1lo7A | 0.9721 |
| 1lpyA | 0.974 |
| 1m4jA | 0.9675 |
| 1m8aA | 0.9473 |
| 1mk0A | 0.9759 |
| 1mugA | 0.9816 |
| 1nb9A | 0.9681 |
| 1ne2A | 0.9416 |
| 1npsA | 0.9535 |
| 1nrvA | 0.9684 |
| 1ny1A | 0.9925 |
| 1o1zA | 0.9859 |
| 1p90A | 0.9783 |
| 1pchA | 0.9813 |
| 1pkoA | 0.9301 |
| 1qf9A | 0.9859 |
| 1qjpA | 0.915 |
| 1ql0A | 0.9891 |
| 1r26A | 0.9762 |
| 1roaA | 0.966 |
| 1rw1A | 0.9755 |
| 1rw7A | 0.988 |
| 1rybA | 0.9751 |

| | |
|---|---|
| 1smxA | 0.9336 |
| 1svyA | 0.967 |
| 1t8kA | 0.9648 |
| 1tifA | 0.8781 |
| 1tqgA | 0.9726 |
| 1tqhA | 0.989 |
| 1tzvA | 0.9745 |
| 1vfyA | 0.8971 |
| 1vhuA | 0.9847 |
| 1vjkA | 0.9689 |
| 1vmbA | 0.9424 |
| 1vp6A | 0.9779 |
| 1w0hA | 0.9897 |
| 1whiA | 0.9605 |
| 1wjxA | 0.9606 |
| 1wkcA | 0.9795 |
| 1xdzA | 0.9885 |
| 1xffA | 0.9915 |
| 1xkrA | 0.985 |
| 2arcA | 0.9811 |
| 2cuaA | 0.9771 |
| 2hs1A | 0.9623 |
| 2mhrA | 0.9808 |
| 2phyA | 0.9819 |
| 2tpsA | 0.992 |
| 2vxnA | 0.9874 |
| 3borA | 0.9683 |
| 3dqgA | 0.9337 |
| 5ptpA | 0.9857 |
| Mean | 0.966608667 |
| Median | 0.9749 |

**S4 Table.** Target-by-target reconstruction performance on 150 soluble proteins for true $C_\beta$–$C_\beta$ hybrid interaction maps at tri-level thresholding.

| | |
|---|---|
| 1a3aA | 0.9522 |
| 1a6mA | 0.9386 |
| 1a70A | 0.9318 |
| 1aapA | 0.7809 |
| 1abaA | 0.8931 |
| 1ag6A | 0.937 |
| 1aoeA | 0.9635 |
| 1atlA | 0.9674 |
| 1atzA | 0.9393 |
| 1avsA | 0.8859 |
| 1bdoA | 0.8996 |
| 1bebA | 0.9735 |
| 1behA | 0.9489 |
| 1bkrA | 0.9543 |
| 1brfA | 0.8717 |
| 1bsgA | 0.9617 |
| 1c44A | 0.8659 |
| 1c52A | 0.9312 |
| 1c9oA | 0.8678 |
| 1cc8A | 0.9017 |
| 1chdA | 0.9544 |
| 1cjwA | 0.9447 |
| 1ckeA | 0.8949 |
| 1ctfA | 0.8866 |
| 1cxyA | 0.9361 |
| 1cznA | 0.9458 |
| 1d0qA | 0.8763 |
| 1d1qA | 0.9499 |
| 1d4oA | 0.9531 |
| 1dbxA | 0.9264 |
| 1dixA | 0.9509 |
| 1dlwA | 0.9382 |
| 1dmgA | 0.9115 |
| 1dqgA | 0.9513 |
| 1dsxA | 0.9266 |
| 1eazA | 0.9541 |
| 1ej0A | 0.9494 |
| 1ej8A | 0.9397 |
| 1ek0A | 0.9483 |

| | |
|---|---|
| 1f6bA | 0.9183 |
| 1fcyA | 0.9687 |
| 1fk5A | 0.8909 |
| 1fl0A | 0.9531 |
| 1fnaA | 0.9138 |
| 1fqtA | 0.9295 |
| 1fvgA | 0.9499 |
| 1fvkA | 0.9345 |
| 1fx2A | 0.9194 |
| 1g2rA | 0.9194 |
| 1g9oA | 0.8824 |
| 1gbsA | 0.9442 |
| 1gmiA | 0.9504 |
| 1gmxA | 0.9266 |
| 1guuA | 0.9077 |
| 1gz2A | 0.9393 |
| 1gzcA | 0.9694 |
| 1h0pA | 0.9499 |
| 1h2eA | 0.9634 |
| 1h4xA | 0.9258 |
| 1h98A | 0.9271 |
| 1hdoA | 0.9598 |
| 1hfcA | 0.9477 |
| 1hh8A | 0.9635 |
| 1htwA | 0.9514 |
| 1hxnA | 0.9586 |
| 1i1jA | 0.331 |
| 1i1nA | 0.9551 |
| 1i4jA | 0.9102 |
| 1i58A | 0.937 |
| 1i5gA | 0.9596 |
| 1i71A | 0.8989 |
| 1ihzA | 0.9439 |
| 1iibA | 0.9056 |
| 1im5A | 0.9559 |
| 1iwdA | 0.9528 |
| 1j3aA | 0.9524 |
| 1jbeA | 0.9436 |
| 1jbkA | 0.9489 |
| 1jfuA | 0.9499 |
| 1jfxA | 0.9615 |

| | |
|---|---|
| 1jkxA | 0.9652 |
| 1jl1A | 0.9436 |
| 1jo0A | 0.952 |
| 1jo8A | 0.9096 |
| 1josA | 0.9302 |
| 1jvwA | 0.9146 |
| 1jwqA | 0.9694 |
| 1jyhA | 0.9404 |
| 1k6kA | 0.9531 |
| 1k7cA | 0.9701 |
| 1k7jA | 0.9668 |
| 1kidA | 0.9309 |
| 1kq6A | 0.9386 |
| 1kqrA | 0.9569 |
| 1ktgA | 0.9614 |
| 1ku3A | 0.9001 |
| 1kw4A | 0.9273 |
| 1lm4A | 0.8913 |
| 1lo7A | 0.9497 |
| 1lpyA | 0.9354 |
| 1m4jA | 0.9331 |
| 1m8aA | 0.9283 |
| 1mk0A | 0.9472 |
| 1mugA | 0.951 |
| 1nb9A | 0.9432 |
| 1ne2A | 0.8998 |
| 1npsA | 0.8947 |
| 1nrvA | 0.9555 |
| 1ny1A | 0.9709 |
| 1o1zA | 0.9632 |
| 1p90A | 0.9305 |
| 1pchA | 0.9476 |
| 1pkoA | 0.9038 |
| 1qf9A | 0.9585 |
| 1qjpA | 0.9104 |
| 1ql0A | 0.9676 |
| 1r26A | 0.9599 |
| 1roaA | 0.9401 |
| 1rw1A | 0.9462 |
| 1rw7A | 0.9524 |
| 1rybA | 0.9418 |

| | |
|---|---|
| 1smxA | 0.8576 |
| 1svyA | 0.9227 |
| 1t8kA | 0.947 |
| 1tifA | 0.8956 |
| 1tqgA | 0.9545 |
| 1tqhA | 0.9711 |
| 1tzvA | 0.9768 |
| 1vfyA | 0.8481 |
| 1vhuA | 0.9562 |
| 1vjkA | 0.9466 |
| 1vmbA | 0.9098 |
| 1vp6A | 0.943 |
| 1w0hA | 0.9678 |
| 1whiA | 0.8731 |
| 1wjxA | 0.9364 |
| 1wkcA | 0.9406 |
| 1xdzA | 0.9558 |
| 1xffA | 0.9604 |
| 1xkrA | 0.959 |
| 2arcA | 0.9489 |
| 2cuaA | 0.9326 |
| 2hs1A | 0.9147 |
| 2mhrA | 0.9612 |
| 2phyA | 0.9378 |
| 2tpsA | 0.9471 |
| 2vxnA | 0.9647 |
| 3borA | 0.9515 |
| 3dqgA | 0.907 |
| 5ptpA | 0.9498 |
| Mean | 0.931099333 |
| Median | 0.94375 |

**S5 Table.** Target-by-target *ab initio* folding performance on 40 CASP FM targets.

| Targets | DConStruct | DMPfold | CONFOLD2 | ROSETTA | CGLFold |
|---|---|---|---|---|---|
| T0859-D1 | 0.193 | 0.2446 | 0.1617 | 0.2232 | 0.19 |
| T0862-D1 | 0.5056 | 0.2755 | 0.2599 | 0.3039 | 0.61 |
| T0863-D1 | 0.503 | 0.2933 | 0.3374 | 0.2979 | 0.53 |
| T0863-D2 | 0.2296 | 0.1721 | 0.2728 | 0.2702 | 0.39 |
| T0864-D1 | 0.695 | 0.4792 | 0.5638 | 0.2935 | 0.28 |
| T0866-D1 | 0.582 | 0.7369 | 0.6356 | 0.6355 | 0.55 |
| T0869-D1 | 0.7448 | 0.7729 | 0.6522 | 0.7132 | 0.47 |
| T0870-D1 | 0.6724 | 0.4912 | 0.5863 | 0.6267 | 0.56 |
| T0886-D1 | 0.3042 | 0.3214 | 0.3095 | 0.3277 | 0.29 |
| T0886-D2 | 0.6944 | 0.6894 | 0.307 | 0.4633 | 0.5 |
| T0892-D2 | 0.696 | 0.6411 | 0.5397 | 0.4258 | 0.35 |
| T0896-D3 | 0.1558 | 0.1627 | 0.1773 | 0.1566 | 0.22 |
| T0897-D1 | 0.2031 | 0.2187 | 0.2038 | 0.2262 | 0.2 |
| T0897-D2 | 0.2122 | 0.2463 | 0.227 | 0.2518 | 0.29 |
| T0898-D1 | 0.6463 | 0.3716 | 0.4882 | 0.3413 | 0.52 |
| T0900-D1 | 0.6251 | 0.6207 | 0.2337 | 0.3583 | 0.45 |
| T0904-D1 | 0.4335 | 0.4073 | 0.7023 | 0.6011 | |
| T0912-D3 | 0.5784 | 0.5599 | 0.3651 | 0.2007 | 0.26 |
| T0918-D1 | 0.5547 | 0.5858 | 0.2849 | 0.3806 | 0.43 |
| T0918-D2 | 0.3157 | 0.5632 | 0.41 | 0.3767 | 0.47 |
| T0918-D3 | 0.5148 | 0.4087 | 0.3084 | 0.4842 | 0.43 |
| T0941-D1 | 0.2739 | 0.288 | 0.1869 | 0.2293 | 0.26 |
| T0950-D1 | 0.5019 | 0.3062 | 0.3278 | 0.3339 | 0.26 |
| T0953s1-D1 | 0.3997 | 0.3567 | 0.2486 | 0.2989 | 0.4 |
| T0953s2-D2 | 0.6466 | 0.5258 | 0.464 | 0.4689 | 0.3 |
| T0953s2-D3 | 0.5092 | 0.2404 | 0.2825 | 0.2266 | |
| T0957s1-D1 | 0.3496 | 0.214 | 0.3741 | 0.3645 | |
| T0957s2-D1 | 0.7022 | 0.5481 | 0.6063 | 0.5632 | 0.58 |
| T0963-D2 | 0.2306 | 0.2244 | 0.2863 | 0.2732 | 0.45 |
| T0968s1-D1 | 0.6823 | 0.6635 | 0.5202 | 0.5209 | 0.55 |
| T0968s2-D1 | 0.7371 | 0.5256 | 0.6818 | 0.3609 | 0.54 |
| T0960-D2 | 0.3601 | 0.25 | 0.2657 | 0.2546 | 0.41 |
| T0969-D1 | 0.7688 | 0.5698 | 0.4974 | 0.4909 | |
| T0980s1-D1 | 0.2905 | 0.2611 | 0.4544 | 0.4596 | |
| T0990-D1 | 0.3254 | 0.5156 | 0.2871 | 0.3605 | |
| T0990-D2 | 0.256 | 0.3714 | 0.2563 | 0.3679 | |
| T0990-D3 | 0.2479 | 0.2625 | 0.2414 | 0.2415 | |

| | | | | | |
|---|---|---|---|---|---|
| T1021s3-D1 | 0.4904 | 0.3951 | 0.4469 | 0.4878 | |
| T1021s3-D2 | 0.3345 | 0.2975 | 0.2939 | 0.2854 | |
| T1022s1-D1 | 0.2682 | 0.5253 | 0.5316 | 0.4479 | |
| Mean | 0.46 | 0.42 | 0.38 | 0.37 | 0.40 |
| Median | 0.50 | 0.38 | 0.32 | 0.36 | 0.43 |
| Correct fold | 20 | 15 | 10 | 6 | 8 |
| P-value | | 0.03329857 | 0.001616589 | 0.001311146 | |

**S6 Table.** Target-by-target *ab initio* folding performance on 510 membrane proteins.

| Target | DConStruct | Xu's DTL with CNS |
|---|---|---|
| 1a0sP | 0.5269 | 0.2815 |
| 1ar1B | 0.4934 | 0.651 |
| 1bccE | 0.5235 | 0.2483 |
| 1bctA | 0.4103 | 0.38 |
| 1bhaA | 0.6311 | 0.6281 |
| 1c17M | 0.4385 | 0.4613 |
| 1e7pC | 0.6162 | 0.6473 |
| 1ehkB | 0.2438 | 0.5301 |
| 1fftB | 0.561 | 0.2299 |
| 1fftC | 0.7429 | 0.5934 |
| 1fw2A | 0.6645 | 0.2897 |
| 1fx8A | 0.654 | 0.6239 |
| 1gzmA | 0.4837 | 0.7332 |
| 1h2sB | 0.5391 | 0.4751 |
| 1h6s1 | 0.6223 | 0.6571 |
| 1izlA | 0.3153 | 0.2777 |
| 1izlC | 0.515 | 0.4025 |
| 1jb0K | 0.4186 | 0.4397 |
| 1k24A | 0.6996 | 0.5633 |
| 1kf6C | 0.4795 | 0.4833 |
| 1kf6D | 0.556 | 0.4938 |
| 1kqfB | 0.6082 | 0.2832 |
| 1kqfC | 0.7434 | 0.6434 |
| 1kzuA | 0.4865 | 0.4428 |
| 1lghA | 0.5248 | 0.4573 |
| 1m56B | 0.5316 | 0.399 |
| 1m56D | 0.7187 | 0.6127 |
| 1m57A | 0.8704 | 0.8187 |
| 1mm4A | 0.2595 | 0.517 |
| 1mprA | 0.2657 | 0.284 |
| 1n7lA | 0.4774 | 0.5187 |
| 1nekC | 0.6404 | 0.6401 |
| 1nekD | 0.5867 | 0.706 |
| 1o5wA | 0.7687 | 0.3439 |
| 1occD | 0.3207 | 0.2525 |
| 1oedC | 0.4874 | 0.3664 |
| 1orsC | 0.5548 | 0.4434 |
| 1p49A | 0.6714 | 0.573 |
| 1p4tA | 0.7519 | 0.8164 |

| | | |
|---|---|---|
| 1p7bA | 0.3219 | 0.2976 |
| 1pw4A | 0.7725 | 0.7793 |
| 1q16C | 0.5805 | 0.6371 |
| 1q90A | 0.3747 | 0.238 |
| 1q90B | 0.6578 | 0.6961 |
| 1qcrD | 0.3155 | 0.3692 |
| 1qd6C | 0.69 | 0.6994 |
| 1qleC | 0.7464 | 0.6321 |
| 1rh5B | 0.3296 | 0.4668 |
| 1rh5C | 0.5023 | 0.6246 |
| 1rwtA | 0.4505 | 0.4808 |
| 1s5lB | 0.2487 | 0.2838 |
| 1s5lE | 0.3049 | 0.2851 |
| 1s5lX | 0.5962 | 0.668 |
| 1sqqK | 0.4734 | 0.5394 |
| 1t16A | 0.7173 | 0.6906 |
| 1tlwA | 0.5881 | 0.5191 |
| 1tqqA | 0.4957 | 0.492 |
| 1uunA | 0.6082 | 0.5962 |
| 1uynX | 0.3806 | 0.7472 |
| 1vclA | 0.2834 | 0.2259 |
| 1vf5B | 0.3353 | 0.304 |
| 1vf5D | 0.3171 | 0.2505 |
| 1wrgA | 0.4174 | 0.3969 |
| 1xioA | 0.8301 | 0.8187 |
| 1xl4A | 0.4535 | 0.2865 |
| 1yc9A | 0.4663 | 0.3928 |
| 1yewC | 0.5809 | 0.3008 |
| 1yq3C | 0.5559 | 0.5304 |
| 1yq3D | 0.6778 | 0.5006 |
| 1zrtE | 0.2585 | 0.4679 |
| 1zzaA | 0.2848 | 0.3265 |
| 2a0lA | 0.3574 | 0.3828 |
| 2a9hA | 0.6124 | 0.6239 |
| 2akhA | 0.3107 | 0.3312 |
| 2akhB | 0.5311 | 0.3755 |
| 2bg9A | 0.3933 | 0.2753 |
| 2bl2A | 0.8082 | 0.8331 |
| 2cpbA | 0.2452 | 0.4598 |
| 2d57A | 0.6245 | 0.715 |
| 2ervA | 0.7336 | 0.6133 |

| | | |
|---|---|---|
| 2evuA | 0.6434 | 0.7779 |
| 2f1cX | 0.6168 | 0.6544 |
| 2f93B | 0.4896 | 0.343 |
| 2f95B | 0.3856 | 0.3799 |
| 2fynB | 0.5135 | 0.4227 |
| 2ge4A | 0.562 | 0.6059 |
| 2gfpA | 0.4583 | 0.4858 |
| 2gr7A | 0.3878 | 0.2875 |
| 2gr8A | 0.4775 | 0.4445 |
| 2h8aA | 0.4811 | 0.4087 |
| 2h8pC | 0.7068 | 0.568 |
| 2hdfA | 0.7489 | 0.8037 |
| 2ibzG | 0.2953 | 0.2729 |
| 2ibzI | 0.4438 | 0.3764 |
| 2iubA | 0.4246 | 0.6307 |
| 2j58A | 0.2708 | 0.2477 |
| 2j7aC | 0.2457 | 0.3366 |
| 2jafA | 0.8189 | 0.7918 |
| 2jlnA | 0.6831 | 0.338 |
| 2jo1A | 0.3378 | 0.3925 |
| 2jp3A | 0.3518 | 0.2967 |
| 2k0lA | 0.4969 | 0.5511 |
| 2k21A | 0.238 | 0.2082 |
| 2k73A | 0.6629 | 0.6078 |
| 2k9pA | 0.4038 | 0.3949 |
| 2kluA | 0.3387 | 0.3251 |
| 2kogA | 0.2466 | 0.2278 |
| 2ks9A | 0.6674 | 0.7056 |
| 2ksdA | 0.2958 | 0.3225 |
| 2kseA | 0.3718 | 0.3121 |
| 2ksfA | 0.3109 | 0.3174 |
| 2ksrA | 0.658 | 0.6323 |
| 2kyhA | 0.5838 | 0.4183 |
| 2l35A | 0.435 | 0.3813 |
| 2l8sA | 0.4896 | 0.4847 |
| 2lckA | 0.5268 | 0.5581 |
| 2lhfA | 0.5887 | 0.6081 |
| 2lkgA | 0.3927 | 0.3744 |
| 2llyA | 0.5247 | 0.5304 |
| 2lmeA | 0.3616 | 0.5203 |
| 2lnlA | 0.4934 | 0.4946 |

| | | |
|---|---|---|
| 2lomA | 0.2924 | 0.2861 |
| 2loqA | 0.2092 | 0.1977 |
| 2lorA | 0.3229 | 0.3104 |
| 2losA | 0.2545 | 0.2579 |
| 2lotA | 0.3319 | 0.2867 |
| 2lp1A | 0.3288 | 0.3181 |
| 2m0qA | 0.2187 | 0.2923 |
| 2m20A | 0.3974 | 0.4096 |
| 2m67A | 0.2536 | 0.2548 |
| 2m6bA | 0.4999 | 0.4726 |
| 2m7gA | 0.3132 | 0.3586 |
| 2m8rA | 0.2885 | 0.2379 |
| 2mafA | 0.4911 | 0.4976 |
| 2mfrA | 0.3819 | 0.4094 |
| 2mgyA | 0.5898 | 0.5636 |
| 2mm8A | 0.1479 | 0.1544 |
| 2mmuA | 0.4418 | 0.4572 |
| 2mn6A | 0.4084 | 0.4014 |
| 2mpnA | 0.3132 | 0.3182 |
| 2mxbA | 0.4022 | 0.6764 |
| 2n4xA | 0.199 | 0.2566 |
| 2n6lA | 0.4541 | 0.4381 |
| 2n7qA | 0.5217 | 0.5783 |
| 2nmrA | 0.6475 | 0.7519 |
| 2nq2A | 0.8088 | 0.8165 |
| 2nr9A | 0.7935 | 0.7991 |
| 2nrgA | 0.289 | 0.3023 |
| 2o01F | 0.2991 | 0.3315 |
| 2oarA | 0.2649 | 0.2944 |
| 2pnoA | 0.5729 | 0.7261 |
| 2q67A | 0.6807 | 0.6934 |
| 2q7mA | 0.6188 | 0.747 |
| 2qomA | 0.7164 | 0.7422 |
| 2r6gF | 0.3302 | 0.2109 |
| 2r6gG | 0.4308 | 0.7224 |
| 2vpwC | 0.6364 | 0.7857 |
| 2w1pA | 0.6052 | 0.4157 |
| 2wjqA | 0.6449 | 0.3183 |
| 2wpdJ | 0.6637 | 0.6021 |
| 2wpvB | 0.3113 | 0.3071 |
| 2wsc1 | 0.2946 | 0.2357 |

| | | |
|---|---|---|
| 2wsc3 | 0.2647 | 0.2373 |
| 2wscF | 0.3147 | 0.324 |
| 2wscG | 0.2313 | 0.193 |
| 2wscH | 0.2638 | 0.1855 |
| 2wscK | 0.266 | 0.2404 |
| 2wscL | 0.385 | 0.4405 |
| 2wswA | 0.7736 | 0.7599 |
| 2wwbB | 0.2532 | 0.3536 |
| 2wwbC | 0.4406 | 0.4106 |
| 2x4mA | 0.7129 | 0.335 |
| 2xq2A | 0.5796 | 0.3434 |
| 2xutA | 0.7523 | 0.7454 |
| 2y5yA | 0.7956 | 0.7817 |
| 2y69D | 0.2917 | 0.2641 |
| 2y69G | 0.2654 | 0.2445 |
| 2y69I | 0.3457 | 0.4361 |
| 2y69J | 0.407 | 0.4419 |
| 2y69K | 0.3978 | 0.3867 |
| 2y69L | 0.4056 | 0.4621 |
| 2y69M | 0.426 | 0.4614 |
| 2yevB | 0.5969 | 0.4294 |
| 2yevC | 0.7111 | 0.6005 |
| 2yiuA | 0.7255 | 0.5086 |
| 2ynkA | 0.6681 | 0.2935 |
| 2z73A | 0.5899 | 0.2948 |
| 2ziyA | 0.3481 | 0.7057 |
| 2zjsE | 0.6019 | 0.4842 |
| 2zxeB | 0.5177 | 0.2496 |
| 2zxeG | 0.4656 | 0.5955 |
| 3a2sX | 0.5921 | 0.6867 |
| 3a7kA | 0.8192 | 0.7488 |
| 3anzA | 0.3625 | 0.4512 |
| 3b4rA | 0.6689 | 0.569 |
| 3b5dA | 0.625 | 0.4111 |
| 3b9wA | 0.5918 | 0.4255 |
| 3bryA | 0.7026 | 0.3173 |
| 3chxB | 0.4165 | 0.2232 |
| 3chxC | 0.7087 | 0.4695 |
| 3cn5A | 0.6137 | 0.5536 |
| 3cx5C | 0.7911 | 0.7107 |
| 3d31C | 0.5306 | 0.6618 |

| | | |
|---|---|---|
| 3ddlA | 0.7723 | 0.7807 |
| 3dhwA | 0.6188 | 0.782 |
| 3dinE | 0.3319 | 0.3446 |
| 3dl8C | 0.573 | 0.77 |
| 3dl8E | 0.3563 | 0.3891 |
| 3dwoX | 0.7049 | 0.645 |
| 3dwwA | 0.5136 | 0.4331 |
| 3dzmA | 0.3355 | 0.2757 |
| 3effK | 0.4821 | 0.5159 |
| 3eh3A | 0.7495 | 0.6252 |
| 3ejzA | 0.8617 | 0.7434 |
| 3emnX | 0.6038 | 0.5523 |
| 3emoA | 0.2366 | 0.2862 |
| 3fhhA | 0.7555 | 0.7939 |
| 3fidA | 0.7199 | 0.2853 |
| 3g67A | 0.2347 | 0.4816 |
| 3gi8C | 0.7914 | 0.7468 |
| 3hd6A | 0.5817 | 0.4131 |
| 3hw9A | 0.532 | 0.291 |
| 3iyzA | 0.6 | 0.7739 |
| 3iz1A | 0.5221 | 0.6045 |
| 3j08A | 0.2722 | 0.3292 |
| 3j1zP | 0.5173 | 0.5538 |
| 3j9tR | 0.7322 | 0.7646 |
| 3jbrE | 0.5363 | 0.5389 |
| 3jcuD | 0.2071 | 0.2185 |
| 3jcuH | 0.4147 | 0.434 |
| 3jcuK | 0.3805 | 0.3667 |
| 3jcuR | 0.5059 | 0.4017 |
| 3jcuS | 0.4908 | 0.3986 |
| 3jcuW | 0.4565 | 0.4223 |
| 3jcuX | 0.415 | 0.6773 |
| 3jcuZ | 0.676 | 0.6279 |
| 3jycA | 0.3133 | 0.294 |
| 3k3fA | 0.809 | 0.7263 |
| 3kj6A | 0.7528 | 0.8199 |
| 3kp9A | 0.425 | 0.4483 |
| 3kvnA | 0.4614 | 0.3454 |
| 3l1lA | 0.7796 | 0.8534 |
| 3lnmB | 0.306 | 0.3101 |
| 3lw54 | 0.3932 | 0.38 |

| | | |
|---|---|---|
| 3lw5H | 0.2353 | 0.2484 |
| 3m71A | 0.8324 | 0.8828 |
| 3mk7A | 0.7962 | 0.8205 |
| 3mk7B | 0.5235 | 0.2603 |
| 3mk7C | 0.2503 | 0.2057 |
| 3mktA | 0.7518 | 0.7786 |
| 3mp7A | 0.4221 | 0.4096 |
| 3mp7B | 0.3917 | 0.4711 |
| 3njtA | 0.438 | 0.2536 |
| 3nymA | 0.356 | 0.4003 |
| 3o0rB | 0.8283 | 0.8475 |
| 3o7pA | 0.7098 | 0.7546 |
| 3ohnA | 0.4836 | 0.2714 |
| 3orgA | 0.5042 | 0.4766 |
| 3oufA | 0.7368 | 0.5915 |
| 3p5nA | 0.7939 | 0.8294 |
| 3pjsK | 0.5094 | 0.5404 |
| 3pjzA | 0.7995 | 0.7571 |
| 3pwhA | 0.7371 | 0.8031 |
| 3q7kA | 0.6235 | 0.7033 |
| 3qe7A | 0.7341 | 0.5936 |
| 3qnqA | 0.4609 | 0.4555 |
| 3qraA | 0.727 | 0.7347 |
| 3rbzA | 0.4813 | 0.4529 |
| 3rgwS | 0.7305 | 0.3446 |
| 3rkoA | 0.4705 | 0.4831 |
| 3rkoB | 0.7773 | 0.8133 |
| 3rkoC | 0.8831 | 0.8926 |
| 3rkoD | 0.7656 | 0.8324 |
| 3rkoF | 0.3956 | 0.4901 |
| 3rkoG | 0.628 | 0.7313 |
| 3s0xA | 0.502 | 0.4617 |
| 3sljA | 0.7694 | 0.5647 |
| 3sybA | 0.6238 | 0.2713 |
| 3tijA | 0.6332 | 0.5008 |
| 3tx3A | 0.6225 | 0.7126 |
| 3udcA | 0.3595 | 0.3383 |
| 3ug9A | 0.5215 | 0.5921 |
| 3ukmA | 0.447 | 0.3512 |
| 3um7A | 0.3764 | 0.4527 |
| 3uq7A | 0.4548 | 0.3376 |

| | | |
|---|---|---|
| 3ux4A | 0.658 | 0.6959 |
| 3v2wA | 0.4069 | 0.5358 |
| 3v5sA | 0.7771 | 0.714 |
| 3vmqA | 0.7461 | 0.5257 |
| 3vouA | 0.3278 | 0.5042 |
| 3vr8C | 0.3862 | 0.5007 |
| 3vr8D | 0.5395 | 0.4983 |
| 3vwiA | 0.7459 | 0.6482 |
| 3wdoA | 0.6548 | 0.4826 |
| 3wmfA | 0.3968 | 0.3838 |
| 3wmm1 | 0.4146 | 0.522 |
| 3wmmM | 0.4311 | 0.476 |
| 3wo7A | 0.6794 | 0.5283 |
| 3wvfA | 0.4714 | 0.2973 |
| 3wxvA | 0.7971 | 0.7177 |
| 3x29A | 0.7871 | 0.8011 |
| 3x2rA | 0.4121 | 0.411 |
| 3x3bA | 0.7106 | 0.5401 |
| 3ze3A | 0.6792 | 0.539 |
| 3zevA | 0.7579 | 0.8241 |
| 3zjzA | 0.644 | 0.7247 |
| 3zk1A | 0.533 | 0.5287 |
| 3zuxA | 0.7555 | 0.6698 |
| 4a2nB | 0.4887 | 0.5256 |
| 4atvA | 0.788 | 0.7374 |
| 4aw6A | 0.8146 | 0.5563 |
| 4b4aA | 0.5988 | 0.6279 |
| 4bemJ | 0.6619 | 0.7172 |
| 4bgnA | 0.435 | 0.457 |
| 4bog3 | 0.3463 | 0.2481 |
| 4bpmA | 0.5268 | 0.6695 |
| 4bwzA | 0.6707 | 0.7494 |
| 4c9jA | 0.6974 | 0.7378 |
| 4cadC | 0.6688 | 0.6779 |
| 4cfgA | 0.1658 | 0.2822 |
| 4chvA | 0.3095 | 0.3621 |
| 4cskA | 0.6 | 0.6345 |
| 4czbA | 0.7274 | 0.8492 |
| 4d5bA | 0.4624 | 0.236 |
| 4d6tD | 0.6198 | 0.4858 |
| 4d6tG | 0.3183 | 0.3553 |

| | | |
|---|---|---|
| 4d6tJ | 0.4831 | 0.3882 |
| 4d6uD | 0.5639 | 0.3104 |
| 4djiA | 0.722 | 0.4747 |
| 4dojA | 0.7083 | 0.648 |
| 4dveA | 0.7842 | 0.736 |
| 4dxwA | 0.4014 | 0.4211 |
| 4e1tA | 0.7204 | 0.7013 |
| 4ea3A | 0.8219 | 0.668 |
| 4ezcA | 0.6454 | 0.6065 |
| 4f35A | 0.6594 | 0.6328 |
| 4f4lA | 0.6626 | 0.7303 |
| 4fqeA | 0.6701 | 0.7767 |
| 4fuvA | 0.5398 | 0.3823 |
| 4g1uA | 0.8449 | 0.3981 |
| 4g7vS | 0.6707 | 0.7479 |
| 4g80I | 0.6328 | 0.669 |
| 4gbyA | 0.87 | 0.8771 |
| 4gd3A | 0.6493 | 0.6337 |
| 4gx5A | 0.379 | 0.2974 |
| 4gycB | 0.4636 | 0.3227 |
| 4h33A | 0.5708 | 0.6844 |
| 4he8A | 0.5192 | 0.4227 |
| 4he8C | 0.7757 | 0.6369 |
| 4he8D | 0.4096 | 0.4894 |
| 4hkrA | 0.4304 | 0.5478 |
| 4hqjE | 0.6997 | 0.6634 |
| 4httA | 0.3703 | 0.4451 |
| 4huqS | 0.7089 | 0.7746 |
| 4huqT | 0.4824 | 0.4769 |
| 4hw9A | 0.2708 | 0.3283 |
| 4hycA | 0.709 | 0.4693 |
| 4hyoA | 0.7659 | 0.6789 |
| 4hzuS | 0.7682 | 0.8406 |
| 4iffA | 0.3658 | 0.4807 |
| 4il3A | 0.8274 | 0.5235 |
| 4in5H | 0.3293 | 0.4018 |
| 4in5L | 0.4116 | 0.5423 |
| 4j05A | 0.8409 | 0.8879 |
| 4j72A | 0.7775 | 0.7985 |
| 4j7cI | 0.7474 | 0.6315 |
| 4jkvA | 0.4171 | 0.5273 |

| | | |
|---|---|---|
| 4k1cA | 0.6228 | 0.6158 |
| 4kjrA | 0.8065 | 0.7262 |
| 4knfA | 0.8399 | 0.7806 |
| 4kppA | 0.6619 | 0.5542 |
| 4kt0F | 0.261 | 0.3246 |
| 4kt0K | 0.5048 | 0.4736 |
| 4ky0A | 0.635 | 0.4199 |
| 4l6rA | 0.5337 | 0.5422 |
| 4l6v6 | 0.3761 | 0.3959 |
| 4l6v8 | 0.5395 | 0.5715 |
| 4ltoA | 0.5488 | 0.5733 |
| 4m58A | 0.7495 | 0.7513 |
| 4m64A | 0.7119 | 0.473 |
| 4mbsA | 0.6698 | 0.6489 |
| 4meeA | 0.6672 | 0.4395 |
| 4mndA | 0.4946 | 0.318 |
| 4mqsA | 0.7275 | 0.8229 |
| 4mt4A | 0.4275 | 0.4304 |
| 4n74A | 0.7335 | 0.2806 |
| 4n75A | 0.6768 | 0.5265 |
| 4njnA | 0.7974 | 0.8275 |
| 4nppA | 0.4635 | 0.3358 |
| 4ntjA | 0.5233 | 0.5783 |
| 4nykA | 0.5547 | 0.2433 |
| 4o6mA | 0.4872 | 0.4464 |
| 4o6yA | 0.7727 | 0.7953 |
| 4o9pA | 0.5137 | 0.5024 |
| 4o9pB | 0.5975 | 0.3882 |
| 4o9uB | 0.4712 | 0.3253 |
| 4od4A | 0.8394 | 0.848 |
| 4ogqC | 0.2345 | 0.2838 |
| 4oh3A | 0.636 | 0.7527 |
| 4oo9A | 0.4832 | 0.4754 |
| 4or2A | 0.5175 | 0.6078 |
| 4p6vB | 0.6718 | 0.7323 |
| 4p6vC | 0.7186 | 0.592 |
| 4p6vD | 0.5634 | 0.7646 |
| 4p6vE | 0.7958 | 0.758 |
| 4p6vF | 0.6329 | 0.4363 |
| 4p79A | 0.7562 | 0.822 |
| 4pgrA | 0.6699 | 0.6633 |

| | | |
|---|---|---|
| 4phzA | 0.3107 | 0.2097 |
| 4pirA | 0.3036 | 0.3284 |
| 4px7A | 0.4935 | 0.4682 |
| 4q2eA | 0.5953 | 0.6021 |
| 4qncA | 0.8424 | 0.7278 |
| 4qndA | 0.6597 | 0.6323 |
| 4qtnA | 0.6382 | 0.7773 |
| 4quvA | 0.6427 | 0.6513 |
| 4r1iA | 0.4466 | 0.5739 |
| 4rdqA | 0.3845 | 0.3497 |
| 4rfsS | 0.7466 | 0.6987 |
| 4ri2A | 0.3579 | 0.4477 |
| 4rjwA | 0.5297 | 0.4822 |
| 4rl8A | 0.7471 | 0.7557 |
| 4rl9A | 0.4503 | 0.3291 |
| 4rlcA | 0.6894 | 0.8185 |
| 4rngA | 0.734 | 0.6903 |
| 4rp8A | 0.5488 | 0.2837 |
| 4ryiA | 0.7922 | 0.7285 |
| 4s0vA | 0.5354 | 0.3733 |
| 4tkrA | 0.7265 | 0.6963 |
| 4tq3A | 0.7081 | 0.6874 |
| 4tquM | 0.4755 | 0.7068 |
| 4tquN | 0.3466 | 0.6698 |
| 4twkA | 0.31 | 0.3344 |
| 4u15A | 0.5867 | 0.5872 |
| 4u4tA | 0.7649 | 0.8076 |
| 4u9lA | 0.6021 | 0.651 |
| 4uc1A | 0.8659 | 0.7907 |
| 4us3A | 0.8744 | 0.7706 |
| 4v1fA | 0.6338 | 0.7732 |
| 4wd7A | 0.7395 | 0.658 |
| 4wgvA | 0.7245 | 0.7758 |
| 4wmzA | 0.7877 | 0.7337 |
| 4x5mA | 0.6624 | 0.6395 |
| 4xk83 | 0.4518 | 0.4564 |
| 4xnkA | 0.2699 | 0.5554 |
| 4xnvA | 0.636 | 0.6519 |
| 4xu4A | 0.7309 | 0.3846 |
| 4xxjA | 0.8275 | 0.6595 |
| 4xydB | 0.5457 | 0.4872 |

| | | |
|---|---|---|
| 4y25A | 0.781 | 0.6107 |
| 4y28G | 0.4373 | 0.3723 |
| 4y28K | 0.5202 | 0.4484 |
| 4y28L | 0.4904 | 0.5705 |
| 4y7jA | 0.4532 | 0.463 |
| 4ymkA | 0.6606 | 0.3382 |
| 4ymsC | 0.4637 | 0.7708 |
| 4ytpC | 0.3568 | 0.537 |
| 4ytpD | 0.5864 | 0.542 |
| 4z34A | 0.5615 | 0.6433 |
| 4z3nA | 0.757 | 0.7924 |
| 4z7fA | 0.7618 | 0.7758 |
| 4zp0A | 0.8243 | 0.8488 |
| 4zr0A | 0.5228 | 0.3276 |
| 4zr1A | 0.4574 | 0.387 |
| 4zw9A | 0.8757 | 0.5697 |
| 5a1sA | 0.4329 | 0.3952 |
| 5a40A | 0.6244 | 0.7976 |
| 5a63C | 0.7314 | 0.6612 |
| 5a63D | 0.5658 | 0.5393 |
| 5a6eB | 0.8291 | 0.7337 |
| 5abbZ | 0.4605 | 0.411 |
| 5araT | 0.2078 | 0.3324 |
| 5araW | 0.4414 | 0.5197 |
| 5awwG | 0.6435 | 0.6673 |
| 5awwY | 0.5922 | 0.4242 |
| 5awzA | 0.7942 | 0.8204 |
| 5aymA | 0.8081 | 0.7657 |
| 5azbA | 0.6844 | 0.7149 |
| 5bwkE | 0.4651 | 0.4785 |
| 5c6oA | 0.8266 | 0.8493 |
| 5c8jI | 0.4038 | 0.3291 |
| 5cfbA | 0.3658 | 0.3602 |
| 5ctgA | 0.798 | 0.7663 |
| 5d0yA | 0.735 | 0.7649 |
| 5dirA | 0.7698 | 0.594 |
| 5doqA | 0.8274 | 0.6941 |
| 5doqB | 0.7743 | 0.4748 |
| 5ee7A | 0.5575 | 0.4956 |
| 5ek0A | 0.4517 | 0.3798 |
| 5ekeA | 0.5233 | 0.5223 |

| | | |
|---|---|---|
| 5eulE | 0.5052 | 0.4719 |
| 5ezmA | 0.445 | 0.4381 |
| 5f1cA | 0.2454 | 0.2858 |
| 5fn2B | 0.3184 | 0.2641 |
| 5gaeh | 0.4653 | 0.3615 |
| 5gaqA | 0.1761 | 0.1734 |
| 5garO | 0.5138 | 0.6123 |
| 5hk1A | 0.6744 | 0.4621 |
| 5i1mV | 0.3702 | 0.3898 |
| 5i20A | 0.7517 | 0.7972 |
| 5i32A | 0.6282 | 0.4813 |
| 5i6cA | 0.5711 | 0.3472 |
| 5i6zA | 0.8654 | 0.821 |
| 5id3A | 0.2839 | 0.2051 |
| 5iofA | 0.7028 | 0.754 |
| 5irxA | 0.2581 | 0.203 |
| 5ivaA | 0.287 | 0.2933 |
| 5iwsA | 0.4699 | 0.486 |
| 5ixmB | 0.453 | 0.7179 |
| 5jagA | 0.8189 | 0.7728 |
| Mean | 0.546479804 | 0.521108627 |
| Fold | 294 | 255 |
| p-Value | | 6.54197E-06 |

**S7 Table.** Target-by-target stagewise reconstruction performance on EVfold dataset for true $C_\beta$–$C_\beta$ contact maps at 8, 10, and 12Å thresholds.

| Targets | 8 Å | | | | | 10 Å | | | | | 12 Å | | | | |
|---|---|---|---|---|---|---|---|---|---|---|---|---|---|---|---|
| | stage 1 | stage 2 | stage 3 | $\Delta_1$ (stage 2-stage 1) | $\Delta_2$ (stage 3-stage 2) | stage 1 | stage 2 | stage 3 | $\Delta_1$ (stage 2-stage 1) | $\Delta_2$ (stage 3-stage 2) | stage 1 | stage 2 | stage 3 | $\Delta_1$ (stage 2-stage 1) | $\Delta_2$ (stage 3-stage 2) |
| 1bkrA | 0.749 | 0.8652 | 0.9031 | 0.1162 | 0.0379 | 0.6087 | 0.8781 | 0.9334 | 0.2694 | 0.0553 | 0.4376 | 0.893 | 0.9541 | 0.4554 | 0.0611 |
| 1e6kA | 0.6059 | 0.7923 | 0.8735 | 0.1864 | 0.0812 | 0.5139 | 0.843 | 0.911 | 0.3291 | 0.068 | 0.4136 | 0.8697 | 0.9179 | 0.4561 | 0.0482 |
| 1f21A | 0.6504 | 0.8178 | 0.8769 | 0.1674 | 0.0591 | 0.5361 | 0.8644 | 0.9214 | 0.3283 | 0.057 | 0.4223 | 0.8832 | 0.9132 | 0.4609 | 0.03 |
| 1g2eA | 0.4311 | 0.6716 | 0.7697 | 0.2405 | 0.0981 | 0.3934 | 0.7598 | 0.8251 | 0.3664 | 0.0653 | 0.3003 | 0.7803 | 0.8692 | 0.48 | 0.0889 |
| 1hzxA | 0.7519 | 0.8705 | 0.9126 | 0.1186 | 0.0421 | 0.6074 | 0.9135 | 0.951 | 0.3061 | 0.0375 | 0.4202 | 0.9311 | 0.953 | 0.5109 | 0.0219 |
| 1oddA | 0.5953 | 0.6706 | 0.7672 | 0.0753 | 0.0966 | 0.4254 | 0.7456 | 0.87 | 0.3202 | 0.1244 | 0.3529 | 0.7378 | 0.8813 | 0.3849 | 0.1435 |
| 1r9hA | 0.4405 | 0.7418 | 0.8207 | 0.3013 | 0.0789 | 0.4371 | 0.8212 | 0.8829 | 0.3841 | 0.0617 | 0.3293 | 0.8461 | 0.8996 | 0.5168 | 0.0535 |
| 1rqmA | 0.5908 | 0.7974 | 0.8402 | 0.2066 | 0.0428 | 0.5443 | 0.8976 | 0.9218 | 0.3533 | 0.0242 | 0.4139 | 0.8937 | 0.9245 | 0.4798 | 0.0308 |
| 1wvnA | 0.4507 | 0.7116 | 0.7808 | 0.2609 | 0.0692 | 0.4141 | 0.7621 | 0.8083 | 0.348 | 0.0462 | 0.3156 | 0.7354 | 0.7846 | 0.4198 | 0.0492 |
| 2hdaA | 0.2412 | 0.5251 | 0.6091 | 0.2839 | 0.084 | 0.2575 | 0.6491 | 0.7728 | 0.3916 | 0.1237 | 0.2576 | 0.6409 | 0.7748 | 0.3833 | 0.1339 |
| 2it6A | 0.5176 | 0.786 | 0.8619 | 0.2684 | 0.0759 | 0.4344 | 0.8228 | 0.9147 | 0.3884 | 0.0919 | 0.3643 | 0.8782 | 0.9253 | 0.5139 | 0.0471 |
| 2o72A | 0.3793 | 0.71 | 0.8067 | 0.3307 | 0.0967 | 0.3709 | 0.7893 | 0.877 | 0.4184 | 0.0877 | 0.3059 | 0.8034 | 0.9031 | 0.4975 | 0.0997 |
| 3tgiE | 0.6413 | 0.8794 | 0.9375 | 0.2381 | 0.0581 | 0.5496 | 0.9196 | 0.9564 | 0.37 | 0.0368 | 0.4423 | 0.9253 | 0.9539 | 0.483 | 0.0286 |
| 5p21A | 0.6113 | 0.8791 | 0.9312 | 0.2678 | 0.0521 | 0.5602 | 0.8956 | 0.9365 | 0.3354 | 0.0409 | 0.4344 | 0.9108 | 0.9558 | 0.4764 | 0.045 |
| 5ptiA | 0.4365 | 0.6378 | 0.7908 | 0.2013 | 0.153 | 0.4046 | 0.7334 | 0.8538 | 0.3288 | 0.1204 | 0.3507 | 0.7283 | 0.8467 | 0.3776 | 0.1184 |
| Mean | 0.53952 | 0.75708 | 0.832126667 | 0.21756 | 0.075046667 | 0.470506667 | 0.819673333 | 0.889073333 | 0.349166667 | 0.0694 | 0.370726667 | 0.83048 | 0.897133333 | 0.459753333 | 0.066653333 |

**S8 Table.** Target-by-target stagewise reconstruction performance on EVfold dataset for true $C_\alpha$–$C_\alpha$ contact maps at 8, 10, and 12Å thresholds.

| Targets | 8 Å | | | | | 10 Å | | | | | 12 Å | | | | |
|---|---|---|---|---|---|---|---|---|---|---|---|---|---|---|---|
| | stage 1 | stage 2 | stage 3 | $\Delta_1$ (stage 2-stage 1) | $\Delta_2$ (stage 3-stage 2) | stage 1 | stage 2 | stage 3 | $\Delta_1$ (stage 2-stage 1) | $\Delta_2$ (stage 3-stage 2) | stage 1 | stage 2 | stage 3 | $\Delta_1$ (stage 2-stage 1) | $\Delta_2$ (stage 3-stage 2) |
| 1bkrA | 0.6891 | 0.8324 | 0.9021 | 0.1433 | 0.0697 | 0.6525 | 0.927 | 0.9645 | 0.2745 | 0.0375 | 0.4458 | 0.9571 | 0.9736 | 0.5113 | 0.0165 |
| 1e6kA | 0.3945 | 0.7814 | 0.8244 | 0.3869 | 0.043 | 0.5539 | 0.8892 | 0.9145 | 0.3353 | 0.0253 | 0.4318 | 0.9377 | 0.9233 | 0.5059 | -0.0144 |
| 1f21A | 0.6073 | 0.7883 | 0.8351 | 0.181 | 0.0468 | 0.5736 | 0.8887 | 0.9191 | 0.3151 | 0.0304 | 0.4469 | 0.9125 | 0.9603 | 0.4656 | 0.0478 |
| 1g2eA | 0.4059 | 0.6749 | 0.7443 | 0.269 | 0.0694 | 0.4578 | 0.8586 | 0.8554 | 0.4008 | -0.0032 | 0.3396 | 0.8229 | 0.8927 | 0.4833 | 0.0698 |
| 1hzxA | 0.7088 | 0.8073 | 0.8813 | 0.0985 | 0.074 | 0.6171 | 0.925 | 0.9634 | 0.3079 | 0.0384 | 0.4499 | 0.9662 | 0.9764 | 0.5163 | 0.0102 |
| 1oddA | 0.5462 | 0.6849 | 0.8033 | 0.1387 | 0.1184 | 0.4965 | 0.8008 | 0.8896 | 0.3043 | 0.0888 | 0.3099 | 0.8182 | 0.9052 | 0.5083 | 0.087 |
| 1r9hA | 0.3951 | 0.296 | 0.3249 | -0.0991 | 0.0289 | 0.4843 | 0.8394 | 0.9155 | 0.3551 | 0.0761 | 0.3517 | 0.8582 | 0.94 | 0.5065 | 0.0818 |
| 1rqmA | 0.4757 | 0.734 | 0.8283 | 0.2583 | 0.0943 | 0.5593 | 0.9123 | 0.9501 | 0.353 | 0.0378 | 0.4023 | 0.9455 | 0.9539 | 0.5432 | 0.0084 |
| 1wvnA | 0.3695 | 0.5584 | 0.7416 | 0.1889 | 0.1832 | 0.4363 | 0.7911 | 0.7527 | 0.3548 | -0.0384 | 0.323 | 0.8093 | 0.8395 | 0.4863 | 0.0302 |
| 2hdaA | 0.1846 | 0.2494 | 0.4379 | 0.0648 | 0.1885 | 0.2997 | 0.6601 | 0.7487 | 0.3604 | 0.0886 | 0.2236 | 0.6655 | 0.7685 | 0.4419 | 0.103 |
| 2it6A | 0.5121 | 0.6736 | 0.7843 | 0.1615 | 0.1107 | 0.4454 | 0.8744 | 0.9395 | 0.429 | 0.0651 | 0.3685 | 0.9095 | 0.9538 | 0.541 | 0.0443 |
| 2o72A | 0.3943 | 0.6761 | 0.7665 | 0.2818 | 0.0904 | 0.2303 | 0.9003 | 0.9222 | 0.67 | 0.0219 | 0.2236 | 0.8865 | 0.915 | 0.6629 | 0.0285 |
| 3tgiE | 0.6393 | 0.8758 | 0.9341 | 0.2365 | 0.0583 | 0.5616 | 0.9521 | 0.9718 | 0.3905 | 0.0197 | 0.2473 | 0.9666 | 0.9757 | 0.7193 | 0.0091 |
| 5p21A | 0.5273 | 0.7845 | 0.8533 | 0.2572 | 0.0688 | 0.5523 | 0.9362 | 0.9502 | 0.3839 | 0.014 | 0.4462 | 0.9357 | 0.97 | 0.4895 | 0.0343 |
| 5ptiA | 0.4433 | 0.6765 | 0.8368 | 0.2332 | 0.1603 | 0.4022 | 0.7791 | 0.8629 | 0.3769 | 0.0838 | 0.3544 | 0.7061 | 0.8131 | 0.3517 | 0.107 |
| Mean | 0.4862 | 0.6729 | 0.766546667 | 0.1867 | 0.093646667 | 0.488186667 | 0.862286667 | 0.90134 | 0.3741 | 0.039053333 | 0.357633333 | 0.873166667 | 0.9174 | 0.515533333 | 0.044233333 |

**S9 Table.** Target-by-target stagewise recovery of secondary structure topology on EVfold dataset for true Cβ–Cβ contact maps at 8, 10, and 12Å thresholds.

| Target | 8 Å | | | | | | 10 Å | | | | | | 12 Å | | | | | |
|---|---|---|---|---|---|---|---|---|---|---|---|---|---|---|---|---|---|---|
| | Stage 1 | | Stage 2 | | Stage 3 | | Stage 1 | | Stage 2 | | Stage 3 | | Stage 1 | | Stage 2 | | Stage 3 | |
| | $Q_H$ | $Q_E$ | $Q_H$ | $Q_E$ | $Q_H$ | $Q_E$ | $Q_H$ | $Q_E$ | $Q_H$ | $Q_E$ | $Q_H$ | $Q_E$ | $Q_H$ | $Q_E$ | $Q_H$ | $Q_E$ | $Q_H$ | $Q_E$ |
| 1bkrA | 37.14285714 | | 74.28571429 | | 98.57142857 | | 22.85714286 | | 72.85714286 | | 98.57142857 | | 38.57142857 | | 67.14285714 | | 100 | |
| 1e6kA | 37.25490196 | 0 | 70.58823529 | 20 | 92.15686275 | 85 | 17.64705882 | 0 | 23.52941176 | 0 | 96.07843137 | 85 | 9.803921569 | 0 | 45.09803922 | 10 | 100 | 55 |
| 1f21A | 29.31034483 | 4.166666667 | 63.79310345 | 27.08333333 | 98.27586207 | 60.41666667 | 18.96551724 | 0 | 41.37931034 | 31.25 | 96.55172414 | 58.33333333 | 36.20689655 | 2.083333333 | 24.13793103 | 18.75 | 96.55172414 | 64.58333333 |
| 1g2eA | 14.28571429 | 0 | 80.95238095 | 0 | 100 | 36 | 0 | 0 | 71.42857143 | 16 | 100 | 24 | 23.80952381 | 8 | 47.61904762 | 0 | 100 | 44 |
| 1hzxA | 1.104972376 | 0 | 60.22099448 | 0 | 86.1878453 | 25 | 14.36464088 | 0 | 49.17127072 | 37.5 | 86.74033149 | 12.5 | 7.73480663 | 12.5 | 52.48618785 | 12.5 | 88.95027624 | 12.5 |
| 1oddA | 9.375 | 0 | 59.375 | 0 | 100 | 42.85714286 | 9.375 | 0 | 53.125 | 0 | 96.875 | 71.42857143 | 15.625 | 0 | 84.375 | 0 | 100 | 57.14285714 |
| 1r9hA | 0 | 0 | 42.85714286 | 0 | 100 | 77.77777778 | 0 | 0 | 35.71428571 | 5.555555556 | 78.57142857 | 80.55555556 | 0 | 11.11111111 | 71.42857143 | 38.88888889 | 71.42857143 | 72.22222222 |
| 1rqmA | 10.25641026 | 0 | 51.28205128 | 24 | 89.74358974 | 68 | 30.76923077 | 0 | 58.97435897 | 24 | 87.17948718 | 80 | 0 | 0 | 61.53846154 | 20 | 87.17948718 | 80 |
| 1wvnA | 19.35483871 | 0 | 74.19354839 | 17.64705882 | 100 | 70.58823529 | 29.03225806 | 5.882352941 | 87.09677419 | 23.52941176 | 100 | 82.35294118 | 0 | 0 | 41.93548387 | 23.52941176 | 100 | 82.35294118 |
| 2hdaA | | 5.263157895 | | 0 | | 47.36842105 | | 5.263157895 | | 21.05263158 | | 68.42105263 | | 0 | | 26.31578947 | | 89.47368421 |
| 2it6A | 11.11111111 | 0 | 66.66666667 | 8.823529412 | 100 | 58.82352941 | 0 | 11.76470588 | 48.14814815 | 5.882352941 | 100 | 58.82352941 | 0 | 2.941176471 | 37.03703704 | 29.41176471 | 96.2962963 | 70.58823529 |
| 2o72A | | 2.127659574 | | 27.65957447 | | 87.23404255 | | 8.510638298 | | 27.65957447 | | 61.70212766 | | 0 | | 12.76595745 | | 80.85106383 |
| 3tgiE | 0 | 2.631578947 | 0 | 13.15789474 | 57.14285714 | 71.05263158 | 42.85714286 | 2.631578947 | 0 | 13.15789474 | 100 | 77.63157895 | 0 | 1.315789474 | 42.85714286 | 26.31578947 | 57.14285714 | 81.57894737 |
| 5p21A | 0 | 0 | 75.80645161 | 5.128205128 | 98.38709677 | 74.35897436 | 12.90322581 | 0 | 50 | 15.38461538 | 98.38709677 | 71.79487179 | 17.74193548 | 0 | 53.22580645 | 28.20512821 | 98.38709677 | 87.17948718 |
| 5ptiA | 0 | 0 | 50 | 40 | 100 | 60 | 0 | 0 | 100 | 26.66666667 | 100 | 86.66666667 | 0 | 20 | 37.5 | 40 | 100 | 100 |
| Mean | 13.01508851 | 1.013504506 | 59.23240687 | 13.10711399 | 93.8819648 | 61.74838725 | 15.29009364 | 2.432316712 | 53.18648263 | 17.68847879 | 95.30422524 | 65.65787347 | 11.49950097 | 4.139386456 | 51.26012046 | 20.47733785 | 91.99510071 | 69.8194837 |

**S10 Table.** Target-by-target stagewise recovery of secondary structure topology on EVfold dataset for true $C_\alpha$–$C_\alpha$ contact maps at 8, 10, and 12Å thresholds.

| Target | 8 Å | | | | | | 10 Å | | | | | | 12 Å | | | | | |
|---|---|---|---|---|---|---|---|---|---|---|---|---|---|---|---|---|---|---|
| | Stage 1 | | Stage 2 | | Stage 3 | | Stage 1 | | Stage 2 | | Stage 3 | | Stage 1 | | Stage 2 | | Stage 3 | |
| | $Q_H$ | $Q_E$ | $Q_H$ | $Q_E$ | $Q_H$ | $Q_E$ | $Q_H$ | $Q_E$ | $Q_H$ | $Q_E$ | $Q_H$ | $Q_E$ | $Q_H$ | $Q_E$ | $Q_H$ | $Q_E$ | $Q_H$ | $Q_E$ |
| 1bkrA | 18.57142857 | | 72.85714286 | | 95.71428571 | | 21.42857143 | | 77.14285714 | | 97.14285714 | | 28.57142857 | | 75.71428571 | | 97.14285714 | |
| 1e6kA | 5.882352941 | 0 | 68.62745098 | 0 | 94.11764706 | 80 | 5.882352941 | 0 | 68.62745098 | 30 | 96.07843137 | 45 | 1.960784314 | 0 | 80.39215686 | 20 | 96.07843137 | 40 |
| 1f21A | 0 | 0 | 63.79310345 | 16.66666667 | 94.82758621 | 68.75 | 32.75862069 | 0 | 86.20689655 | 47.91666667 | 96.55172414 | 62.5 | 13.79310345 | 0 | 86.20689655 | 29.16666667 | 100 | 75 |
| 1g2eA | 14.28571429 | 0 | 71.42857143 | 0 | 95.23809524 | 8 | 0 | 4 | 76.19047619 | 0 | 100 | 24 | 0 | 0 | 90.47619048 | 0 | 100 | 16 |
| 1hzxA | 17.12707182 | 0 | 62.98342541 | 0 | 83.97790055 | 0 | 12.15469613 | 25 | 70.71823204 | 0 | 87.84530387 | 0 | 13.8121547 | 0 | 78.45303867 | 0 | 87.29281768 | 12.5 |
| 1oddA | 15.625 | 0 | 65.625 | 0 | 96.875 | 100 | 0 | 0 | 93.75 | 0 | 96.875 | 71.42857143 | 0 | 0 | 90.625 | 28.57142857 | 100 | 71.42857143 |
| 1r9hA | 0 | 0 | 28.57142857 | 0 | 64.28571429 | 16.66666667 | 28.57142857 | 0 | 57.14285714 | 11.11111111 | 78.57142857 | 91.66666667 | 0 | 2.777777778 | 57.14285714 | 16.66666667 | 78.57142857 | 80.55555556 |
| 1rqmA | 0 | 0 | 43.58974359 | 36 | 89.74358974 | 88 | 28.20512821 | 8 | 66.66666667 | 24 | 92.30769231 | 88 | 30.76923077 | 0 | 71.79487179 | 32 | 89.74358974 | 72 |
| 1wvnA | 22.58064516 | 0 | 70.96774194 | 0 | 100 | 47.05882353 | 0 | 0 | 87.09677419 | 0 | 100 | 70.58823529 | 0 | 11.76470588 | 77.41935484 | 23.52941176 | 100 | 82.35294118 |
| 2hdaA | | 0 | | 0 | | 10.52631579 | | 0 | | 21.05263158 | | 63.15789474 | | 0 | | 0 | | 89.47368421 |
| 2it6A | 11.11111111 | 0 | 70.37037037 | 0 | 85.18518519 | 23.52941176 | 0 | 5.882352941 | 81.48148148 | 23.52941176 | 96.2962963 | 58.82352941 | 0 | 5.882352941 | 48.14814815 | 0 | 100 | 58.82352941 |
| 2o72A | | 0 | | 17.0212766 | | 74.46808511 | | 4.255319149 | | 12.76595745 | | 80.85106383 | | 0 | | 31.91489362 | | 65.95744681 |
| 3tgiE | 0 | 0 | 42.85714286 | 23.68421053 | 100 | 67.10526316 | 0 | 5.263157895 | 57.14285714 | 32.89473684 | 100 | 81.57894737 | 0 | 1.315789474 | 42.85714286 | 48.68421053 | 100 | 76.31578947 |
| 5p21A | 11.29032258 | 0 | 69.35483871 | 30.76923077 | 100 | 66.66666667 | 14.51612903 | 2.564102564 | 85.48387097 | 35.8974359 | 98.38709677 | 58.97435897 | 17.74193548 | 0 | 79.03225806 | 38.46153846 | 100 | 87.17948718 |
| 5ptiA | 12.5 | 0 | 87.5 | 26.66666667 | 100 | 86.66666667 | 0 | 0 | 87.5 | 26.66666667 | 100 | 86.66666667 | 0 | 0 | 62.5 | 13.33333333 | 100 | 86.66666667 |
| Mean | 9.921049729 | 0 | 62.9635354 | 10.77200366 | 92.30500031 | 52.67413567 | 11.03976362 | 3.926066611 | 76.55003235 | 18.988187 | 95.38891004 | 63.08828103 | 8.203741329 | 1.552901863 | 72.36632316 | 20.1662964 | 96.06377881 | 65.30383371 |

**S11 Table.** *Ab initio* folding performance of DConStruct on EVfold dataset using top hybrid interaction maps with tri-level thresholding at increasing xL values (x = 2, 4, 8, 16).

| Target | 2L | 4L | 8L | 16L |
|---|---|---|---|---|
| 1bkrA | 0.8383 | 0.8267 | 0.8257 | 0.7377 |
| 1e6kA | 0.8332 | 0.8516 | 0.8972 | 0.784 |
| 1f21A | 0.7557 | 0.7753 | 0.8162 | 0.7386 |
| 1g2eA | 0.5946 | 0.7795 | 0.7998 | 0.6301 |
| 1hzxA | 0.6165 | 0.715 | 0.7326 | 0.6934 |
| 1oddA | 0.194 | 0.2153 | 0.2592 | 0.2625 |
| 1r9hA | 0.544 | 0.7663 | 0.7929 | 0.6209 |
| 1rqmA | 0.7086 | 0.7168 | 0.7427 | 0.6888 |
| 1wvnA | 0.719 | 0.807 | 0.8566 | 0.5849 |
| 2hdaA | 0.2272 | 0.2188 | 0.2816 | 0.1883 |
| 2it6A | 0.1248 | 0.2079 | 0.2617 | 0.2008 |
| 2o72A | 0.1724 | 0.1568 | 0.1749 | 0.1777 |
| 3tgiE | 0.729 | 0.7821 | 0.8513 | 0.8407 |
| 5p21A | 0.781 | 0.7736 | 0.8141 | 0.7656 |
| 5ptiA | 0.6135 | 0.7171 | 0.7072 | 0.5867 |
| Mean | 0.563453333 | 0.620653333 | 0.654246667 | 0.566713333 |
